# Eliminating Aggressive Cancers via PROTAC-like Inducers of Ferroptosis

**DOI:** 10.1101/2024.12.28.630572

**Authors:** Avital Oknin-Vaisman, Deepanjan Panda, Rostislav Novak, Eliya Bitman-Lotan, Nikolett Pahor, Yamen Abu Ahmed, Guy Kamnesky, Markus E. Diefenbacher, Ashraf Brik, Amir Orian

## Abstract

Aggressive and therapy-resistant cancers present a significant challenge to treatment and are associated with poor patients’ survival. Identifying molecular pathways and compounds that target these pathways is critical for improving patient outcomes. RNF4, an E3-ubiquitin ligase, is pivotal in oncoprotein stabilization and DNA repair, enhancing cancer cell survival driving tumorigenesis. Elevated RNF4 levels are associated with poor prognosis in cancer patients. Here, we describe the development of R4VPs, proteolysis-targeted chimeras-like (PROTACs-like). R4VPs promote RNF4 degradation and reduce the levels of its stabilized oncoproteins. Notably, R4VPs induce ferroptotic cell death selectively in cancer cells, sparing non-tumorigenic and primary cells. Surprisingly, R4VPs-induced ferroptosis is independent of RNF4 but preferentially targets tumor-driving mutations, particularly those in the EGFR pathway, while not affecting PI3K-transformed cells. R4VPs effectively induce cell death in therapy-resistant melanoma and sarcomas including patient-derived sarcoma tumor cells. Our findings highlight the potential of ferroptosis inducers such as R4VPs as a therapeutic strategy for therapy resistance, aggressive, and hard-to-treat cancers.

**Teaser:** R4VPs induce ferroptotic death of melanoma and sarcoma cells including patient-derived tumor cells sparing non-transomed cells.

## Introduction

Despite major advances in cancer therapies, the patient’s response to treatment for aggressive, advanced, and/or therapy-resistant tumors remains a challenge. Cancer development and progression are intimately linked to increased oncoprotein stabilization. Moreover, the development of resistance to chemotherapy and molecular treatments such as receptor tyrosine kinases inhibitors (RTKi), and/or immune check inhibitors (ICI) in melanoma, is also associated with increased oncoprotein stabilization (1–8). These hallmarks are observed in the majority of patients, and therefore overcoming therapy resistance is an unmet clinical need. The development of tailored precision therapeutics targeting aggressive and therapy-resistant tumors is required for improving patient outcome.

We recently discovered a novel ubiquitin-dependent pathway that stabilizes and potentiates oncoprotein activity promoting tumorigenesis. Oncoprotein stabilization requires their phosphorylation by mitogenic kinases regardless of the ubiquitin machinery or the degron’s that mediate the degradation of these otherwise short-lived oncoproteins (7, 9–11). The central enzyme in this pathway is the ubiquitin ligase RNF4, which ubiquitinates these oncoproteins generating heterotypic K11, K33 poly-ubiquitin chains leading to stabilization, and potentiating the transcriptional activity of multiple phosphorylated oncoproteins such as p-c-Myc, p-c-Jun, p-β-catenin and others (9). Moreover, in addition to its role in protein stabilization, RNF4 has multiple roles supporting cancer development. Among these functions are DNA damage repair, nuclear protein control, enhancement of oncogenic transcription and translation, and tumorigenic impact on the tumor microenvironment including fostering angiogenesis (Fig. 1A, B; 12-20).

**Figure 1:**
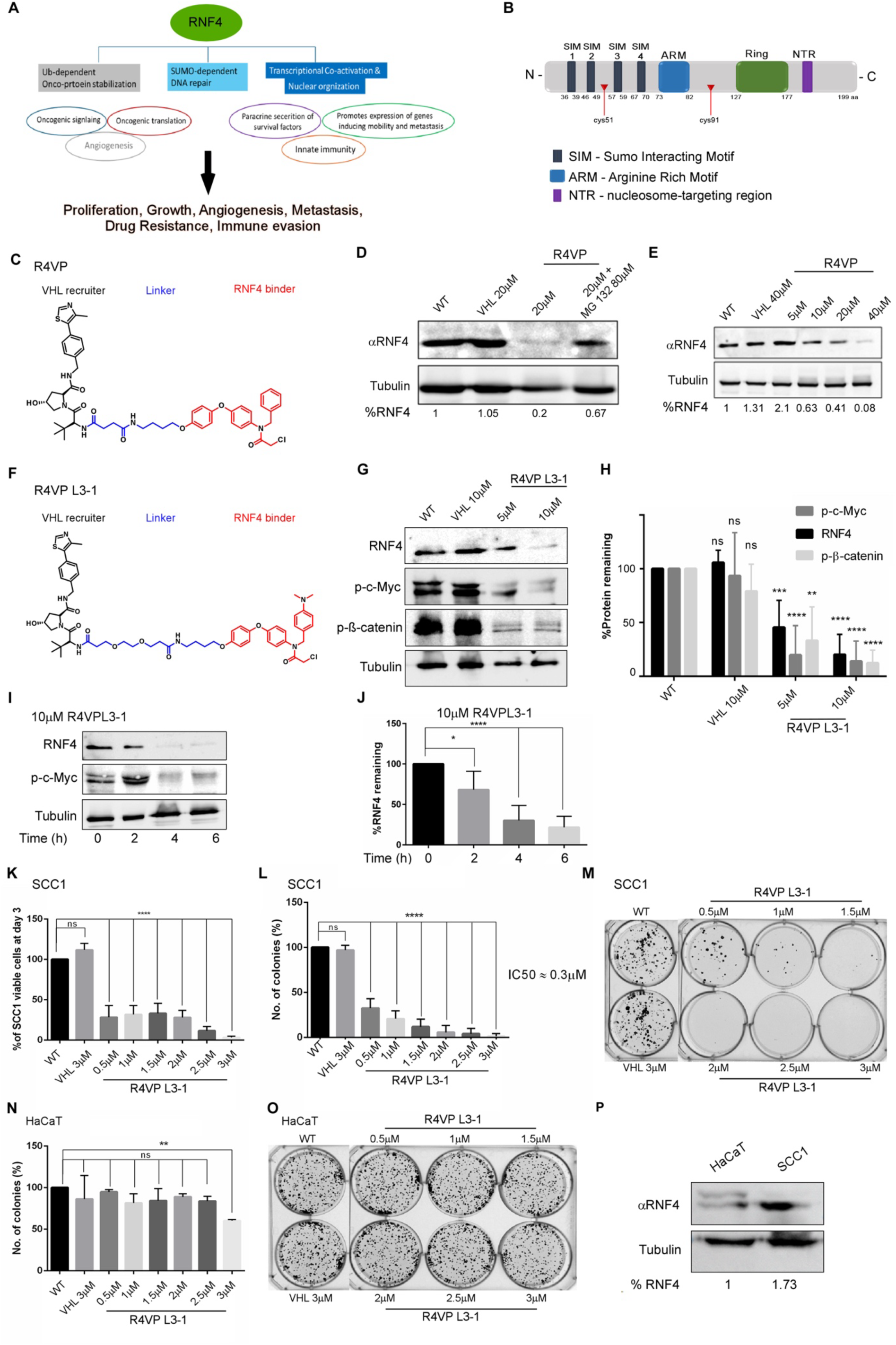
R4VP degraders force the proteasomal degradation of RNF4, its stabilized proteins, and reduce cancer cells survival. **(A)** Cancer-promoting activities of RNF4. **(B)** Schematic diagram of RNF4 structure, red triangles mark animo acids residues Cys51 and 91 that are required for R4VPs binding to RNF4. **(C)** Chemical structure of R4VP. **(D)** Western-blot analysis of endogenous RNF4 protein level in extracts derived from B16F10 mouse melanoma cells. VHL-only compound (VHL-r), or RV4P were added for three hours, and where indicated, proteasome inhibitor 80μM MG132 was added two hours prior to treatment with the compounds. Tubulin serves as a loading control. **(E)** R4VP treatment results in a dose-dependent reduction in RNF4 levels in B16F10 mouse melanoma cells. **(F)** Structure of R4VPL3-1. **(G-J)** Western-blot analysis of endogenous RNF4, p^Ser62^-c-Myc, and p^Ser45^-β-catenin proteins levels in extracts derived from A375R PLX4032-resistant human melanoma cells treated with VHL-r or R4VPL3-1 at the indicated concentrations (G, H), and time (I, J). Tubulin serves as a loading control, and in all experiments n=3 and ****= p<0.0001, ***= p<0.001, **=p<0.01. **(K-M)** Proliferation (K) and sphere formation (SFA; L, M) of SSC1 human squamous skin carcinoma cells is attenuated upon treatment with R4VPL3-1 at the indicated doses, but not upon treatment with VHL-r compound. (**N-O)** SFA of HaCat, a non-tumorigenic skin keratinocyte cell line is only minimally inhibited by R4VPL3L-1 Statistical analysis: Statistics: In all experiments, ****= p<0.0001, ***= p<0.001, **=p<0.01. ns= non-significance. (H) 2 -way Anova Dunnett’s multiple comparisons test. RNF4 (n=5), p-c-Myc (n=4), p-b-catenin (n=3) (J, K, L, N) 1-way Anova Dunnett’s multiple comparisons test. n=3 **(P)** Endogenous RNF4 protein levels in HaCat and SSC1 cells, Tubulin serves as a loading control.

RNF4, belongs to a small group of evolutionarily conserved RING ubiquitin E3 ligases termed SUMO-targeted ubiquitin ligases (STUbL) that ubiquitinate SUMOylated proteins (21–23). RNF4 has a tumor suppressive function in the case of acute pre-myelocytic leukemia (APL), in part by the SUMO-dependent ubiquitination and degradation of the leukemia oncoprotein driver PML-RARα (24–25). However, in a large group of tumors comprising carcinomas (breast and colo-rectal carcinomas, cholangiocarcinoma, and hepatocellular carcinoma), melanoma, sarcomas, and Myc-driven lymphomas, RNF4 has a pro-tumorigenic role (9–11, 26–28). We previously characterized the molecular and biochemical actions of RNF4 *in vitro*, *in cellulo*, and *in vivo*. We found that RNF4 enhanced the tumorigenic properties of cancer cells, being essential for cancer cell survival. In melanoma RNF4 confers resistance to molecular therapy *in vitro* and *in vivo* (9, 10). In sarcomas, RNF4 drives the expression of the survival factors; BMP6 and its co-receptor RGMb that are secreted from the tumor cells and act locally (11). High levels of RNF4 are observed in about ∼30-40% of colon cancer, melanoma and sarcoma biopsies, and are associated with poor prognosis in breast cancer, melanoma, and multiple types of sarcomas. While RNF4 is non-oncogenic on its own, cancer cells are “addicted” RNF4, in which eliminating genetically RNF4 resulted in the death of aggressive and therapy-resistant cancer cells and tumors (9–11). Thus, RNF4 is potentially an “Achilles’ heel” of multiple tumor entities, which makes it an excellent target for precise cancer therapy.

Inhibition of the enzymatic activity of RING proteins is challenging. Taking a different approach, we decided to eliminate RNF4 via targeted protein degradation (TDP), rather than inhibit its activity, by designing specific proteolysis targeting chimera molecules (PROTACS; 29-35). PROTACS are hetero-bifunctional molecules that are double-headed; on one-hand bind to the protein of interest (POI) and on the other hand bind to an E3-ligase recruiter, that are connected via a short linker. This proximity results in ubiquitination and proteasomal degradation of the POI. Several E3 recruiters have so far discovered and are in advanced clinical trials (34). Among these E3-recruiters are well-known Cerbelon (Crbn) and the Von-Hipple Lindau (VHL) ubiquitin ligase proteins (30, 36–38). In addition, RNF4 was suggested to be an E3 recruiter, that when linked to a PROTAC containing the small molecule JQ1 induced the degradation of the oncogenic BRD4 transcriptional co-factor (39). However, eliminating RNF4 in cancer using a PROTAC strategy has not been reported.

Here we report on the development of PROTAC-like R4VPs that promote the proteasomal degradation of RNF4 and reduces the protein level of its stabilized oncoproteins. Surprisingly, we found that R4VPs induces ferroptosis independently of RNF4, leading to the rapid death of human melanoma cells resistant to RTKi, and sarcoma cells, including patients-derived primary tumor cells, but have no effect on non-tumorigenic cell lines or primary MEFs.

## Results

### R4VP∼PROTAC-like mediated proteasomal degradation of RNF4

As a starting point, we tested whether eliminating RNF4 by a PROTAC-like molecule will be a suitable strategy to eradicate cancer cells. We used the previously reported RNF4 binding molecule CCW16, termed here L1 (39), and linked it to an established VHL recruiter moiety (VHL-r), thereby creating a PROTAC-like molecule that we termed R4VP (Fig.1C; for details for synthesis of all compounds and their validation see Chemical SI).

Next, we evaluated the steady-state protein levels of endogenous RNF4 in murine B16F10 melanoma cells upon treatment with either R4VP or VHL-recruiter (VHL-r) alone, respectively. Treatment with R4VP (3h), but not VHL-r, resulted in a dose-dependent reduction in endogenous RNF4 protein level, that was inhibited by the proteasome inhibitor MG132 (Figure 1D, E), supporting the proteasomal degradation of RNF4 in B16F10 cells via R4VP.

### Anti-cancer activities of R4VP

To test the anti-cancer activities of the degrader-like molecule R4VP towards multiple cancer and non-tumorigenic cells we treated a set of cell lines with increasing amounts of R4VP or VHL-r. We observed that treatment with R4VP, but not VHL-r, inhibited the proliferation and sphere formation (SFA) of SCC1 human squamous skin cancer cells, as well as human melanoma cell line (A375R) that is resistant to PLX4032, a Vemurafenib^®^ analog. Notably, R4VP did not affect the proliferation of non-tumorigenic human keratinocytes cell line HaCat. Moreover, an inactive R4VP lacking the reactive chloro-atom, that mediates the covalent reaction of R4VP with RNF4, showed no anti-proliferative activity (Fig. S1A).

To improve the activity of R4VP we prepared a focused compound-library, based on L1, for *in vitro* screening and identified L3 as an improved binder of RNF4 (Figs. S1B, Chemical SI). Since the structure of the linker is critical for PROTAC function (40), we also modify the original R4VP linker region and generated a series of R4VPs [R4VPL3-1, R4VPL3-2, and R4VPL3-3 (Figure 1F, Fig. S1C, Chemical SI]. While R4VPL3-2 was inactive, R4VPL3-1 and R4VPL3-3 exhibited similar potent anti-proliferative activities without affecting non-tumorigenic cells and MEFs (see below and Fig. S1C).

### Induced protein degradation and anti-cancer activity of R4VPL3-1

To further characterize the newly developed R4VPL3-1, we tested its impact on the protein level of endogenous RNF4 and the phosphorylated oncoproteins substrates that RNF4 stabilizes, such as p^Ser62^-c-MYC and p^Ser45^-β-Catenin (9). We observed a dose and time-dependent decrease in the levels of RNF4 and its stabilized substrates upon treatment with of R4VPL3-1, resulting in significant reduction in the level of RNF4, p-c-Myc, and p-β-catenin (Fig. 1F-J). Next, we tested R4VPL3-1 anti-cancer activity in SCC1cells and observed that R4VPL3-1 inhibited proliferation and SFA of this cell line with an IC_50_ of ∼0.3 μM (Fig. 1K-M). In contrast, at these concentrations, R4VPL3-1 showed only a minimal effect on non-tumorigenic HaCat cells, albeit the observation that both cell lines express RNF4 (Fig.1N-P).

### R4VPL3-1 induced rapid cell death of RTKi-resistant melanoma cells

One major clinical challenge is overcoming therapy resistance (2–4). RNF4 was shown to confer resistance to RTKi in melanoma cells such as A375R (10). Therefore, we tested whether R4VPL3-1 is able to overcome drug resistance of this cell line, A375R. We observed that R4VPL3-1 inhibited the proliferation and SFA of aggressive A375R under PLX4032 treatment (Fig. 2A-C). To gain further insights regarding the inhibitory effects of R4VPL3-1 on cell viability and SFA of A375R cells we performed FACS analysis using Annexin V and Propidium Iodide (PI) staining. Treatment with R4VPL3-1 induced rapid cell death of A375R cells resulting in >50% cell death in a time and dose-dependent manner (Fig.2D-G).

**Figure 2:**
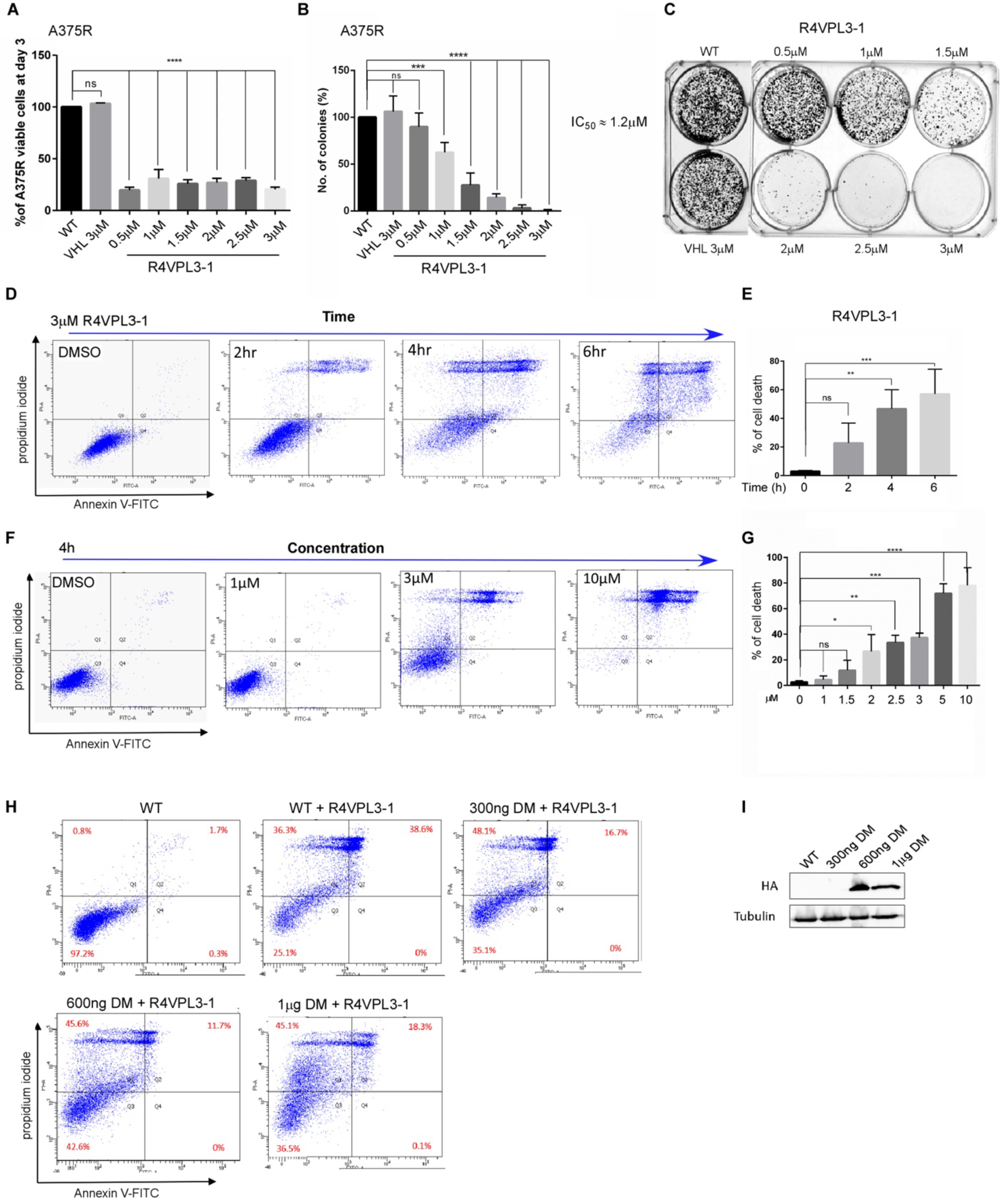
R4VPL3-1 induces human A375 RTKi-resistant melanoma cells death. **(A-C)** Proliferation (A) and SFA (B, C) of human PLX4032-resistant 375R cells is attenuated upon treatment with R4VPL3-1 at the indicated doses, but not upon treatment with VHL-r compound, and (C) is a representative experiment. **(D-G)** Cell death determined using FACS analysis using propidium iodide and Annexin V-FITC. Treatment of A375R cells with R4VPL3-1 results in rapid cell death. (D, E) Time-course of cell death upon treatment with 3μM R4VP3-L1. (F, G) Dose-dependent four hours treatment with R4VP3-L1, (D, F) are representative experiments. **(H)** Expression of increasing amounts of HA-RNF4^DM^ (RNF4^C51A,C91A^) mutant does not cancel R4VPL3-1∼induced cell death. **(I)** HA-RNF4^DM^ protein levels in the experiments performed in (H), Tubulin serves as a loading control. Statistical analysis was performed using 1-way Anova Dunnett’s multiple comparisons test (A, n=2; E n=4 ;G n=3).

While protein degraders are designed to induce the degradation of a specific POI, the potential binding to other proteins is well-known (41). To address the question of specificity, we mapped the binding site(s) of R4VPL3-1 to RNF4 using a biotin-tagged R4VPL3-1, where the VHL-r was replaced with biotin (biotin-R4VPL3-1). Mass spectrometry and *in vitro* binding analyses identified that biotin-R4VPL3-1 binds to Cys51 and 91 within RNF4 (Gotthardt & Müller, personal communication). Subsequently, we tested whether overexpression of RNF4 double mutant HA-RNF4^C51A, C91A^ termed RNF4^DM^, where the Cys acceptor a.a. residues were replaced with Ala, and that does not bind to R4VPs, may protects A375R cells from R4VPL3-1-induced cell death. We found that increasing doses of HA-RNF4^DM^ did not prevent R4VPL3-1-induced cell death (Fig. 2H, I). Thus, while R4VPL3-1 binds to RNF4 and reduces RNF4 levels, along with a decrease in the levels of its stabilized oncogenic substrates, the cell death induced by R4VPL3-1 cannot be majorly attributed to RNF4.

### R4VPL3-1 induces ferroptosis in cancer cells

To identify the cellular pathways involved in R4VPL3-1-induced cell death and anti-cancer activities we performed transcriptional analysis using whole transcriptomic sequencing and compared A375R cells treated for two hours by VHL-r or R4VPL3-1 (Fig. 3A-C, Fig. S3). We identified 316 differentially regulated genes (DEGs) specific to R4VPL3-1 treatment. Of these, 191 genes were upregulated and 125 downregulated upon treatment with R4VPL3-1 (Table S1). Subsequent pathway analysis using Kyoto Encyclopedia of Genes and Genomes (KEGG) analysis identified ferroptosis and necroptosis as predominant pathways transcriptionally regulated by R4VPL3-1.

**Figure 3:**
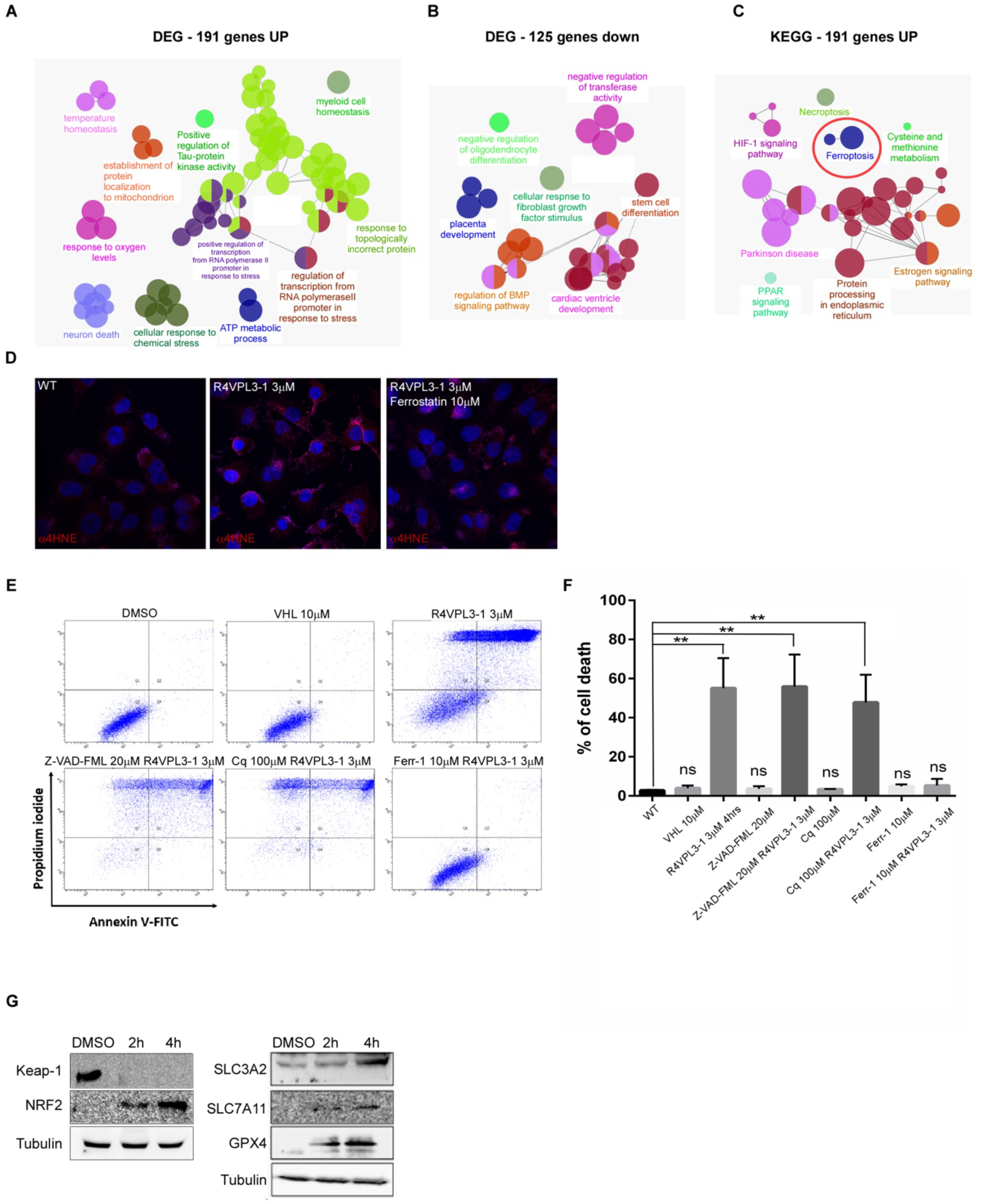
R4VPL3-1 induces ferroptosis of A375R RTKi resistant melanoma cells. **(A-C)** Cytoscape and KEGG analysis of RNA-seq results identifying statistically significant upregulated and repressed pathways, as well as cellular process effected by two hours treatment 10μM of R4VPL3-1 compared to VHL-r treated A375R cells. In (C) red circle marks ferroptosis and full gene lists are in Table S1. **(D)** 5μM R4VPL3-1 induces lipid peroxidation that is inhibited by 10μM a Ferrostatin-1 (Ferr-1), a specific ferroptosis inhibitor, as evident by α-4HNE immune-staining. **(E, F**) FACS analysis of R4VPL3-1 induced cell death. 10μM Ferr-1 but not 20μM Z-VAD-FMK or 100μM Chloroquine (Cq) inhibits R4VPL3-1 induced cell death. (F) is a representative experiment. Statistical analysis: n=2 and ****= p<0.0001, ***= p<0.001, **=p<0.0, ns= no-significance. 1way Anova Dunnett’s multiple comparisons test (n=2). **(G)** Westen blot analysis of protein extracts derived from A374R cells treated with R4VPL3-1 for the indicated times resulted in reduction in Keap1 protein levels and an increase in the levels of NRF2, SLC3A2, SLC7A11 and GPX4 proteins.

Ferroptosis is a form of regulated cell death characterized by iron-dependent lipid peroxidation and increase in free cellular iron (42–46). To determine whether R4VPL3-1 induces lipid peroxidation, a hallmark of ferroptosis, we treated A375R cells with 5μM of R4VPL3-1, or R4VPL3-1 along with the ferroptosis inhibitor 10μM Ferrostatin-1 (Ferr-1). We used established markers for lipid peroxidation; A major product that is formed by lipid peroxidation is 4-Hydroxynonenal (4HNE), and we used α-4HNE antibody to monitor lipid-peroxidation. We identified that R4VPL3-1 induced lipid peroxidation, as evident by positive 4HNE immuno-staining, suggesting that the R4VPL3-1 indeed induced ferroptosis. This finding is further supported by the observation that 4-HNE formation was inhibited by pre-treatment with 10μM Ferr-1 (Fig. 3D; 43). Collectively, these results suggest that R4VP3L-1 induces ferroptotic cell death.

To test whether additional pathways may also be involved in the cell death induced by R4VPL3-1 we pre-treated tumor cells with either alone, or pre-incubated with Z-VAD-FMK (pan-caspase inhibitor), chloroquine (lysosomal inhibitor) or Ferr-1, and subsequently add R4VPL3-1. While Z-VAD and chloroquine did not inhibit R4VPL3-1-induced cell death, Ferr-1 completely prevented R4VPL3-1-induced cell death (Fig. 3E, F). The significant ferroptotic stress imposed by R4VPL3-1 concomitantly resulted in a failed counter-activation of ferroptosis inhibiting genes, both at the transcriptional and protein level, such as the nuclear factor erythroid-related factor 2 (NRF2/NFE2L2), a major anti-ferroptotic transcription factor (47). Furthermore, we observed a reduction in protein levels of the Kelch domain E3 ligase Keap1, which is essential for the ubiquitin-dependent degradation of NRF2 (Fig. 3G; 48-50). In addition, we observe an accumulation of metabolic enzymes known to reduce oxidative stress and that are NRF2 targets-genes and inhibit ferroptosis such as GPX4 and Cys transporters SLC3A2, SLC7A11(Fig. 3G, Supp table 1; 43, 51-54).

### Molecular determinant for R4VPL3-1 selectivity

We were intrigued by the differential effects of R4VPs that induced cell death of cancer cells, but not affecting non-tumorigenic HaCat cells nor primary MEFs. Therefore, we tested for inhibition of SFA upon treatment with the different R4VPL3-1 components, as well R4VPL3-1 where we replaced the VHL-r moiety with Biotin (Biotin-R4VPL3-1, Fig. 4A). We observed that the RNF4 binding moiety, R4B inhibited SFA of both non-tumorigenic HaCat and primary MEFs as well as cancer cells. In contrast, R4VPL3-1 had potent activity towards cancer cells but had no activity against non-tumorigenic cells (Fig.4B-E). Moreover, Biotin-R4VPL3-1 was an inactive compound that had no activity against cancer cells and non-tumorigenic cells (Fig. 4F-I).

**Figure 4:**
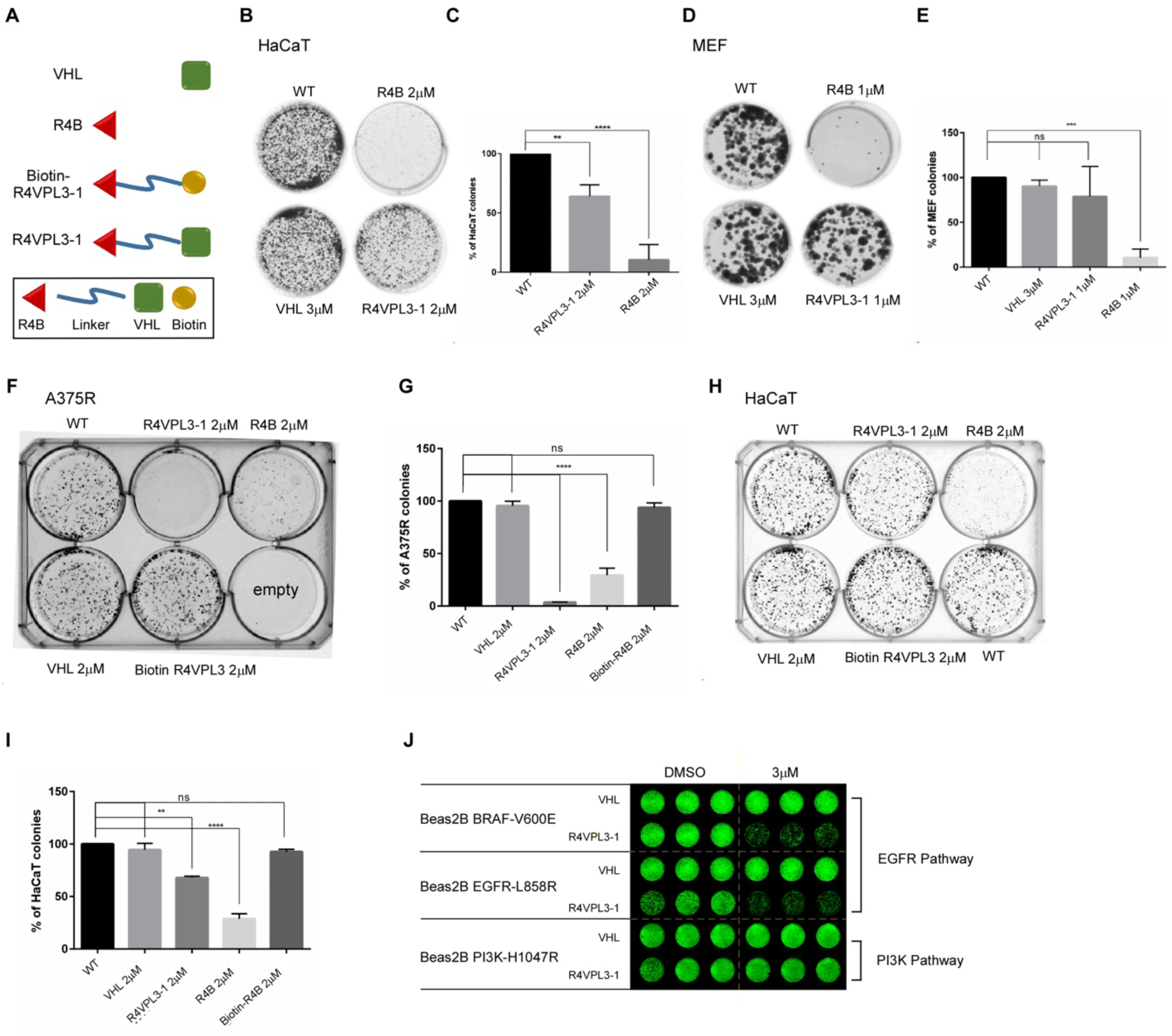
Structural determinants of R4VPL3-1 involved in its differential anti-cancer activity. **(A)** Schematic diagram of R4VP-related compounds; **(B-E)** Activity of R4VP-related compounds toward non-tumorigenic HaCaT cells (B, C), and MEFs (D, E); (B) and (D) are representative experiments. **(F-I)** SFA; R4VPL3-1 where the VHL moiety was replaced with biotin (Biotin-R4VPL3-1) has no anti-cancer activity towards A375R (F, G) or HaCat (H, I). **(J)** Live Cell proliferation assays using α-LamA/C indicated transformed epithelial lung cells (see methods). R4VPL3-1 has selective anti-proliferative activity against Beas2B lung cell transformed with either BRAF^V600C^ or EGFR^L858R^ activating mutations, but not PI3K^H1047R^ mutation. Statistical analysis: 1way Anova Dunnett’s multiple comparisons test (C, n=3; E, n=3; G, n=2; I, n=2); Significance: ****= p<0.0001, ***= p<0.001, **=p<0.0, ns= no-significance.

Interestingly, we found that R4VPL3-1 has differential activity towards cancer cells depending on the oncogenic driving mutation used to induce cell transformation. In this experiment we used a human tracheal epithelial cell line BEAS-2B, that was retrovirally transformed with activated oncogenes: BRAF^V600A^, or EGFR^L858R^,or PI3K^H1017R^ (55). R4VPL3-1 inhibited the proliferation of BRAF^V600A^, EGFR^L858R^, transformed lung cancer cells, but did not inhibit proliferation of PI3K^H1017R^ transformed cells (Fig 5J). This is likely due to ferroptosis resistance induced by PI3K∼mTor signaling (56; see discussion). Taken together, this set of experiments suggests that the activity of R4VPL3-1 towards aggressive cancer cells requires the VHL recruiting moiety and is cancer-pathway specific.

**Figure 5:**
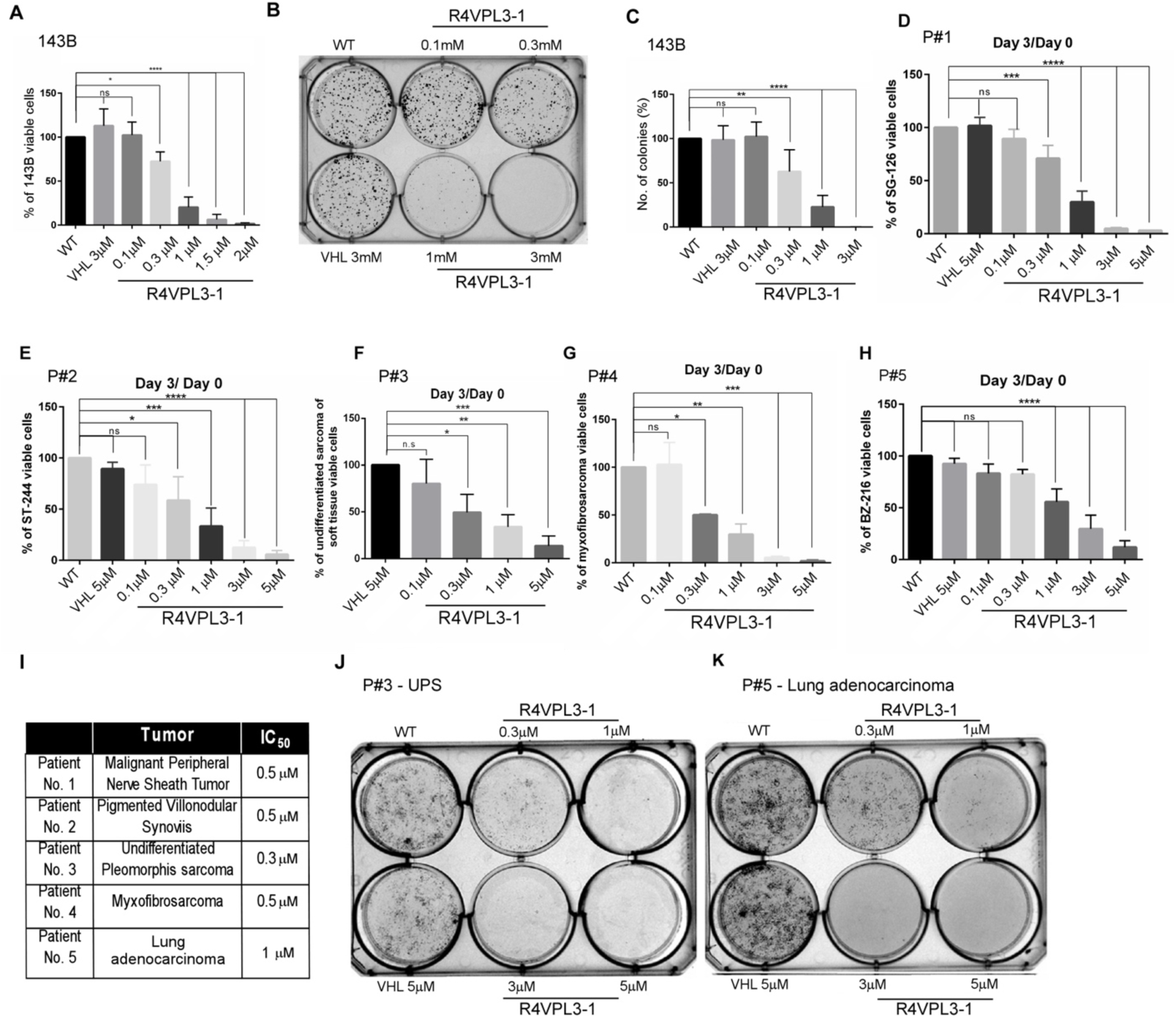
R4VPL3-1 is active towards sarcoma cells, patients-derived primary tumor cells. **(A-C)** R4VL3-1 inhibits cell proliferation (A) and SFA (B, C) of 143B, a highly metastatic human sarcoma cell line. **(D-I)** R4VL3-1but not the VHL-r inhibits cell proliferation of patients-derived primary tumor cells in a dose dependent manner. (I) Table summarizing IC_50_ of R4VPL3-1 in different patients-derived cells. Patients #1-4 are sarcoma tumor cells and patient #5 is a lung adenocarcinoma tumor. **(J, K)** R4VPL3-1 inhibits SFA of undifferentiated polymorphic sarcoma (UPS) (J), and lung adenocarcinoma tumor cells (K). Statistical analysis: 1way Anova Dunnett’s multiple comparisons test (A, n=3; C, n=4; D, n=4; E, n=3; F, n=3, G, n=2; H, n=3); Significance.****= p<0.0001, ***= p<0.001, **=p<0.01. ns=non-significant.

### Pre-clinical testing of R4VPL3-1 in patient-derived primary sarcomas

One unmet clinical need is the treatment for sarcomas (57–59). We previously observed that RNF4 is essential for sarcoma tumorigenicity, and high levels of RNF4 are observed in multiple sarcoma entities and are associated with poor sarcoma patient survival (11). While other proteins may mediate R4VPs anti-cancer activity we decided to test the impact of R4VPs on the metastatic human sarcoma cell line 143B. R4VP treatment of these cells resulted in the inhibition of proliferation and SFA (Fig 5A-C). Furthermore, R4VPL3-1 was potent towards primary untreated patient-derived sarcoma and lung cells isolated during onco-surgical resections, inhibiting proliferation and sphere formation in culture with IC_50_ ∼0.5μM of different sarcoma entities (Fig. 5D-K). Thus, R4VPL3-1 is highly active against sarcoma cells including diverse primary patients-derived sarcoma tumor cells.

## Discussion

We develop an ferroptosis-inducing PROTAC-like molecules, R4VPs, which eliminated multiple cancer cells, including RTKi-resistant human melanoma, primary patient-derived sarcoma, and metastatic lung cancer cells depending on the tumor driving mutation. In contrast, non-tumorigenic cell lines and primary MEFs are not affected by R4VPs. R4VPs induce the proteasomal degradation of RNF4 and its stabilized p-oncoproteins. However, R4VPs-induced ferroptosis is independent of RNF4, a cell death that was inhibited by the ferroptosis inhibitor Ferr-1.

We used CCW16, an RNF4 binding moiety, as a basis for a focused screen and discovered an RNF4 binder with higher affinity to RNF4 (R4B) and used this molecule to generate more potent R4VPs. Previous mass spectrometry analysis suggested that TRH1-23, an early version of CCW16, reacts with Cys132 and Cys135 residues within the RING domain of RNF4, and that CCW16 was shown to potentially react with Cys135 in a computational covalent docking prediction (39). However, mutating these Cys residues within the RING domain, that were reported to be required for binding to CCW16 and its derivatives (39),reduced but did not abolish its interaction with biotin-R4VPL3-1.

R4B, the RNF4 binding moiety, induced cell death indiscriminately between tumors and non-tumors cells. RNF4 was suggested to serve as an E3 mediating proteasomal degradation of POIs in Targeted protein degradation (TPD; 30). Thus, our observations suggest that a careful examination should be given to potential harmful effects of CCW16-like molecules. R4VPs activity is cell-type specific and induced the proteasomal degradation of RNF4 in B16F10 melanoma cells. Furthermore, it reduced the level of RNF4 and its stabilized proteins in A375R human melanoma cells. However, R4VPs did not affect the levels of RNF4 in HeLa or HEK293 cells. In addition, we observed that the VHL-r moiety was essential for R4VPL3-1 anti-cancer activities, as its replacement with biotin, or with the well-established recruiter moiety that recruits the E3 Crbn (30), resulted in inactive compounds. Thus, the VHL recruiting moiety may be required for the degradation other suppressors of ferroptotic cell death POIs. Moreover, our attempt to inhibit the anti-cancer activities of R4VPs by inhibiting the ubiquitin pathway using the E1 inhibitor TAK-243 was not successful, likely due to significant rapid cell-toxicity upon co-treatment. Considering these limitations, we termed R4VPs PRTOAC-like molecule.

While RV4Ps efficiently induced the proteasomal degradation of RNF4, this case is not trivial, as R4VPs bring into proximity two E3 ubiquitin ligases, and the outcome of this interaction is empiric and even may be context dependent. In our study the VHL E3 was dominant, resulting in RNF4 degradation. This is similar to the case of a VHL-based PROTAC targeting the oncogenic E3 ubiquitin ligase MDM2 in triple-negative breast cancer (60).

Defining target specificity and selectivity are central questions in developing small molecule inhibitors and protein-degraders. While we have shown that R4VP treatment leads to the degradation of RNF4, expression of RNF^DM^, that does not bind to R4VPL3-1, did not prevent R4VPL3-1-induced ferroptosis. Due to technical difficulties, and albeit extensive efforts, we were not able to perform the optimal experiment, in which the endogenous RNF4 is eliminated (by CRISPER/CAS9 gene editing or by 3’ UTR directed shRNA) and replaced by the co-expression of RNF4^DM^ in A375R. As RNF4 functions as a dimer (61), we therefore cannot formally rule out the possibility that the RNF4^DM^ expressed in these cells dimerizes with wild-type RNF4 that binds to R4VP leading to the degradation of the RNF4/RNF4^DM^ heterodimer.

However, an alternative explanation is that R4VPs have additional substrate(s) that their elimination induces ferroptosis and cell death. It is interesting to note that CCW28-3, a BRD-4 degrader based on CCW16 was shown in a TMT-based proteomic quantification to affect the level of multiple proteins likely because of its reactive chloroacetamide moiety, including proteins involved in ferroptosis (39). Thus, suggesting that in addition to RNF4, R4VPL3-1 may induce the degradation of ferroptosis inhibitors in ferroptosis-sensitive cancer cells.

Ferroptosis is a cell death pathway with distinct molecular and histopathological features (42). It primarily involves lipid-peroxidation and accumulation of free iron, leading to an increased oxidative stress, ending with rupture of the cytoplasmic membrane. Ferroptosis is intrinsically inhibited by NRF2, the key antioxidant transcription factor, that its degradation involves by Keap1 and also RNF4 (48–50). It is also inhibited by multiple enzymes and metabolic pathways such as Cys and Cys/Glu transporters that increase activity of anti-oxidative enzymes, reduce free iron, as well as enzymes that minimize lipid-peroxidation such as in Glutathion Peroxidase 4 (GPX4, 42,51, 53, 54). Indeed, the strong ferroptotic induction by R4VPL3-1 induced a transcriptional and proteomic response that upregulated anti-ferroptotic proteins similar to reported studies (62).

One hallmark of cancer cells is their ability to evade cell death by multiple mechanisms including the development of resistance to therapy (1, 63). The sensitivity to ferroptosis varies among cancer cells and is context dependent (42). In this regard, we observed that even within the same cell of origin, the sensitivity to R4VPL3-1 was greatly dependent on the identity of the cancer driving mutation used for cell transformation. Epithelial cells transformed with activated oncogenes within the EGFR pathway were highly sensitive to R4VPL3-1. However, parental cells transformed with PI3K^H1017R^, a PI3K mutation that activate the oncogenic PI3K pathway, were resistant to R4VPL3-1. Indeed, PI3K signaling via the mechanistic target of rapamycin (mTOR) pathway is known to suppress ferroptosis by increasing the processing and activity of the ER-anchored transcription factor sterol-regulatory element binding protein (SREBP). In turn, SREBP induces the expression of stearoyl-CoA desaturase (SCD1) that enhances the production of ferroptosis-suppressing monounsaturated fatty acid (MUFA; 64). Moreover, R4VPs differential activity may be dependent on the presence of R4VPs substrate(s) that inhibit ferroptosis, and that these cancer cells are addicted too. Treatment with R4VPs likely results in their degradation and a shift in the balance between anti- and pro-ferroptotic cell states. Thus, as R4VPs anti-cancer activity stems from inducing ferroptosis, ferroptosis-resistant cells will be likely also resistant to R4VPs.

A growing amount of evidence suggest that inducing ferroptosis may be a potent strategy to induce cancer cell death including in therapy and aggressive cancers (46, 65). Ferroptosis can be induced genetically by expression of ferroptotic inducer genes such as NCOA4, miR-129p, miR-672-3p, or by small molecules collectively termed ferroptosis inducers [FINs; for a comprehensive review regarding FINs and FS see (66)]. Moreover, ferroptosis can be induced by a PROTAC inducing the degradation of GPX4 (67). However, the case of R4VPs induce ferroptosis is different as it takes place in cancer cells albeit high level of GPX4.

Our study suggest that R4VPs are bona-fide FINs that induce ferroptosis but not apoptosis and may have selectivity to specific cancer cells such as RTKi-resistant melanoma, and multiple sarcoma entities. However, the full spectrum of R4VPL3-1 sensitive and resistant cancer entities awaits future studies. Moreover, our observation that R4VPL3-1 has relatively long half-life (>12h) in the plasma of mice *in vivo* suggest that R4VPL3-1 may be potentially tested in an *in vivo* setting and more broadly that R4VPs potentially can be used as experimental FINs *in cellulo* and *in vivo*.

Sarcoma of different types including bone and soft tissue sarcomas present a clinical challenge. Albeit extensive efforts, no major improvement in patients’ survival and more specifically in advanced metastatic disease (58, 59, 68). Thus, our observation that R4VPL3-1 is effective in eliminating multiple patient-derived sarcoma entities is encouraging. Thus, testing the impact of R4VPs *in vivo* in a sarcoma xenograft model and patients-derived tumor xenograft will be a logical immediate step. Finally, our observation that R4VPs are potent ferroptotic inducer strongly suggests that compounds inducing ferroptosis may be a powerful strategy for the elimination of sarcomas and aggressive cancers.

## Materials and Methods

### Chemical synthesis and compound validation

Detailed of all chemical procedures, chemical structures and NMR validation are described under chemical supporting SI.

### Antibodies

αRNF4 antibody (1:200, sc-517643) and mouse α-tubulin (1:2000, SC-5286) were obtained from Santa Cruz Biotechnology. α-HA antibody was a kind gift from Ami Aronheim. p-c-Myc (1:500, #94015), p-β-catenin (1:300, #9564) and Ferroptosis Antibody Sampler Kit (#29650) were obtained from Cell Signaling Technology. α-4HNE was from Abcam (1:200, #ab46545). Lamin A/C primary labelled antibody (# Santa Cruz Biotechnology)

### Biological materials and compounds

Bacterially purified RNF4 was purchased from Boston Biochem (#E3-210), Image-iT™ Lipid Peroxidation Kit from Thermo Fisher Scientific (#C10445), Annexin from ENCO (#BLG-640906), Annexin V Binding Buffer was purchased from ENCO (#422201), Propidium iodide was purchased from Sigma-Aldrich (#P4170-10MG); 5-TMRIA [Tetramethylrhodamine-5-iodoacetamide] was from Megapharm (#AS-81410), BRAF inhibitor PLX4032^®^ was purchased from Selleckchem (#S1267). Ubiquitin E1 inhibitor TAK-243 was from MedChem express. uMLN-7243 aMG132 a Proteasome inhibitor was purchased from Mercury (#MBS474790, phosphatase inhibitor (#4906845001) and protease inhibitor (#11873580001) cocktails were from Sigma-Aldrich.

### Plasmids used in this study

pcDNA3 HA-hRNF4 was as in Thomas et al. (ref). pcDNA3 RNF4^C51A^, pcDNA3 RNF4^C91A^, RNF4^C51,91A^ were generated by quick-change site-directed mutagenesis (SDM^®^), according to the kit guidelines, using the relevant primers and validated by sequencing.

Primers: Primers used in this study are listed below:

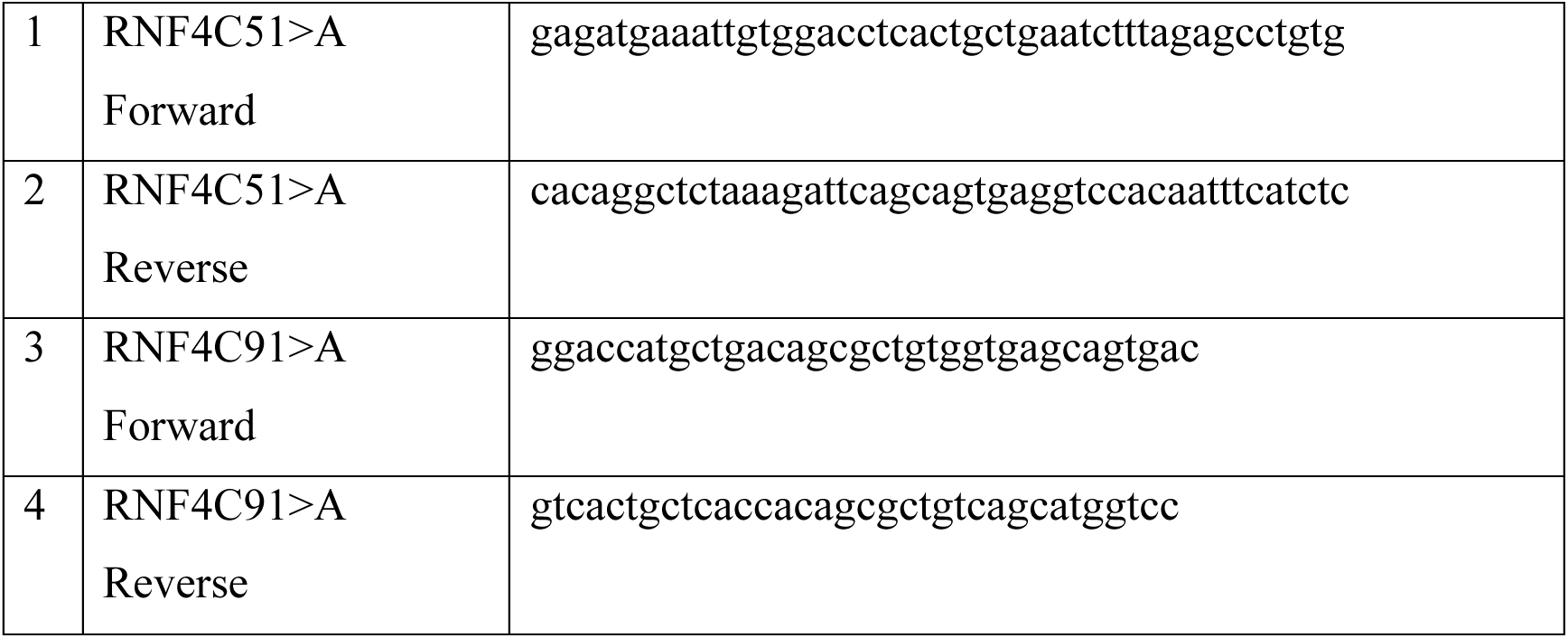

## Methods and experimental design

### Gel-Based 5-TAMRA-Iodoacetamide (IA) displacement assay

The binding of bacterially purified RNF4 to the indicated RNF4 binding moieties (R4Bs) was determined using 5-TMRIA displacement assay similar to (39); 200ng bacterially purified human RNF4 was incubated with the R4B molecules at the indicated concentrations in PBS solution. Reactions were incubated for 30 min at RT. Subsequently, 250nM of the 5-TMRIA-Tetramethylrhodamine-5-iodoacetamide,(dissolved DMSO) was added and incubated for 1 hour incubation at RT with a total reaction volume of 40μl. Binding was terminated by the addition of 8μL of 5× Lemli Sample buffer and heated at 90°C for 5 min. The samples were resolved over a 12.5% SD-PAGE. Fluorescent imaging was performed on LAS4000 (Image Quant) and quantified by ImageJ.

### Cultured cell line transfections and infections

A375R, HEK293, were as reported in (10). MEFs were a kind git of Yuval Shaked. All other cell lines were obtained from ATCC. Beas2B, HaCat, SCC1, and 143B were maintained in DMEM with 100U/ml penicillin, 0.1mg/ml streptomycin, Glutamine 4mM, and 10% FBS. A375R was cultured in DMEM with 10% FCS, penicillin/streptomycin, and 2µM PLX4032. The BEAS2B oncogenic cell line, reflects early oncogenic transformation stage of lung cancer (BEAS-2B EGFR^L8585^, BEAS-2B^BRAF-V600E^, BEAS-2BPIK3^H1047R^ were as previously described (55). All cell lines were authenticated by STR profiling. Cells were transfected using CalFectin transfection reagent (Sinagen Laboratories) according to manufacturer instructions.

### Cell Viability

Indirect cell viability was determined using CellTiter-Glo® Luminescent Cell Viability Assay (ATP-Lite; Promega). In brief, 10^3^ cells were seeded in triplicates for each time-point, in white 96-well plates with transparent bottom and cultured for the indicated time. 30 microliters CellTiter-Glo® solution were added to each well and viability was quantified by monitoring luminesce using a Plate-Reader according to the kit protocol.

### Sphare formation assay

A375R, SCC1 143B cancer cells and MEF, and HaCaT cells (1000 cells/well) were seeded in 6-well plates in 2ml DMEM medium, maintained at 37°C in humidified incubator. After 8 days, cells were gently once washed with PBS and fixed for 1 hour with 5% formaldehyde, washed thrice with PBS, stained with 0.05% crystal violet for 20 min, photographed and counted.

### Live cell proliferation

2000 BEAS2B transformed cells expressing the indicated oncogenic driver were seeded in 96-well plates. The following day, cells were treated with R4VPL3-1 or DSMO (solvent control) and were incubated for 72 hr. Subsequently, cells were fixed using ice-cold 100% Methanol (Sigma-Aldrich) for 10min on ice, washed three times with PBS for 5 min each, and blocked with 5% BSA in PBS, for 1 hour at RT. Afterwards, cells were incubated with Lamin A/C primary labelled antibody (Santa Cruz Biotechnology) in the dark for 1 hour at RT, followed by three washes with PBS. Imaging was carried out using the LI-COR Odyssey M Imaging System, which detects the intracellular fluorescent signaling in whole cells.

### Preparation of cell extract and determination of protein levels

Cells were washed three times with ice-cold PBS and harvested in RIPA supplemented with 10mM EDTA pH8.0 buffer along with protease and phosphatase inhibitors (Sigma-Aldrich). Cell extracts were sonicated using a microtip for 10” followed by centrifuge for 20 minutes at 13,000rpm. Extracts were resolved over SDS-PAGE, proteins were identified using the indicated antibodies and visualized and quantified using chemiluminescence LAS4000 and Image-Quant software. Fold changes in protein levels were calculated relative to Tubulin.

### Cell death analysis using flow cytometry

Cell death was determined by FACS using Annexin V-fluorescein isothiocyanate (FITC)/PI using flow cytometer. A375R were trypsinized, centrifuged, and washed with cold PBS. Subsequently, cells were resuspended in 500μL Annexin binding buffer, mixed with 5μL Annexin V + FITC and 20μL PI, followed by incubation at RT for 15 min in dark. Cells were analyzed using a flow cytometer following the manufacturer’s protocol. Annexin V and/or Propidium Iodide-labeled cell population were counted by flow cytometer.

### Determination of lipid peroxidation

10^5^ A375R cells were grown on coverslips in a 24-wells plate were treated with 3μM R4VPL3-1 as indicated for two hours. Where indicated 10μM Ferr1 were added two hours prior to RV4Ps treatment. Cells were washed with DMEM, fixed with 4% paraformaldehyde (PFA) for 15 minutes at room temperature on a on rotator. After fixation, the PFA was aspirated, and the cells were washed three times with PBS. Blocking was performed using a solution of 10% normal goat serum (NGS) and 0.3% Triton X-100 in 1× PBS for two hours at room temperature on a shaker to reduce nonspecific antibody binding. Primary antibody, α-4-HNE, a marker for lipid peroxidation, was diluted in a solution containing 5% NGS and 0.1% Triton X-100 in 1× PBS at a final concentration of 1:200. Cells were incubated with the primary antibody solution overnight at 4°C on a shaker, The following day, coverslips were washed five times with PBS for 10 minutes each on a shaker. Secondary anti-Rabbit antibody was diluted 1:1000 in PBS, with the inclusion of 4’,6-diamidino-2-phenylindole (DAPI) for mark nuclei. Coverslips were incubated with the secondary antibody for 1 hour at room temperature on a rotator. Post-incubation, the coverslips were washed three times with PBS for 10 minutes. Cells were mounted with mounting medium, and coverslips were placed on slides. Fluorescence images were acquired using a Zeiss LSM700 confocal microscope and magnification shown is X40.

### Establishment patients-derived sarcoma cells

Patients-derived sarcoma cells were resected from tumors during onco-orthopedic operation of non-treated patients, and according to Helsinki committee permit RBM-0536-22. Tumor specimens were collected in 50mL sterile tube with 20mL DMEM medium. Samples were washed with PBSx2, minced into minimal possible small fragments. In parallel part of the surgical specimens were analyzed by an experienced sarcoma pathologist and the exact nature of tumor was re-confirmed. Tumor slices were seeded on a 10cm dish and re-sliced and were seeded in 10cm plate. Attached cells were maintained in DMEM supplemented with 20% FBS, 1% glutamine, 1% Penicillin/Streptomycin and passaged once at 85-90% confluence by washing with PBS and seeded for experimentation at a split ratio not exceeding 1:3. 10^3^ cells were used for proliferation and SFA assays.

## RNA-seq and data analyses

RNA-Seq was performed in a set of three conditions using A375R cells and three independent biological repeats ; We compared differentially expressed genes (DEGs) between control treated A375R (DMSO) to cells treated with 10µM for 2h with R4VPL3-1 (the full PROTAC-like molecule) or with VHL-r. RNA extraction was performed using MACHEREY-NAGEL NucleoSpin RNA, Mini kit for RNA purification. RNA quality was evaluated using the TapeStation 4200 (Agilent) with the RNA kit (# 5067-5576). The RINe values of all samples were in the range of 9.3-10, indicating high quality. Libraries were constructed simultaneously using the NEBNext Ultra II Directional RNA Library Prep Kit for Illumina (NEB, #E7760), according to the manufacturer protocol. 800ng total RNA was used as starting material. mRNA pull-down was performed using the NEBNext® Poly(A) mRNA Magnetic Isolation Module (NEB, #E7490). RNA-seq library QC was performed by measuring library concentration using Qubit (Invitrogen) with the dsDNA HS Assay Kit (Invitrogen, #Q32854) and size determination using the TapeStation 4200 with the High Sensitivity D1000 kit (# 5067-5584). All libraries were then mixed into a single tube with equal molarity. The RNA-seq data was generated on Illumina NextSeq2000, using P2 100 cycles (Read1-100; Index1-8; Index2-8) (Illumina, #20046811). Single reads (100 bps) were aligned to the Homo sapiens (GRCh38.109) reference genome using STAR (V2.5.3a) with mismatch ratio allowed < 0.2, the minimum and maximum intron sizes were set to 20 and 1,000,000, respectively. The number of reads per gene was counted using Htseq-count (v2.0.2) with ‘reverse’ mode. Normalization and differential expression analyses were conducted using DESeq2 R package (v1.36.0). The similarity between samples was evaluated within DESeq2 package using a Euclidean distance matrix (shown in a heatmap with a clustering dendrogram) and a principal component analysis (PCA). The latter is drawn from the most variable genes (50, 500, 1000, 2000, 5000, 10000 genes). The threshold for significantly differentially expressed genes is determined by two factors: FDR adjusted p-value ≤ 0.05 (for mouse B16 experiment a p-value ≤ 0.01 threshold was specifically chosen since the DE analysis yielded over 4000 differential genes) and the ‘base-mean independent filtering’ threshold, which is calculated by the DESeq2 algorithm for each comparison. FDR was calculated using the default approach of DESeq2, Benjamini-Hochberg (BH). Since in both experiments, no major batch effect was indicated, a simple single-factor model was applied and lists of DEGs were created for all the possible contrasts.

### Data analysis and Bioinformatics tools

For RNA-seq analysis Gene ontology analyses was performed using Cytoscape software including: ClueGO app (v2.5.10) in Cytoscape (v 3.10.1) was used to conduct GO enrichment analyses. In our study, ClueGO we used to identify different functional groups in the following terms: 1. Biological Process (BP) A p-value ≤ 0.001 was used as the cut-off criterion; 2. KEGG A p-value ≤ 0.05 was used as the cut-off criterion

### Statistical analysis

Statistical analysis is detailed under relevant figures. Protein levels as measured using Western blots analysis were quantified using Image Quant. All experiments were performed at least in three independent biological replicates unless stated otherwise, and representative images are shown.

## Supporting information

Chemical SI

Table S1 -R4VPs regulated genes

## Acknowledgments

We thank Igor Voldavsky for pathological analysis, Gina Gotthardt and Stefan Müller for sharing unpublished data. Avi Maimon, Sima Lev, Yuval Shaked, for discussions and reagents. We thank the Azrieli found for supporting genomic experiment performed at the ATGC genomic center. AOV is a Rubinstein PhD fellow

## Author contributions

Conceptualization: AO, AB

Methodology: AVO, DP, RN EBL, MDE, NP, AB, AO

Investigation: YAA, AVO, DP, R, EBL

Supervision: MDE, AB, AO

Writing—original draft: MDE, AB, AO

Writing—review & editing: AOV, DP, RN, NP, EBL MDE, AB, AO

## Ethical statement

All experiments involving human patients were performed with patient’s informed consents and according to Helsinki ethics committee permit:

## Data and materials availability

All data are available in the main text or the supplementary materials. The RNA-seq-data is available at: GSE284185

## Supplementary Materials

1. Supplemental Figures S1A-C, S3
2. Supplemental chemical SI
3. Supplemental Table S1

