## Supplemental data- and figures for "Eliminating Aggressive Cancers via PROTAC-like Inducers of Ferroptosis"

**This PDF file includes:**

1. Figs. S1A, S1B, S1C, anc
2. Chemical SI

**Other Supplementary Materials for this manuscript include the following:**

Data S1 Excel file: R4VPL3-1 regulated genes

**AVO et al. Figure S1A**

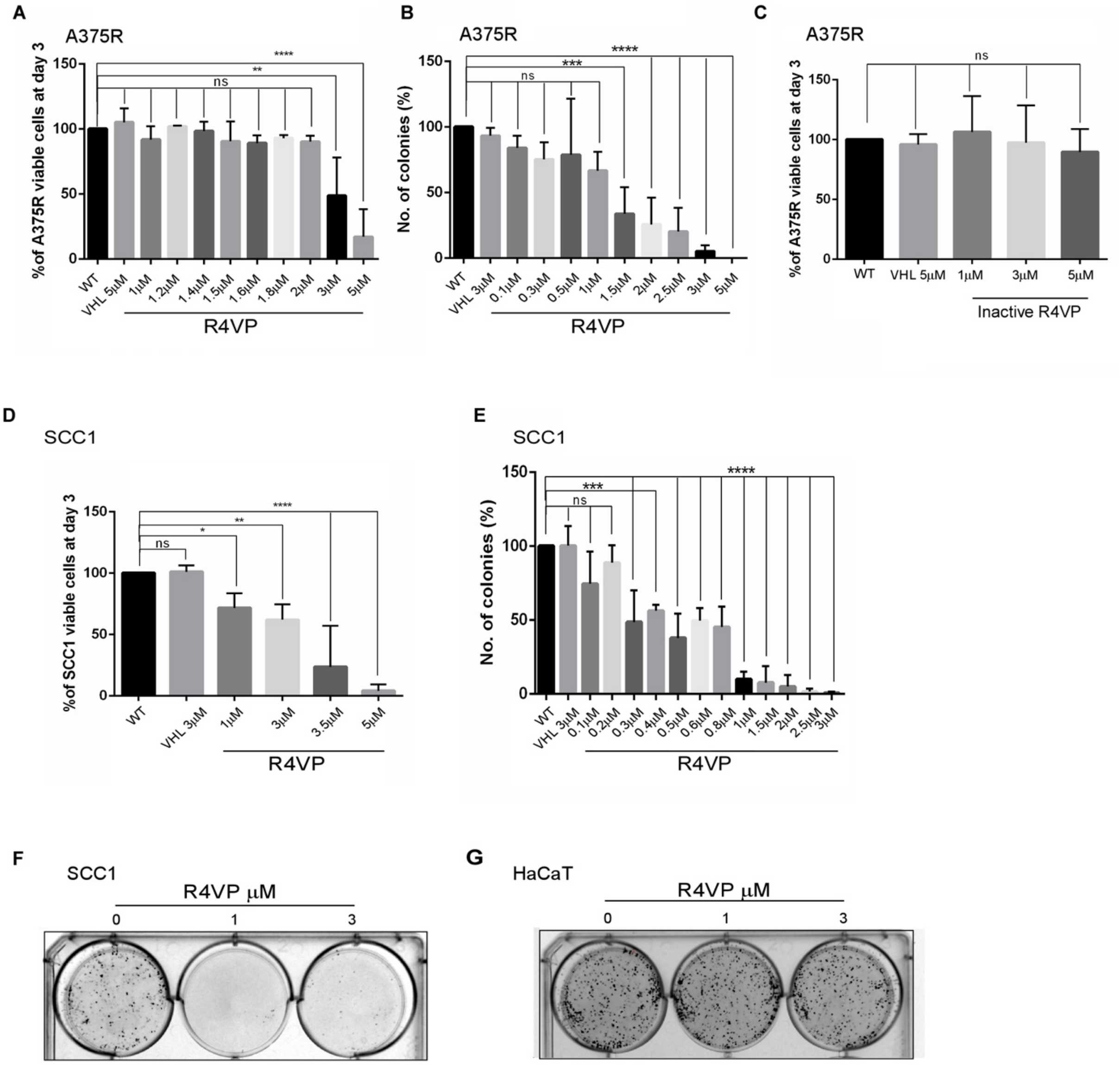


**Figure S1A: R4VP force the degradation of RNF4 and reduces survival of cancer cells. (A,B)** R4VP but not VHL-r compound, inhibits the proliferation (A) and SFA (B) of A375R melanoma cells. **(C)** Inactive R4VP lacking a critical Chloro atom has no impact on A375R cells proliferation . **(D)** R4VP treatment, but not VHL-only inhibit proliferation of human skin cancer cells SSC1. **(E-G)** R4VP inhibits SFA of SSC1 cells (E, F) but had no impact on non-tumorigenic human HaCat cell line (G). Cell proliferation was measured indirectly by ATP-Lite assay. In all experiments n=3 and ***= $p<0.001$ **=$p<0.01$,*=$p<0.1$, statistical analysis was performed using 1way Anova Dunnett's multiple comparisons test.

**AOV et al. Figure S1B**

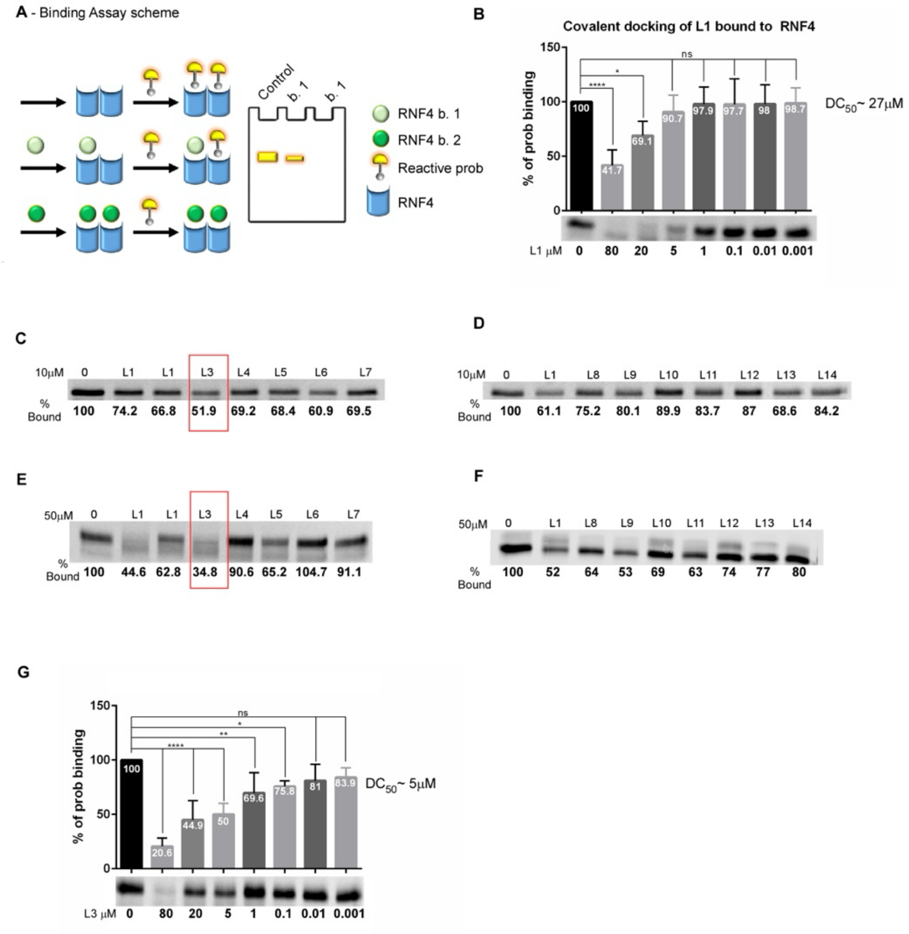


**Figure S1B: Screening of for an enhanced RNF4 binding moiety.** We generated CCW16-related compounds and measured their ability to directly bind bacterially purified RNF4 *in vitro* using a 5-TAMRIA -Iodoacetamide (T-I) labeling and displacement assay enabling quantitative visualization of RNF4 (39). Using this assay, we determined that the binding of the original CCW16 molecule (termed L1) to RNF4 resulted in reduced signal in a dose-dependent manner, with a 50% binding inhibition concentration of ~27μM. Using this assay, we compared the binding of L1 to newly developed L1-related molecules. Of the compound tested we identify L3 as an RNF4 binder with improved binding affinity with 50% T-I inhibition at ~5μM. **(A)** Schematic diagram of a direct 5-TAMRA-Iodacetmide dye displacement assay by the RNF4 binding moiety. **(B)** Upper panel: concentration dependent dye displacement by the L1, n=3, ***= $p<0.001$. Lower panel: Representative experiment. **(C-F)** Screening of TI-displacement of potential RNF4 binders at 10μM (C, D) and 50μM (E, F). **(G)** Upper panel: concentration dependent dye displacement by L3, an improved RNF4 binding moiety n=3, ****= $p<0.0001$. Lower pan is a representative experiment.

**AOV et al. Figure S1C**

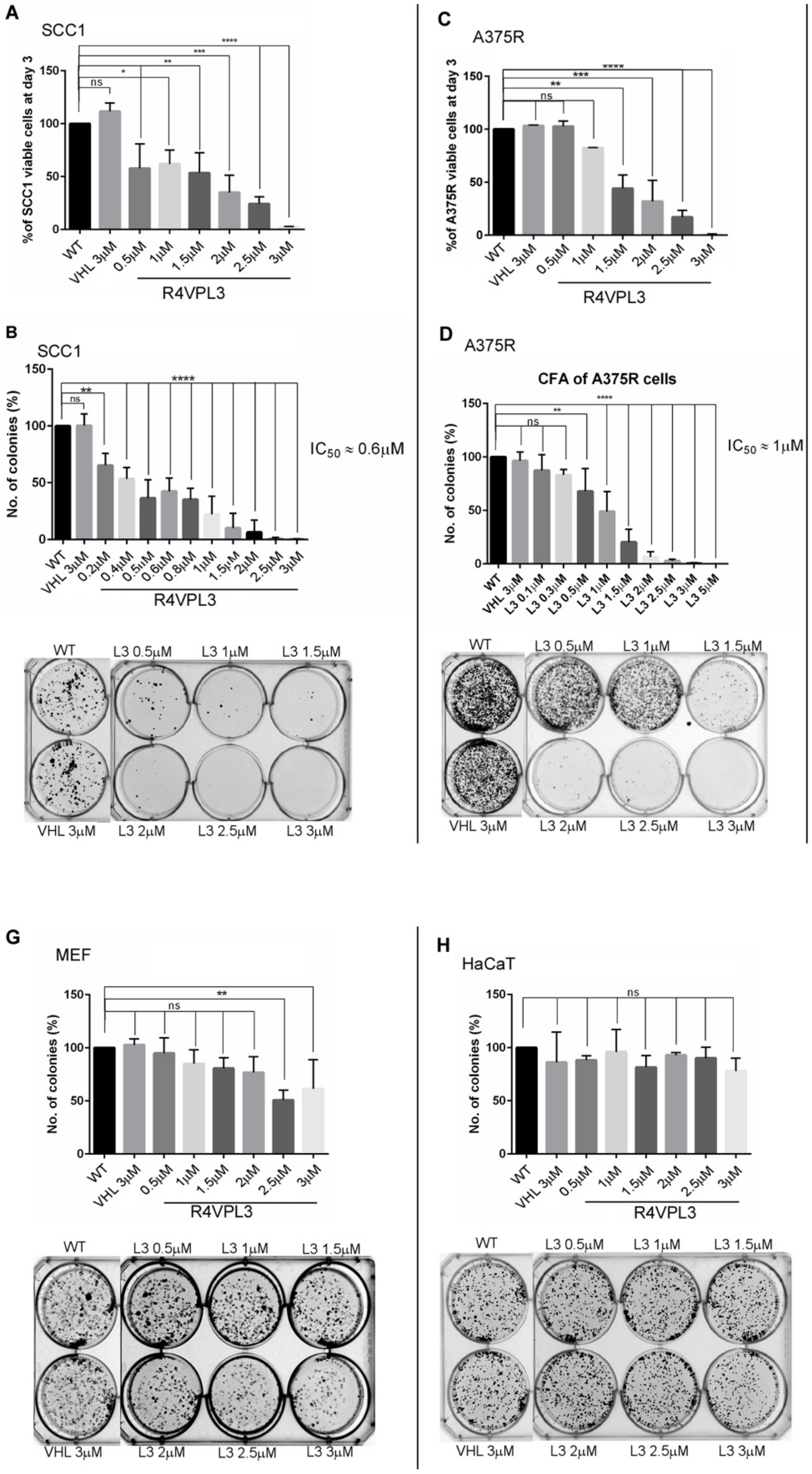

**Figure S1C: Anti-cancer activities of R4VPL3, R4VPL3-2 and R4VPL3-3.** Based on L3, we synthesized R4VPL3, a VHL-dependent PROTAC-like using L3 as the RNF4 binder that was more potent than L1 and retained its selective activity against cancer cells but had no impact on proliferation or SFA of MEFs. **(A-D)** Dose-dependent effect of R4VPL3 on viability and SFA of SCC1 cells (A, B) and A375R melanoma (C, D) cells. Lower panels in A and C are representative experiments. Cell proliferation was measured indirectly by MTT (B, D). **(E)** R4VPL3-2 had no impact on SFA of human melanoma A375R cells. **(F)** R4VPL3-3 inhibited SFA of A375R cells. **(G, H)** R4VPL3 had minimal or no impact on SFA of non-tumorigenic cells; Mouse embryonic fibroblasts (MEFs; G)) and HaCaT (H). In all experiments n=3 and ***=p<0.001, **=p<0.01. Statistical analysis was performed using 1way Anova Dunnett's multiple comparisons test. (C, n=2; B-D, n=3; E, n=4; F-G, n=3).

**AOV et al. Figure S3**

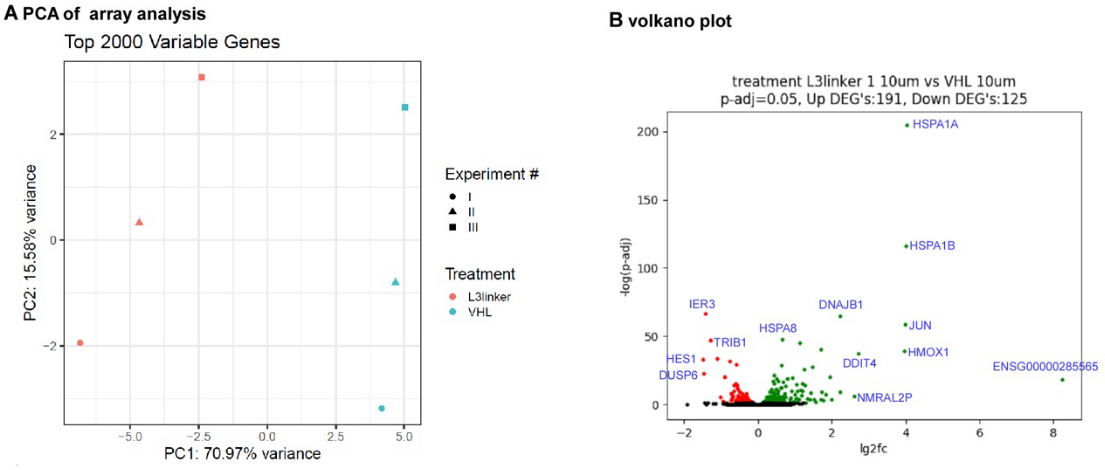


**Figure S3: Impact of R4VPL3-1 on gene expression.** RNA-Seq results comparing changes in gene expression signature between human melanoma A375R cells treated with 10µM of VHL-only moiety or R4VPL3-1. **(A)** Three principal components (PCA) analysis of three experimental biological repeats. **(B)** Volcano plots of changes in gene expression.

See methods for experimental and analysis details.
