## Supplementary material for "Eliminating Aggressive Cancers via PROTAC-like Inducers of Ferroptosis": Chemical SI

#### 2. Chemical SI

##### Materials and Methods

Materials were purchased from commercial vendors and used without further purification. All reactions were generally carried out under inert atmosphere unless otherwise noted. TLC was performed on Merck Kieselgel 60 F254 plates, and spots were visualized under UV light. Products were purified by column chromatography on silica gel (100-200 mesh, Merck). $^{1}$H and $^{13}$C NMR spectra were recorded on Brüker ADVANCE 400 (400 MHz and 100 MHz), instrument using deuterated solvents as detailed and at ambient probe temperature (300 K). Chemical shifts are reported in parts per million (ppm) and are referred to the residual solvent peak. The following notations are used: singlet (s); doublet (d); triplet (t); quartet (q); multiplet (m); broad (br). Coupling constants are quoted in Hertz and are denoted as J. Mass spectra were recorded on a Micromass® Q-Tof (ESI) spectrometer. HPLC was performed on a Thermo instrument (Dionex Ultimate 3000) using analytical Thermo Scientific (Hypersil Gold, C18, 3 µm, 4.6 × 150 mm) columns at flow rate of 1.2 ml/min. Preparative HPLC was performed on a Thermo Scientific instrument ultimate 3000 using Waters XSelect C18 (10µm, 19 × 250 mm) and semi preparative HPLC was performed on a Thermo Scientific instrument (Spectra System SCM1000) using XBridge BEH300 C4 (10 µm, 150 × 10 mm) and Jupiter C4 (10 µm, 300 Å, 250 × 10 mm) column, at flow rate of 15 and 4 mL/min S2 respectively. Buffer A: 0.1% TFA in water; buffer B: 0.1% TFA in acetonitrile. (Method: 0-60% B in 33 min).

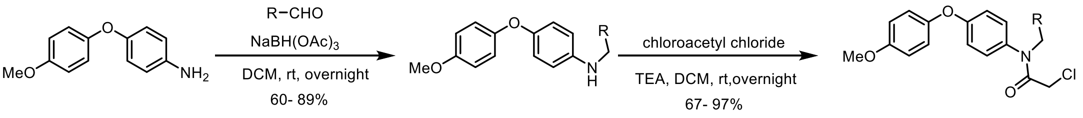


**General procedure 1 for the synthesis of Amine 1-14**: 4-(4-methoxyphenoxy) aniline (500 mg, 2.32 mmol) was dissolved in anhydrous DCM (10 mL) in a round bottom flask. aldehyde (2.32 mmol) was added and the solution was stirred at room temperature for 2h under nitrogen. Sodium triacetoxyborohydride (737.6 mg, 3.48 mmol) was added and the reaction mixture was stirred overnight. After completion, the reaction mixture was extracted with ethyl acetate. The organic layers were washed with brine and dried over anhydrous $Na_2SO_4$. After evaporation the crude residue was purified on a silica gel column (10-40% EtOAc/hexanes).

**General procedure 2 for the synthesis of Ligand 1-14**: To a solution of **corresponding amine** (0.5 mmol) in dry DCM (10 mL), 2-chloroacetyl chloride (67 mg, 0.6 mmol) and triethylamine (84 µL, 0.6 mmol) were added and the reaction mixture was stirred overnight at room temperature. After completion, the crude residue was purified by silica gel column (10-50% EtOAc/hexanes).

**N-benzyl-4-(4-methoxyphenoxy)aniline (Amine 1):** Following the GP-1, the reaction of 4-(4-methoxyphenoxy)aniline (500 mg, 2.32 mmol) and benzaldehyde(246.2 mg, 2.32 mmol) and reduction with sodium triacetoxyborohydride afforded **Amine 1** (425 mg, 60%) as a light yellow solid.

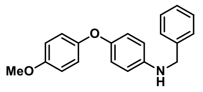


$^{1}$H NMR (400 MHz, $CDCl_3$): δ 7.38-7.37(m, 2H), 7.35- 7.33(m, 2H), 7.26(s, 1H), 6.91- 6.88(m, 2H), 6.86(s, 1H), 6.84- 6.83(m, 2H), 6.81(s, 1H), 6.62-6.60(m, 2H), 4.30(s, 2H), 3.78(s, 3H). $^{13}$C NMR (100 MHz, $CDCl_3$): δ 154.9, 152.3, 149.3, 144.3, 139.4, 128.7, 127.6, 120.1, 118.9, 114.6, 113.8, 55.7, 49.0

**N-benzyl-2-chloro-N-(4-(4-methoxyphenoxy)phenyl)acetamide (Ligand 1):** Following the GP-2, the reaction of N-benzyl-4-(4-methoxyphenoxy)aniline (**Amine 1**) (152.7 mg, 0.5 mmol) with 2-chloroacetyl chloride (67 mg, 0.6 mmol) afforded **Ligand 1** (162 mg, 85%) as a colourless liquid. $^{1}$H NMR (400 MHz, $CDCl_3$): δ 7.31-7.30(m, 3H), 7.26-7.23(m, 2H), 7.03-7.01(m, 2H), 6.96-6.93(m, 4H), 6.90- 6.88(m, 2H), 4.90(s, 2H), 3.91(s, 2H), 3.84(s, 3H). $^{13}$C NMR (100 MHz, $CDCl_3$): δ 166.4, 158.9, 156.5, 148.8, 136.6, 134.7, 129.5, 129.0, 128.5, 127.7, 121.5, 117.8, 115.1, 55.6, 53.8, 42.2.

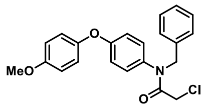

**N-(2,4-dimethoxybenzyl)-4-(4-methoxyphenoxy)aniline (Amine 2):** Following the GP-1, the reaction of 4-(4-methoxyphenoxy)aniline (500 mg, 2.32 mmol) and 2,4-Dimethoxybenzaldehyde (385.5 mg, 2.32 mmol) and reduction with sodium triacetoxyborohydride afforded **Amine 2** (600 mg, 71%) as a white solid. $^{1}$H NMR (400 MHz, $CDCl_3$): δ 7.21- 7.19(m, 1H), 6.91- 6.89(m, 2H), 6.86(s, 1H), 6.84(s, 2H), 6.81(s, 1H), 6.63(s, 1H), 6.61(s, 1H), 6.49- 6.48(m, 1H), 6.46- 6.43(m, 1H). 4.22(s, 2H), 3.84(s, 3H), 3.80(s, 3H), 3.78(s, 3H). $^{13}$C NMR (100 MHz, $CDCl_3$): δ 160.3, 158.5, 154.9, 152.4, 149.1, 144.6, 129.4, 120.1, 118.9, 114.6, 114.2, 103.9, 98.6, 55.7, 55.4, 55.4, 43.9.

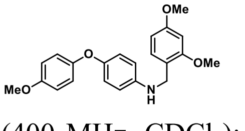

**2-chloro-N-(2,4-dimethoxybenzyl)-N-(4-(4-methoxyphenoxy)phenyl)acetamide (Ligand 2):** Following the GP-2, the reaction of N-(2,4-dimethoxybenzyl)-4-(4-methoxyphenoxy)aniline (**Amine 2**) (182.7 mg, 0.5 mmol) with 2-chloroacetyl chloride (67 mg, 0.6 mmol) afforded **Ligand 2** (181 mg, 82%) as a brown liquid. $^{1}$H NMR (400 MHz, $CDCl_3$): δ 7.13(d, 1H, $J$ = 8.3), 6.93(s, 1H), 6.91(s, 2H), 6.88(s,1H), 6.86(s, 1H), 6.84(s, 1H), 6.80(s, 1H), 6.78(s, 1H), 6.36(dd, 1H, $J$ = 8.3, 2.3), 6.29(d, 1H, $J$ = 2.3), 4.82(s, 2H), 3.83(s, 2H), 3.74(s, 3H), 3.71(s, 3H), 3.52(s, 3H). $^{13}$C NMR (100 MHz, $CDCl_3$): δ 166.3, 160.5, 158.6, 158.5, 156.4, 149.1, 134.9, 131.5, 129.6, 121.2, 117.6, 115.0, 104.2, 98.2, 55.6, 55.3, 55.1, 47.7, 42.2.

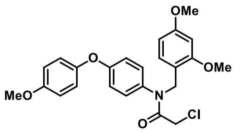

**4-(((4-(4-methoxyphenoxy)phenyl)amino)methyl)-N,N-dimethylaniline (Amine 3):** Following the GP-1, the reaction of 4-(4-methoxyphenoxy)aniline (500 mg, 2.32 mmol) and 4-(Dimethylamino)benzaldehyde (346.1 mg, 2.32 mmol) and reduction with sodium triacetoxyborohydride afforded **Amine 3** (566 mg, 70%) as a white solid. $^{1}$H NMR (400 MHz, $CDCl_3$): δ 7.24(s, 1H), 6.91(s, 1H), 6.89- 6.88(s, 1H), 6.86(s, 1H), 6.84-6.81(m, 3H), 6.74(s, 1H), 6.72(s, 2H), 6.62- 6.60(m, 2H), 4.17(s, 2H), 3.77(s, 3H), 2.94(s, 6H). $^{13}$C NMR (100 MHz, $CDCl_3$): δ 154.9, 152.4, 149.0, 144.6, 128.8, 120.2, 118.9, 114.6, 113.8, 112.9, 112.8, 55.7, 48.6, 40.8.

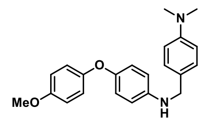

**2-chloro-N-(4-(dimethylamino)benzyl)-N-(4-(4-methoxyphenoxy)phenyl)acetamide (Ligand 3):** Following the GP-2, the reaction of 4-(((4-(4-methoxyphenoxy)phenyl)amino)methyl)-N,N-dimethylaniline (**Amine 3**) (174 mg, 0.5 mmol) with 2-chloroacetyl chloride (67 mg, 0.6 mmol) afforded **Ligand 3** (170 mg, 80%) as a brown solid. $^{1}$H NMR (400 MHz, $CDCl_3$): δ 7.06(s, 1H), 7.03(s,1H), 6.99(s, 1H), 6.97(s, 1H), 6.90(s, 2H), 6.88(s, 2H), 6.85(s, 1H), 6.83(s, 1H), 6.62(s, 1H), 6.60(s, 1H). 4.75(s, 2H), 3.84(s, 2H), 3.79(s, 3H), 2.90(s, 6H). $^{13}$C NMR (100 MHz, $CDCl_3$): δ 166.2, 158.8, 156.5, 150.0, 149.0, 134.8, 130.2, 129.7, 124.5, 121.5, 117.8, 115.0, 112.5, 55.7, 53.3, 42.3, 40.6.

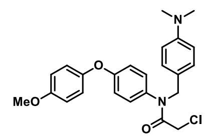

**N,N-diethyl-4-(((4-(4-methoxyphenoxy)phenyl)amino)methyl)aniline (Amine 4):** Following the GP-1, the reaction of 4-(4-methoxyphenoxy)aniline (500 mg, 2.32 mmol) and 4-(Diethylamino)benzaldehyde (411.2 mg, 2.32 mmol) and reduction with sodium triacetoxyborohydride afforded **Amine 4** (570 mg, 65%) as a brown solid. $^{1}$H NMR (400 MHz, $CDCl_3$): δ 7.23- 7.21(m, 2H), 6.91- 6.89(m, 2H), 6.87- 6.85(m, 2H), 6.83-

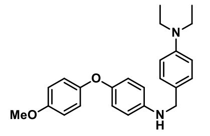

6.80(m, 2H), 6.68- 6.65(m, 2H), 6.63- 6.61(m, 2H), 4.15(s, 2H), 3.78(s, 3H), 3.36- 3.32(m, 4H), 1.18-1.14(m, 6H). $^{13}$C NMR (100 MHz, $CDCl_3$): δ 154.9, 152.4, 149.0, 147.2, 144.7, 129.1, 128.0, 120.2, 118.8, 114.6, 113.7, 111.9, 55.7, 48.6, 44.4, 12.6.

**2-chloro-N-(4-(diethylamino)benzyl)-N-(4-(4-methoxyphenoxy)phenyl)acetamide (Ligand 4):**

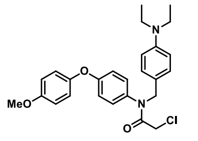

Following the GP-2, the reaction of N,N-diethyl-4-(((4-(4-methoxyphenoxy)phenyl)amino)methyl)aniline (**Amine 4**) (188.3 mg, 0.5 mmol) with 2-chloroacetyl chloride (67 mg, 0.6 mmol) afforded **Ligand 4** (193 mg, 75%) as a white solid. $^{1}$H NMR (400 MHz, $CDCl_3$): δ 7.03(s, 1H), 7.00(s, 2H), 6.98(s, 1H), 6.92-6.89(m, 4H), 6.87-6.85(m, 2H), 6.56-6.54(m, 2H), 4.73(s, 2H), 3.83(s, 2H), 3.81(s, 3H), 3.34-3.29(m, 4H), 1.13(t, 6H, J = 6.8). $^{13}$C NMR (100 MHz, $CDCl_3$): δ 166.0, 158.7, 156.5, 149.0, 147.3, 135.0, 130.4, 129.7, 123.1, 121.4, 117.8, 115.0, 111.6, 55.7, 53.4, 44.3, 42.3, 12.5.

**4-(4-methoxyphenoxy)-N-(4-methylbenzyl)aniline (Amine 5):** Following the GP-1, the reaction of

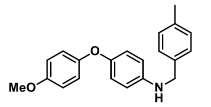

4-(4-methoxyphenoxy)aniline (500 mg, 2.32 mmol) and 4-Methylbenzaldehyde (278.8 mg, 2.32 mmol) and reduction with sodium triacetoxyborohydride afforded **Amine 5** (600 mg, 81%) as a white solid. $^{1}$H NMR (400 MHz, $CDCl_3$): δ 7.28-7.26(m, 2H), 7.17- 7.15(m, 2H), 6.91- 6.88(m, 2H), 6.86- 6.81(m, 4H), 6.61- 6.59(m, 2H), 4.26(s, 2H), 3.78(s, 3H), 2.35(s, 3H). $^{13}$C NMR (100 MHz, $CDCl_3$): δ 154.9, 152.3, 149.2, 144.4, 136.9, 136.3, 129.3, 127.6, 120.1, 118.9, 114.7, 113.8, 55.7, 48.7, 21.1.

**2-chloro-N-(4-(4-methoxyphenoxy)phenyl)-N-(4-methylbenzyl)acetamide (Ligand 5):** Following

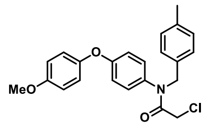

the GP-2, the reaction of 4-(4-methoxyphenoxy)-N-(4-methylbenzyl)aniline (**Amine 5**) (159.7 mg, 0.5 mmol) with 2-chloroacetyl chloride (67 mg, 0.6 mmol) afforded **Ligand 5** (164 mg, 83%) as a colourless liquid. $^{1}$H NMR (400 MHz, $CDCl_3$): δ 6.97- 6.96(m, 4H), 6.88(s, 1H), 6.86(s, 1H), 6.81(s, 1H), 6.79-6.78(m, 2H), 6.77(s, 1H), 6.75(s, 1H), 6.73(s, 1H), 4.81(s, 1H), 3.86(s, 2H), 3.79(s, 3H), 2.30(s, 3H). $^{13}$C NMR (100 MHz, $CDCl_3$): δ 166.3, 158.9, 156.5, 148.9, 137.3, 134.7, 133.7, 129.6, 129.2, 129.0, 121.5, 117.8, 115.1, 55.7, 53.5, 42.2, 21.2.

**N-(4-(tert-butyl)benzyl)-4-(4-methoxyphenoxy)aniline (Amine 6):** Following the GP-1, the reaction

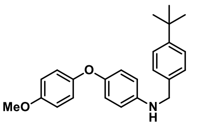

of 4-(4-methoxyphenoxy)aniline (500 mg, 2.32 mmol) and 4-tert-Butylbenzaldehyde (376.4 mg, 2.32 mmol) and reduction with sodium triacetoxyborohydride afforded **Amine 6** (750 mg, 89%) as a white solid. $^{1}$H NMR (400 MHz, $CDCl_3$): δ 7.39- 7.37(m, 2H), 7.33- 7.30(m, 2H), 6.91- 6.89(m, 2H), 6.87(s, 1H), 6.85- 6.84(m, 2H), 6.81(s, 1H), 6.63- 6.60(m, 2H), 4.26(s, 2H), 3.78(s, 3H), 1.33(s, 9H). $^{13}$C NMR (100 MHz, $CDCl_3$): δ 154.9, 152.3, 150.3, 149.2, 144.4, 136.3, 127.4, 125.6, 120.1, 118.9, 114.6, 113.8, 55.7, 48.6, 34.5, 31.4.

**N-(4-(tert-butyl)benzyl)-2-chloro-N-(4-(4-methoxyphenoxy)phenyl)acetamide (Ligand 6):**

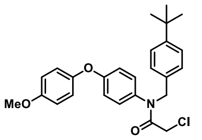

Following the GP-2, the reaction of N-(4-(tert-butyl)benzyl)-4-(4-methoxyphenoxy)aniline (**Amine 6**) (180.6 mg, 0.5 mmol) with 2-chloroacetyl chloride (67 mg, 0.6 mmol) afforded **Ligand 6** (171 mg, 78%) as a white solid. $^{1}$H NMR (400 MHz, $CDCl_3$): δ 7.30(s, 1H), 7.28(s, 1H), 7.15(s, 1H), 7.12(s, 1H), 7.01(s, 1H), 6.98(s, 1H), 6.95(s, 1H), 6.93(s, 1H), 6.92(s, 1H), 6.89(s, 1H), 6.87(s, 1H), 6.85(s, 1H), 4.82(s, 2H), 3.86(s, 2H), 3.81(s, 3H), 1.29(s, 9H). $^{13}$C NMR (100 MHz, $CDCl_3$): δ 166.4, 158.9, 156.5, 150.6, 149.0, 135.0, 133.5, 129.6, 128.7, 125.4, 121.5, 117.8, 115.0, 55.7, 53.6, 42.1, 34.5, 31.3.

**4-(4-methoxyphenoxy)-N-(4-(piperidin-1-yl)benzyl)aniline (Amine 7):** Following the GP-1, the reaction of 4-(4-methoxyphenoxy)aniline (500 mg, 2.32 mmol) and 4-(1-Piperidinyl)benzaldehyde (439 mg, 2.32 mmol) and reduction with sodium triacetoxyborohydride afforded **Amine 7** (681 mg, 76%) as a white solid. $^{1}$H NMR (400 MHz, $CDCl_3$): δ 7.31(s, 2H), 6.98(s, 4H), 6.91(s, 4H), 6.68(s, 2H), 4.25(s, 2H), 3.84(s, 3H), 3.22(s, 4H), 1.78(s, 4H), 1.65(s, 2H). $^{13}$C NMR (100 MHz, $CDCl_3$): δ 154.9, 152.4, 151.6, 149.1, 144.6, 129.7, 128.6, 120.2, 118.9, 116.6, 114.7, 113.8, 55.7, 50.8, 48.5, 25.9, 24.4.

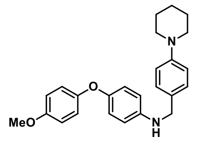

**2-chloro-N-(4-(4-methoxyphenoxy)phenyl)-N-(4-(piperidin-1-yl)benzyl)acetamide (Ligand 7):** Following the GP-2, the reaction of 4-(4-methoxyphenoxy)-N-(4-(piperidin-1-yl)benzyl)aniline (**Amine 7**) (194.3 mg, 0.5 mmol) with 2-chloroacetyl chloride (67 mg, 0.6 mmol) afforded **Ligand 7** (165 mg, 71%) as a white solid. $^{1}$H NMR (400 MHz, $CDCl_3$): δ 7.06(s, 1H), 7.04(s, 1H), 6.99(s, 1H), 6.97(s, 1H), 6.90(s, 1H), 6.89-6.86(m, 2H), 6.84 (s, 2H), 6.83-6.82(m, 2H), 6.79(s, 1H), 4.75(s, 2H), 3.83(s, 2H), 3.79(s, 3H), 3.11(t, 4H, $J$ = 5.3), 1.69-1.65(m, 4H), 1.56-1.55(m, 2H). $^{13}$C NMR (100 MHz, $CDCl_3$): δ 166.18, 158.8, 156.5, 151.6, 148.9, 134.8, 130.0, 129.7, 127.0, 121.4, 117.8, 116.1, 115.0, 55.7, 53.3, 50.4, 42.3, 25.8, 24.3.

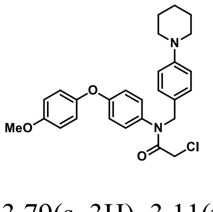

**4-(4-methoxyphenoxy)-N-(oxazol-4-ylmethyl)aniline (Amine 8):** Following the GP-1, the reaction of 4-(4-methoxyphenoxy)aniline (500 mg, 2.32 mmol) and 4-Oxazolecarboxaldehyde (225.2 mg, 2.32 mmol) and reduction with sodium triacetoxyborohydride afforded **Amine 8** (516 mg, 75%) as a brown solid. $^{1}$H NMR (400 MHz, $CDCl_3$): δ 7.87(s, 1H), 7.58(s, 1H), 6.90- 6.81(m, 6H), 6.64- 6.61(m, 2H), 4.26(s, 2H), 3.77(s, 3H). $^{13}$C NMR (100 MHz, $CDCl_3$): δ 155.1, 152.1, 151.3, 149.8, 143.6, 138.4, 135.5, 119.9, 119.1, 114.7, 114.3, 55.7, 40.9.

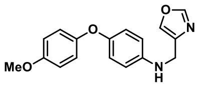

**2-chloro-N-(4-(4-methoxyphenoxy)phenyl)-N-(oxazol-4-ylmethyl)acetamide (Ligand 8):** Following the GP-2, the reaction of 4-(4-methoxyphenoxy)-N-(oxazol-4-ylmethyl)aniline (**Amine 8**) (148.2 mg, 0.5 mmol) with 2-chloroacetyl chloride (67 mg, 0.6 mmol) afforded **Ligand 8** (140 mg, 75%) as a white solid. $^{1}$H NMR (400 MHz, $CDCl_3$): δ 7.78(s, 1H), 7.63(s, 1H), 7.11-7.09(m, 2H), 6.97-6.95(m, 2H), 6.88-6.87(m, 4H), 4.72(s, 2H), 3.83(s, 2H), 3.76(s, 3H). $^{13}$C NMR (100 MHz, $CDCl_3$): δ 166.4, 159.0, 156.5, 150.9, 148.8, 137.6, 135.7, 135.0, 129.4, 121.5, 117.9, 115.1, 55.7, 45.7, 42.0.

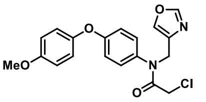

**4-(4-methoxyphenoxy)-N-(thiazol-2-ylmethyl)aniline (Amine 9):** Following the GP-1, the reaction of 4-(4-methoxyphenoxy)aniline (500 mg, 2.32 mmol) and 5-Thiazolecarboxaldehyde (265.5 mg, 2.32 mmol) and reduction with sodium triacetoxyborohydride afforded **Amine 9** (550 mg, 76%) as a white solid. $^{1}$H NMR (400 MHz, $CDCl_3$): δ 8.73(s, 1H), 7.81(s, 1H), 6.91- 6.82(m, 6H), 6.64- 6.62(m, 2H), 4.54(s, 2H), 3.78(s, 3H). $^{13}$C NMR (100 MHz, $CDCl_3$): δ 155.1, 152.9, 151.9, 150.3, 143.0, 141.1, 120.0, 119.2, 114.7, 114.5, 55.7, 41.5.

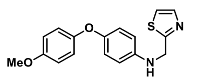

**2-chloro-N-(4-(4-methoxyphenoxy)phenyl)-N-(thiazol-2-ylmethyl)acetamide (Ligand 9):** Following the GP-2, the reaction of 4-(4-methoxyphenoxy)-N-(thiazol-2-ylmethyl)aniline (**Amine 9**) (156.2 mg, 0.5 mmol) with 2-chloroacetyl chloride (67 mg, 0.6 mmol) afforded **Ligand 9** (181 mg, 93%) as a white solid. $^{1}$H NMR (400 MHz, $CDCl_3$): δ 8.71(s, 1H), 7.56(s, 1H), 6.97-6.94 (m, 2H), 6.91(s, 1H), 6.88-6.86(m, 4H), 4.97(s,

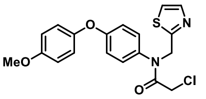

2H), 3.81(s, 2H), 3.76(s, 1H). $^{13}$C NMR (100 MHz, $CDCl_3$): δ 166.4, 159.4, 156.6, 154.5, 148.7, 143.4, 133.8, 132.9, 129.4, 121.6, 118.0, 115.1, 55.7, 45.8, 41.8.

**N-(cyclopentylmethyl)-4-(4-methoxyphenoxy)aniline (Amine 10):** Following the GP-1, the reaction

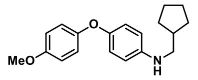

of 4-(4-methoxyphenoxy)aniline (500 mg, 2.32 mmol) and cyclopentanecarboxaldehyde (227.7 mg, 2.32 mmol) and reduction with sodium triacetoxyborohydride afforded **Amine 10** (550 mg, 80%) as a white solid. $^{1}$H NMR (400 MHz, $CDCl_3$): δ 6.90- 6.81(m, 6H), 6.59- 6.57(m,2H), 3.78(s, 3H), 3.01- 2.99(m, 2H), 2.17- 2.13(m, 1H), 1.84- 1.82(m, 2H), 1.65- 1.57(m, 4H), 1.29- 1.24(m, 2H). $^{13}$C NMR (100 MHz, $CDCl_3$): δ 154.9, 152.5, 148.9, 144.9, 120.2, 118.8, 114.6, 113.6, 55.7, 50.2, 39.5, 30.7, 25.3.

**2-chloro-N-(cyclopentylmethyl)-N-(4-(4-methoxyphenoxy)phenyl)acetamide (Ligand 10): :**

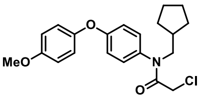

Following the GP-2, the reaction of N-(cyclopentylmethyl)-4-(4-methoxyphenoxy)aniline (**Amine 10**) (148.7 mg, 0.5 mmol) with 2-chloroacetyl chloride (67 mg, 0.6 mmol) afforded **Ligand 10** (181 mg, 97%) as a colourless liquid.$^{1}$H NMR (400 MHz, $CDCl_3$): δ 7.12(s, 1H), 7.09(s, 1H), 7.00(s, 1H), 6.98(s, 1H), 6.93(s, 1H), 6.91(s, 1H), 6.90(s, 1H), 6.88(s, 1H), 3.80(s, 2H), 3.78(s, 3H), 3.64(d, 2H, $J$ = 7.7), 2.05-1.97(m, 1H), 1.62-1.60(m, 4H), 1.48-1.45(m, 2H), 1.26-1.20(m, 2H). $^{13}$C NMR (100 MHz, $CDCl_3$): δ 166.4, 158.8, 156.5, 148.9, 135.0, 129.3, 121.5, 117.9, 115.0, 55.6, 54.3, 42.2, 37.8, 30.2, 25.2.

**N-(cyclopropylmethyl)-4-(4-methoxyphenoxy)aniline(Amine 11):** Following the GP-1, the reaction

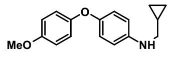

of 4-(4-methoxyphenoxy)aniline (500 mg, 2.32 mmol) and cyclopropanecarboxaldehyde (162.6 mg, 2.32 mmol) and reduction with sodium triacetoxyborohydride afforded **Amine 11** (400 mg, 64%) as a white solid. $^{1}$H NMR (400 MHz, $CDCl_3$): δ 6.91(s, 1H), 6.88(s, 1H), 6.87(s, 1H), 6.85- 6.83(m, 2H), 6.81(s, 1H), 6.60- 6.57(m, 2H), 3.78(s, 3H), 2.93(d, 2H, $J$ = 6.9), 1.14- 1.06(m, 1H), 0.56- 0.54(m, 2H), 0.25- 0.23(m, 2H). $^{13}$C NMR (100 MHz, $CDCl_3$): δ 154.9, 152.4, 149.0, 144.7, 120.2, 118.8, 114.6, 113.8, 55.7, 49.8, 10.9, 3.5.

**2-chloro-N-(cyclopropylmethyl)-N-(4-(4-methoxyphenoxy)phenyl)acetamide (Ligand 11):**

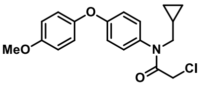

Following the GP-2, the reaction of **Amine 11** (134.7 mg, 0.5 mmol) with 2-chloroacetyl chloride (67 mg, 0.6 mmol) afforded **Ligand 11** (116 mg, 67%) as a brown liquid.$^{1}$H NMR (400 MHz, $CDCl_3$): δ 7.15(s, 1H), 7.12(s, 1H), 6.98(s, 1H), 6.96(s, 1H), 6.92(s, 1H), 6.90(s, 1H), 6.88(s, 1H), 6.86(s, 1H), 3.80(s, 2H), 3.76(s, 3H), 3.52-3.50(m, 2H), 0.93-0.89(m, 1H), 0.40-0.39(m, 2H), 0.11-0.10(m, 2H). $^{13}$C NMR (100 MHz, $CDCl_3$): δ 166.0, 158.8, 156.5, 148.9, 135.1, 129.7, 121.4, 117.9, 115.0, 55.6, 54.3, 42.2, 9.6, 3.7.

**4-(4-methoxyphenoxy)-N-neopentylaniline (Amine 12):** Following the GP-1, the reaction of 4-(4-

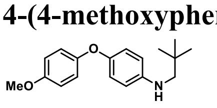

methoxyphenoxy)aniline (500 mg, 2.32 mmol) and Pivaldehyde (199.8 mg, 2.32 mmol) and reduction with sodium triacetoxyborohydride afforded **Amine 12** (494 mg, 75%) as a white solid. $^{1}$H NMR (400 MHz, $CDCl_3$): δ 6.90- 6.86(m, 3H), 6.84- 6.81(m, 3H), 6.61- 6.58(m, 2H), 3.78(s, 3H), 2.87(s, 2H), 1.00(s, 9H). $^{13}$C NMR (100 MHz, $CDCl_3$): δ 154.9, 152.5, 148.7, 145.3, 120.2, 118.7, 114.6, 113.6, 56.6, 55.7, 31.8, 27.7.

**2-chloro-N-(4-(4-methoxyphenoxy)phenyl)-N-neopentylacetamide (Ligand 12):** Following the GP-

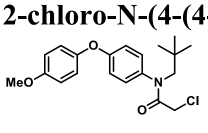

2, the reaction of 4-(4-methoxyphenoxy)-N-neopentylaniline (**Amine 12**) (142.7 mg, 0.5 mmol) with 2-chloroacetyl chloride (67 mg, 0.6 mmol) afforded **Ligand 12** (172 mg, 95%) as a colourless liquid.$^{1}$H NMR (400 MHz, $CDCl_3$): δ 7.17(s, 1H), 7.15(s, 1H), 6.99(s, 1H), 6.97(s, 1H), 6.91(s, 1H), 6.89(s, 2H), 6.87(s, 1H), 3.85(s, 2H), 3.78(s, 1H), 3.62(s, 1H), 0.85(s, 9H). $^{13}$C NMR (100 MHz, $CDCl_3$): δ 167.1, 158.3, 156.4, 149.0, 137.2, 129.1, 121.3, 117.8, 115.0, 61.2, 55.6, 42.3, 34.0, 28.4.

**N-isobutyl-4-(4-methoxyphenoxy)aniline (Amine 13):** Following the GP-1, the reaction of 4-(4-methoxyphenoxy)aniline (500 mg, 2.32 mmol) and isobutyraldehyde (167.3 mg, 2.32 mmol) and reduction with sodium triacetoxyborohydride afforded **Amine 13** (450 mg, 76%) as a white solid. $^{1}$H NMR (400 MHz, $CDCl_3$): δ 6.91- 6.88(m, 3H), 6.86(s, 1H), 6.85-6.81(m, 3H), 6.58- 6.56(m, 2H), 3.78(s, 3H), 2.91(d, 2H, $J$ = 6.7), 1.92- 1.85(m, 1H), 1.00- 0.98(m, 6H). $^{13}$C NMR (100 MHz, $CDCl_3$): δ 154.9, 152.5, 148.8, 144.8, 120.2, 118.8, 114.6, 113.6, 55.7, 52.5, 28.1, 20.5.

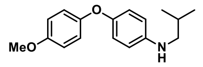

**2-chloro-N-isobutyl-N-(4-(4-methoxyphenoxy)phenyl)acetamide (Ligand 13):** Following the GP-2, the reaction of N-isobutyl-4-(4-methoxyphenoxy)aniline (**Amine 13**) (128.6 mg, 0.5 mmol) with 2-chloroacetyl chloride (67 mg, 0.6 mmol) afforded **Ligand 13** (137 mg, 79%) as a colourless liquid.$^{1}$H NMR (400 MHz, $CDCl_3$): δ 7.11(s, 1H), 7.09(s, 1H), 6.99(s, 1H), 6.96(s, 1H), 6.92(s, 1H), 6.90(s, 1H), 6.89(s, 1H), 6.88(s, 1H), 3.80(s, 2H), 3.77(s, 3H), 3.52(s, 1H), 3.50(s, 1H), 1.78-1.71(m, 1H), 0.88-0.86(m, 6H). $^{13}$C NMR (100 MHz, $CDCl_3$): δ 166.7, 158.8, 156.5, 148.9, 135.2, 129.2, 121.5, 117.9, 115.0, 56.9, 55.6, 42.1, 26.6, 19.9.

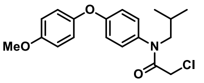

**4-(4-methoxyphenoxy)-N-propylaniline (Amine 14):** Following the GP-1, the reaction of 4-(4-methoxyphenoxy)aniline (500 mg, 2.32 mmol) and propionaldehyde (134.7 mg, 2.32 mmol) and reduction with sodium triacetoxyborohydride afforded **Amine 14** (430 mg, 72%) as a white solid. $^{1}$H NMR (400 MHz, $CDCl_3$): δ 6.91- 6.88(m, 2H), 6.87(s, 1H), 6.85-6.81(m, 3H), 6.59- 6.57(m, 2H), 3.78(s, 3H), 3.06(t, 2H, $J$ = 7), 1.67- 1.61(m, 2H), 1.00(t, 2H, $J$ = 7.4). $^{13}$C NMR (100 MHz, $CDCl_3$): δ 154.9, 152.4, 148.9, 144.7, 120.2, 118.8, 114.6, 113.7, 55.7, 46.5, 22.8, 11.7.

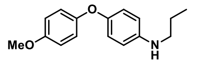

**2-chloro-N-(4-(4-methoxyphenoxy)phenyl)-N-propylacetamide (Ligand 14):** Following the GP-2, the reaction of 4-(4-methoxyphenoxy)-N-propylaniline (**Amine 14**) (128.7 mg, 0.5 mmol) with 2-chloroacetyl chloride (67 mg, 0.6 mmol) afforded **Ligand 14** (160 mg, 96%) as a brown liquid.$^{1}$H NMR (400 MHz, $CDCl_3$): δ 7.08(s, 1H), 7.06(s, 1H), 6.96(s, 1H), 6.94(s, 1H), 6.91(s, 1H), 6.88(s, 1H), 6.86(s, 1H), 6.84(s, 1H), 3.77(s, 2H), 3.76(s, 3H), 3.58(t, 2H, $J$ = 6.0), 1.51-1.45(m, 2H), 0.83(t, 3H, $J$ = 7.3). $^{13}$C NMR (100 MHz, $CDCl_3$): δ 166.2, 158.8, 156.5, 148.9, 135.0, 129.3, 121.4, 117.9, 115.0, 55.6, 51.6, 42.0, 20.6, 11.1.

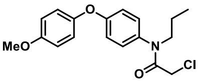

**Synthesis of N-benzyl-N-(4-(4-methoxyphenoxy)phenyl)acetamide (inactive Ligand):**

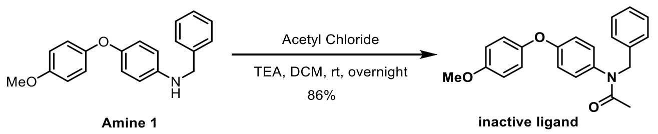


To a solution of N-benzyl-4-(4-methoxyphenoxy)aniline (**amine 1**)(152.7 mg, 0.5 mmol) in dry DCM (10 mL) acetyl chloride (39 mg, 0.6 mmol) and triethylamine (84 μL, 0.6 mmol) were added and the reaction mixture was stirred overnight at room temperature. After completion, the crude residue was purified by silica gel column (1:4 EtOAc/hexanes) to afford **inactive Ligand** (149 mg, 86%) as a colourless oil.$^{1}$H NMR (400 MHz, $CDCl_3$): δ 7.26-7.21(m, 5H), 6.99-6.97(m, 2H), 6.90-6.85(m, 6H), 4.85(s, 2H), 3.80(s, 3H), 1.90(s, 3H). $^{13}$C NMR (100 MHz, $CDCl_3$): δ 170.6, 158.1, 156.3, 149.2, 137.5, 137.0, 129.4, 128.8, 128.3, 127.3, 121.2, 117.7, 115.0, 55.6, 52.8, 22.7.

**Synthesis of 4-(((R)-1-((2R,4R)-4-hydroxy-2-((4-(4-methylthiazol-5 yl) benzyl) carbamoyl) pyrrolidin-1-yl)-3,3-dimethyl-1-oxobutan-2-yl)amino)-4-oxobutanoic acid (VHL acid):**

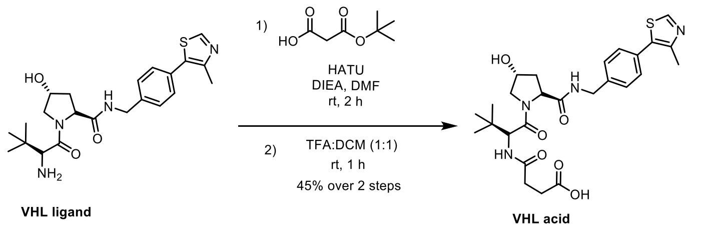


To a solution of **VHL ligand hydrochloride** (250 mg, 0.53mmol) in DMF(5 mL), HATU (244 mg, 0.64 mmol) and 4-(tert-butoxy)-4-oxobutanoic acid(112 mg, 0.64mmol) were added and stirred for 5 min at room temperature. To this, DIEA (277µL, 1.6 mmol) was added and continued the reaction for 2 h. After 2 h, the reaction mixture was diluted with ethyl acetate and ice-cold water. The organic layer was separated, dried over $Na_2SO_4$ and distilled under reduced pressure. To the solution of crude product in DCM(5mL), TFA(5mL) was added dropwise and stirred the reaction for 1 h at room temperature. After 1 h, the reaction mixture was distilled under reduced pressure. The crude was diluted with dichloromethane and distilled under reduced pressure (3 times). Finally, the dried crude product was kept under high vacuum for overnight. The crude product was further purified using preparative HPLC with a gradient of 0-60% ACN/$H_2O$ in 30 min system. The pure product **VHL acid** (127mg, 45%) was obtained as a white solid after the lyophilization. $^1$H NMR (400 MHz, D6-DMSO): $\delta$ 9.06 (s, 1H), 8.64 (t, 1H, $J$ = 6.0), 8.00 (d, 1H, $J$ = 9.2), 7.49-7.42 (m, 4H), 4.59 (d, 1H, $J$ = 9.2), 4.52-4.46 (m, 2H), 4.40 (broad, 1H), 4.29-4.24 (m, 1H), 3.74-3.66 (m, 2H), 2.56-2.55 (m, 4H), 2.46-2.38 (m, 3H), 2.15-2.06(m, 1H), 1.98-1.92(m, 1H), 0.98 (s, 9H). $^{13}$C NMR (400 MHz, D6-DMSO): $\delta$ 174.3, 172.4, 171.3, 170.0, 159.0, 158.6, 152.0, 147.9, 140.0, 129.1, 127.9, 117.1, 114.2, 69.3, 59.1, 56.9, 42.1, 38.3, 35.8, 30.2, 29.7, 26.8, 16.3. HRMS (m/z): $[M+H]^+$calcd. for $C_{26}H_{35}N_4O_6S$, 531.2277; found 531.2277.

**Synthesis of N1-(4-(4-(4-(N-benzyl-2-chloroacetamido)phenoxy)phenoxy)butyl)-N4-((R)-1-((2R,4R)-4-hydroxy-2-((4-(4-methylthiazol-5-yl)benzyl)carbamoyl)pyrrolidin-1-yl)-3,3-dimethyl-1-oxobutan-2-yl)succinimide (R4VP):**

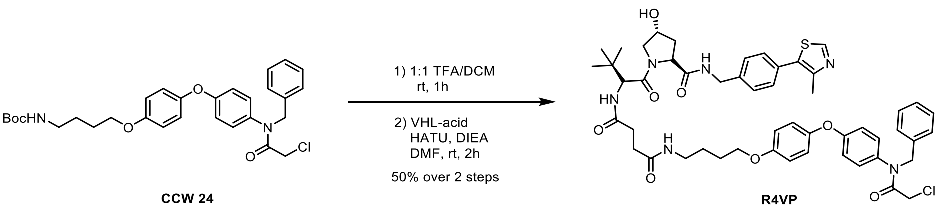


To a solution of **CCW 24**(120.9 mg, 0.227 mmol) in DCM(5mL), TFA (5 mL) was added dropwise and stirred the reaction for 1 h at room temperature. After 1 h, the reaction mixture was distilled under reduced pressure. The crude was diluted with dichloromethane and distilled under reduced pressure (3 times). Then to the solution of this crude product in DMF (5 mL), **VHL acid** (100 mg, 0.227 mmol), HATU (95 mg, 0.250 mmol) and DIEA (160 µL, 0.908 mmol) were added and continued the reaction for 2h at room temperature and was monitored by TLC. The reaction mixture was diluted with cold water and ethyl acetate. The organic layer was separated, dried over $Na_2SO_4$ and distilled under reduced pressure. Then crude was purified by preparative HPLC with a gradient of 0-60% ACN/$H_2O$ in 30 min system with a reverse phase C18 column to afford the **R4VP** (78.1 mg, 36%) as white solid. $^1$H NMR (400 MHz, $CDCl_3$): $\delta$ 9.00 (s, 1H), 7.39 (t, 1H, $J$ = 6.0 ), 7.29 (q, 4H, $J$ = 13.2), 7.20-7.18 (m, 4H), 7.13-7.10 (m, 2H), 6.90-6.88 (m, 2H), 6.84-6.76 (m, 6H), 6.24 (t, 1H, $J$ = 4.8), 4.92 (s, 1H), 4.78 (s, 2H), 4.62 (t, 1H, $J$ = 8.0), 4.48-4.44 (m, 2H), 4.39-4.37 (m, 1H), 4.30-4.25 (m, 1H), 3.98 (d, 1H, $J$ = 11.6), 3.87 (t, 2H, $J$ = 6.0), 3.80 (s, 2H), 3.57-3.53 (m, 1H), 3.22-3.19 (m, 2H), 2.47-2.30 (m, 8H), 2.14-2.08

(m, 1H), 1.73-1.69 (m, 2H), 1.62-1.58 (m, 2H), 0.89 (s, 9H). $^{13}C$ NMR (100 MHz, $CDCl_3$): $\delta$ 172.3, 171.7, 170.8, 170.2, 170.1, 165.7, 159.4, 159.0, 157.9, 154.7, 151.1, 148.0, 144.6, 138.1, 135.4, 133.6, 132.5, 128.5, 128.4, 128.0, 128.0, 127.5, 127.2, 126.7, 120.5, 116.8, 114.6, 69.2, 66.8, 57.8, 57.4, 55.8, 52.9, 42.1, 40.9, 38.5, 35.4, 33.9, 30.2, 30.1, 28.7, 25.5, 25.3, 25.1, 13.5. HRMS (m/z): calcd. for $C_{51}H_{60}ClN_6O_8S$, 951.3882; found 951.3876.

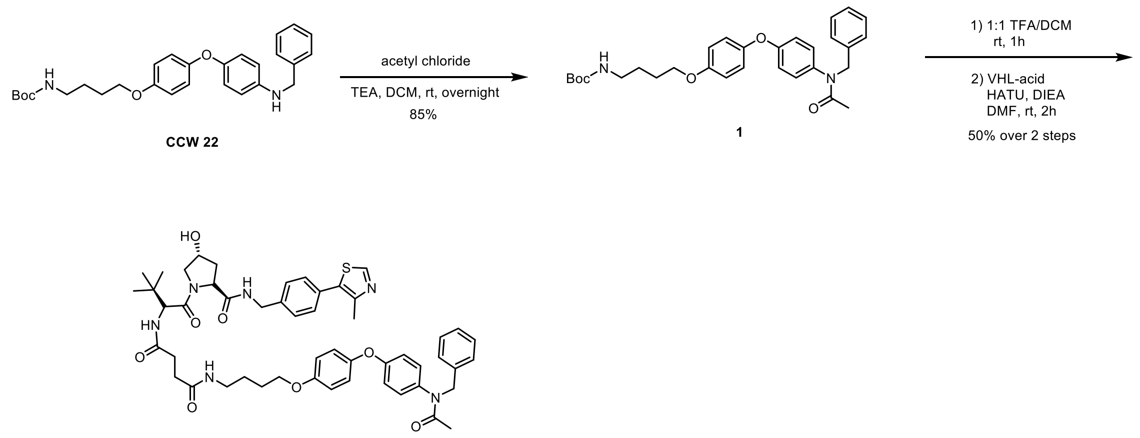


inactive R4VP

**Preparation of tert-butyl (4-(4-(4-(N-benzylacetamido)phenoxy)phenoxy)butyl)carbamate (1):** To a solution of **CCW 22** (200mg, 0.43 mmol) in DCM(5mL), acetyl chloride (40.7 mg, 0.52mmol) and triethylamine (Sigma, mmol) were added and the reaction mixture was stirred for overnight. Upon reaction completion the crude was purified by silica gel chromatography (30% EtOAc/hexanes) to yield 180 mg (82.5 %). $^{1}H$ NMR (400 MHz, $CDCl_3$): δ 7.23- 7.16(m, 5H), 6.96-6.92(m, 2H), 6.87- 6.80(m, 6H), 4.82(s, 4H), 4.51(s, 2H), 3.92(t, 2H, $J$ = 6.8), 3.17- 3.16(m, 2H), 1.78(t, 2H, $J$ = 6.8), 1.68-1.62(m, 2H), 1.41(s, 9H). $^{13}C$ NMR (100 MHz, $CDCl_3$): δ 170.5, 158.1, 157.7, 156.0, 155.6, 154.3, 150.1, 149.0, 129.4, 129.3, 128.7, 128.3, 127.3, 121.2, 117.9, 117.6, 115.7, 115.5, 73.4, 67.8, 52.7, 40.2, 28.4, 26.5, 22.6. HRMS (m/z): $[M+H]^+$ calcd. for $C_{30}H_{37}N_2O_5$, 505.2702; found: 505.2670.

**Preparation of N1-(4-(4-(4-(N-benzylacetamido)phenoxy)phenoxy)butyl)-N4-((R)-1-((2R,4R)-4-hydroxy-2-((4-(4-methylthiazol-5-yl)benzyl)carbamoyl)pyrrolidin-1-yl)-3,3-dimethyl-1-oxobutan-2-yl)succinimide (inactive R4VP):**

To the solution of tert-butyl (4-(4-(4-(N-benzylacetamido)phenoxy)phenoxy)butyl)carbamate (**1**)(100 mg, 0.2 mmol) in DCM(5mL), TFA(5mL) was added dropwise and stirred the reaction for 1 h at room temperature. After 1 h, the reaction mixture was distilled under reduced pressure. The crude was diluted with dichloromethane and distilled under reduced pressure (3 times). Then to the solution of this crude product in DMF(5mL), VHL-acid (106 mg, 0.2 mol), HATU (91 mg, 0.24 mmol) and DIPEA (108μl, 0.6 mmol) were added and continued the reaction for 2h at room temperature and was monitored by TLC. The reaction mixture was diluted with cold water and ethyl acetate. The organic layer was separated, dried over $Na_2SO_4$ and distilled under reduced pressure. Then crude was purified by preparative HPLC with a gradient of 0-60% ACN/$H_2O$ in 30 min system with a reverse phase C18 column to afford the **inactive R4VP** (92 mg, 50%) as white solid. $^{1}H$ NMR (400 MHz, $CDCl_3$): δ 9.03(s, 1H), 7.31- 7.26(m, 4H), 7.18($s_{br}$, 4H), 7.11- 7.09(m, 2H), 6.89- 6.87(2H), 6.79- 6.74(m, 5H), 6.20(s, 1H), 4.77(s, 3H), 4.53- 4.44(m, 3H), 4.38- 4.36(m, 1H), 4.29- 4.24(m, 2H), 4.00- 3.97(m, 1H), 3.87-

3.84(m, 2H), 3.56- 3.54(m, 1H), 3.21- 3.20(m, 2H), 2.47(s, 6H), 2.41- 2.31(m, 2H), 1.84(s, 3H), 1.72- 1.68(m, 2H), 1.63- 1.58(m, 2H), 0.88(s, 9H). $^{13}$C NMR (400 MHz, $CDCl_3$): δ 173.4, 172.8, 172.0, 171.8, 171.2, 158.4, 155.6, 152.2, 149.2, 139.2, 136.8, 136.3, 129.4, 129.2, 128.9, 128.5, 128.3, 127.6, 121.4, 117.8, 115.6, 70.2, 67.9, 58.8, 58.4, 56.8, 53.3, 43.2, 39.6, 36.4, 34.9, 31.2, 31.1, 26.6, 26.3, 26.1, 22.3, 14.3. HRMS (m/z): [M+H] $^+$ calcd. for $C_{51}H_{61}N_6O_8S$, 917.4272; found: 917.4265.

**Preparation of R4VPL3:**

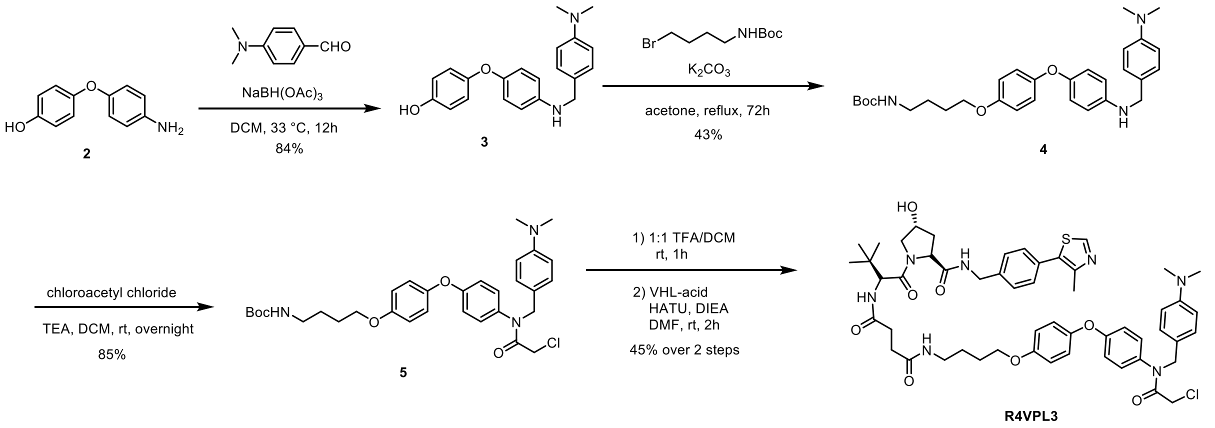


**Preparation of 4-(4-((4-(dimethylamino)benzyl)amino)phenoxy)phenol (3):**

To a solution of 4-(4-aminophenoxy) phenol (302 mg, 1.5 mmol) in 5 mL DCM, 4 (dimethylamino)benzaldehyde (224 mg, 1.5 mmol) was added and the solution was stirred at room temperature for 2h under nitrogen. Sodium triacetoxyborohydride (477 mg, 2.25 mmol) was added and the reaction mixture was stirred overnight. After completion, the reaction mixture was extracted with ethyl acetate. The organic layers were washed with brine and dried over anhydrous $Na_2SO_4$. After evaporation the crude residue was purified on a silica gel column (40% EtOAc/hexanes) and compound **3**(421 mg, 84%) was obtained as yellow liquid. (1H NMR (400 MHz, $CDCl_3$)**:** δ 7.30- 7.28(m, 2H), 6.91- 6.85(m, 4H), 6.79- 6.76(m, 4H), 6.67- 6.65(m, 2H), 4.22(s, 2H), 2.99(s, 6H). $^{13}$C NMR (100 MHz, $CDCl_3$): δ 152.3, 150.9, 150.1, 149.2, 144.5, 129.4, 128.8, 127.1, 120.2, 119.0, 116.6, 116.1, 114.0, 113.0, 48.7, 40.8.

**Preparation of tert-butyl (4-(4-(4-((4 (dimethylamino)benzyl)amino)phenoxy)phenoxy)butyl) carbamate (4):**

To a solution of **3**(679 mg, 2 mmol) in 10 mL Acetone tert-butyl (4-bromobutyl) carbamate(756 mg, 3 mmol), potassium carbonate(622 mg, 4.5 mmol) were added and the solution was put in reflux for 72h. After that the mixture was concentrated and purified by a silica gel column (30% EtOAc/hexanes) and compound **4**(435 mg, 43%) was obtained as yellow liquid. $^1$H NMR (400 MHz, $CDCl_3$)**:** δ 7.23(s, 1H), 7.11(d, 1H, $J$ =8.6), 6.89-6.88(m, 1H), 6.87-6.86(m, 2H), 6.84-6.83(m, 1H), 6.82-6.80(m, 2H), 6.79-6.78(m, 1H), 6.74-6.69(m, 2H), 6.62-6.60(m, 1H), 4.48(s, 2H), 3.94-3.91(m, 2H), 3.18(t, 2H, $J$ = 6.3), 2.94-2.92(m, 6H), 1.81-1.76(m, 2H), 1.67-1.63(m, 2H), 1.44(s, 9H). $^{13}$C NMR (100 MHz, $CDCl_3$): δ 156.0, 154.2, 152.4, 150.0, 149.7, 149.0, 144.6, 128.7, 127.8, 120.2, 118.8, 115.3, 113.7, 112.8, 79.1, 68.0, 53.9, 53.4, 48.5, 40.7, 28.4, 26.6. HRMS (m/z): [M+H] + calcd. for $C_{30}H_{40}N_3O_4$, 506.3019; found: 506.3007.

**Preparation of tert-butyl (4-(4-(4-(2-chloro-N-(4-(dimethylamino)benzyl)acetamido) phenoxy)phenoxy)butyl) carbamate (5):**

To a solution of **4** (101 mg, 0.2 mmol), chloro acetyl chloride (27 mg, 0.24 mmol), Triethyl amine (24 mg, 0.24 mmol) were added and the reaction mixture was stirred for overnight. Upon reaction completion the crude was purified by silica gel chromatography (2% MeOH/DCM) to give compound **5**(99 mg, 85%) as yellow liquid. $^{1}$H NMR (400 MHz, $CDCl_3$): δ 7.06-7.03(m,2H), 6.98-6.96(m, 2H), 6.89-6.83(m, 6H), 6.62-6.60(m, 2H), 4.75(s, 2H), 3.98(t, 2H, *J* = 6.1), 2.92(s, 6H), 1.84-1.80(m, 2H), 1.71-1.65(m, 2H), 1.44(s, 9H). $^{13}$C NMR (100 MHz, $CDCl_3$): δ 166.1, 156.0, 155.8, 150.0, 148.9, 134.8, 130.2, 129.7, 124.3, 121.4, 117.7, 115.6, 112.3, 79.2, 67.9, 53.3, 42.2, 40.5, 40.2, 28.4, 26.8, 26.6. HRMS (m/z): [M+H] + calcd. for $C_{32}H_{41}ClN_3O_5$, 582.2735; found: 582.2744.

**Preparation of N1-(4-(4-(4-(2-chloro-N-(4-(dimethylamino)benzyl)acetamido)phenoxy)phenoxy) butyl)-N4-((R)-1-((2R,4R)-4-hydroxy-2-((4-(4-methylthiazol-5-yl)benzyl)carbamoyl)pyrrolidin-1-yl)-3,3-dimethyl-1-oxobutan-2-yl)succinamide (R4VPL3):**

To the solution of **5** (35 mg, 0.06 mmol) in 2mL DCM, TFA(2mL) was added dropwise and stirred the reaction for 1 h at room temperature. After 1 h, the reaction mixture was distilled under reduced pressure. The crude was diluted with dichloromethane and distilled under reduced pressure (3 times). Then to the solution of this crude product in DMF(5mL), VHL-acid (37 mg, 0.07 mmol), HATU (27 mg, 0.07 mmol) and DIPEA (39 mg, 0.3 mmol) were added and continued the reaction for 2h at room temperature and was monitored by TLC. The reaction mixture was diluted with cold water and ethyl acetate. The organic layer was separated, dried over $Na_2SO_4$ and distilled under reduced pressure. Then crude was purified by preparative HPLC with a gradient of 0-60% ACN/$H_2O$ in 30 min system with a reverse phase C18 column to afford **R4VPL3** (27 mg, 45%) as a white solid. $^{1}$H NMR (400 MHz, $CDCl_3$): δ 9.02(s, 1H), 7.43(s, 1H), 7.41(s, 1H), 7.37-7.34(m, 6H), 7.07-7.05(m, 1H), 6.98-6.96(m, 2H), 6.93-6.91(m, 2H), 6.89-6.86(m, 2H), 6.85(s, 1H), 6.21(s, 1H), 4.85(s, 2H), 4.72-4.68(m, 1H), 4.60-4.51(m, 2H), 4.46-4.43(m, 1H), 4.37-4.32(m, 2H), 3.95(t. 2H, *J* = 6), 3.86(s, 2H), 3.62-3.58(m, 1H), 3.31-3.26(m, 2H), 3.15(s, 6H), 2.54-2.52(m, 4H), 2.48-2.42(m,2H), 2.19-2.14(m, 1H), 1.80-1.75(m, 2H), 1.71-1.64(m, 2H), 0.95(s, 9H). $^{13}$C NMR (100 MHz, $CDCl_3$): δ 172.5, 171.8, 171.0, 166.8, 161.3, 159.2, 155.8, 151.6, 148.8, 146.4, 143.7, 138.9, 134.4, 133.0, 130.9, 129.5, 129.3, 128.3, 121.6, 119.6, 118.0, 115.7, 70.2, 67.9, 58.6, 58.2, 56.7, 53.3, 45.5, 43.2, 41.8, 39.5, 34.9, 31.3, 31.2, 26.4, 14.9. HRMS (m/z): [M+H] + calcd. for $C_{53}H_{65}ClN_7O_8S$, 994.4304; found: 994.4299.

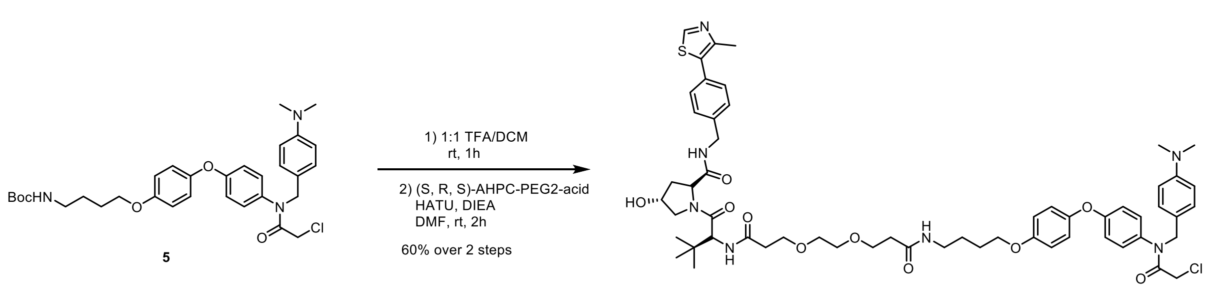


R4VP L3-1

**Preparation of (2R,4R)-1-((R)-2-(tert-butyl)-18-(4-(4-(2-chloro-N-(4 (dimethylamino)benzyl) acetamido) phenoxy)phenoxy)-4,13-dioxo-7,10-dioxa-3,14-diazaoctadecanoyl)-4-hydroxy-N-(4-(4-methylthiazol-5-yl)benzyl)pyrrolidine-2-carboxamide (R4VP L3-1):**

To the solution of **5** (35 mg, 0.06 mmol) in 2mL DCM, TFA(2mL) was added dropwise and stirred the reaction for 1 h at room temperature. After 1 h, the reaction mixture was distilled under reduced pressure. The crude was diluted with dichloromethane and distilled under reduced pressure (3 times).

Then to the solution of this crude product in DMF(5mL), (S, R, S)-AHPC-PEG2-acid (43 mg, 0.07 mmol), HATU (27 mg, 0.07 mmol) and DIPEA (39 mg, 0.3 mmol) were added and continued the reaction for 2h at room temperature and was monitored by TLC. The reaction mixture was diluted with cold water and ethyl acetate. The organic layer was separated, dried over $Na_2SO_4$ and distilled under reduced pressure. Then crude was purified by preparative HPLC with a gradient of 0-60% ACN/$H_2O$ in 30 min system with a reverse phase C18 column to afford **R4VP L3-1**(39 mg, 60%) as a white solid. $^1$H NMR (400 MHz, $CDCl_3$): δ 9.29(s, 1H), 8.21($s_{br}$, 3H), 7.53- 7.51(m, 2H), 7.46- 7.43(m, 5H), 7.40- 7.38(m, 2H), 7.02- 6.98(m, 3H), 6.96(s, 1H), 6.93- 6.90(m, 3H), 4.90(s, 2H), 4.71- 4.65(m, 2H), 4.61- 4.59(m, 2H), 4.41- 4.36(m, 1H), 4.16- 4.13(m, 1H), 3.97(t, 2H, $J$ = 6), 3.91(s, 2H), 3.76- 3.73(m, 4H), 3.65- 3.63(m, 4H), 3.33- 3.32(m, 2H), 3.23(s, 6H), 2.60(s, 4H), 2.57- 2.52(m, 4H), 2.37- 2.28(m, 1H), 1.83- 1.78(m, 2H), 1.73- 1.68(m, 2H), 1.03(s, 9H). $^{13}$C NMR (100 MHz, $CDCl_3$): δ 173.4, 171.6, 167.2, 160.7, 160.3, 159.3, 155.8, 152.7, 148.7, 144.3, 142.4, 139.7, 138.8, 134.4, 134.2, 131.1, 129.3, 129.2, 128.3, 121.6, 120.5, 118.0, 116.8, 115.7, 114.0, 70.2, 70.0, 67.8, 67.1, 66.8, 59.3, 58.2, 53.3, 46.4, 43.1, 41.7, 39.5, 36.9, 36.2, 35.9, 35.2, 26.5, 26.4, 26.3, 25.9, 13.9. HRMS (m/z): [M+H] + calcd. for $C_{57}H_{73}ClN_7O_{10}S$, 1082.4823; found: 1082.4863.

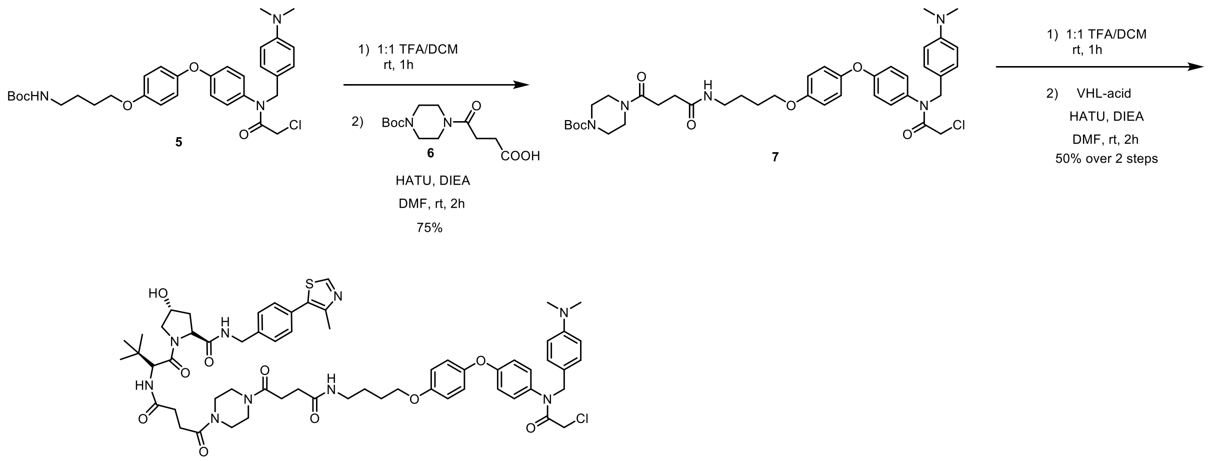


**Preparation of tert-butyl 4-(4-((4-(4-(4-(2-chloro-N-(4 (dimethylamino)benzyl)acetamido) phenoxy)phenoxy) butyl)amino)-4-oxobutanoyl)piperazine-1-carboxylate (7):**

To the solution of **5** (292mg, 0.5 mmol) in 5mL DCM, TFA(5mL) was added dropwise and stirred the reaction for 1 h at room temperature. After 1 h, the reaction mixture was distilled under reduced pressure. The crude was diluted with dichloromethane and distilled under reduced pressure (3 times). Then to the solution of this crude product in DMF(5mL), compound **6** (172 mg, 0.6 mmol), HATU (228 mg, 0.6 mmol) and DIPEA (261 µL, 1.5 mmol) were added and continued the reaction for 2h at room temperature and was monitored by TLC. The reaction mixture was diluted with cold water and DCM. The organic layer was separated, dried over $Na_2SO_4$ and distilled under reduced pressure. Then crude was purified by silica gel chromatography (5% MeOH/DCM) to give compound **7** (281 mg, 75%) as a white solid. $^1$H NMR (400 MHz, $CDCl_3$): δ 7.47- 7.45(m, 2H), 7.40- 7.38(m, 2H), 6.99- 6.97(m, 2H), 6.94- 6.92(m, 2H), 6.89- 6.88(m, 4H), 4.86(s, 2H), 3.95(t, 2H, $J$ = 6), 3.87(s, 2H), 3.58(t, 2H, $J$ = 6), 3.47(s, 4H), 3.40(t, 2H, $J$ = 6), 3.33- 3.31(m, 2H), 3.18(s, 6H), 2.73- 2.70(m, 2H), 2.59- 2.56(m, 2H), 1.82- 1.79(m, 2H), 1.72- 1.68(m, 2H), 1.46(s, 9H), 1.36- 1.34(m, 1H), 1.23- 1.21(m, 1H). $^{13}$C NMR (100 MHz, $CDCl_3$): δ 173.5, 170.9, 167.0, 160.7, 160.4, 159.4, 156.0, 154.6, 148.6, 142.8, 138.4, 134.3, 131.0, 129.2, 121.6, 120.4, 118.0, 115.7, 80.7, 67.8, 53.3, 46.2, 45.3, 41.8, 41.7, 39.5, 31.2, 28.8, 28.3, 26.5, 26.0, 20.5. HRMS (m/z): [M+H] + calcd. for $C_{40}H_{53}ClN_5O_7$, 750.3634; found: 750.3632.

**Preparation of (2R,4R)-1-((R)-2-(4-(4-(4-((4-(4-(4-(2-chloro-N-(4-(dimethylamino)benzyl) acetamido)phenoxy)phenoxy)butyl)amino)-4-oxobutanoyl)piperazin-1-yl)-4-oxobutanamido)-3,3-dimethylbutanoyl)-4-hydroxy-N-(4-(4-methylthiazol-5-yl)benzyl)pyrrolidine-2-carboxamide (R4VPL3-2):**

To the solution of **7** **7**(39 mg, 0.05 mmol) in 2mL DCM, TFA(2mL) was added dropwise and stirred the reaction for 1 h at room temperature. After 1 h, the reaction mixture was distilled under reduced pressure. The crude was diluted with dichloromethane and distilled under reduced pressure (3 times). Then to the solution of this crude product in DMF(5mL), VHL-acid (32 mg, 0.06 mmol), HATU (23 mg, 0.06 mmol) and DIPEA (26µL, 0.15 mmol) were added and continued the reaction for 2h at room temperature and was monitored by TLC. The reaction mixture was diluted with cold water and ethyl acetate. The organic layer was separated, dried over $Na_2SO_4$ and distilled under reduced pressure. Then crude was purified by preparative HPLC with a gradient of 0-60% ACN/$H_2O$ in 30 min system with a reverse phase C18 column to afford **R4VPL3-2** (29 mg, 50%) as a white solid. $^1H$ NMR (400 MHz, $CDCl_3$): δ 8.87(s, 1H), 7.52- 7.40(m, 1H), 7.28- 7.26(m, 6H), 7.18(s, 2H), 6.91- 6.89(m, 2H), 6.86- 6.83(m, 2H), 6.81- 6.78(m, 4H), 6.21- 6.15(m, 1H), 4.76(s, 2H), 4.46- 4.41(m, 2H), 4.31- 4.25(m, 1H), 3.95- 3.93(m, 1H), 3.86(t, 2H, $J$ = 6), 3.78(s, 2H), 3.54- 3.51(m, 2H), 3.47- 3.39(m, 4H), 3.21(t, 2H, $J$ = 6), 3.05(s, 6H), 2.71- 2.59(m, 2H), 2.55- 2.45(m, 7H), 2.33- 2.27(m, 1H), 2.10(s, 1H), 1.72- 1.70(m, 2H), 1.62- 1.60(m, 2H), 0.88(s, 9H). $^{13}C$ NMR (100 MHz, $CDCl_3$): δ 172.8, 172.7, 171.7, 171.2, 171.0, 170.9, 170.8, 166.8, 161.2, 160.8, 159.2, 155.9, 151.6, 148.7, 143.8, 139.0, 136.5, 134.4, 130.9, 129.4, 129.3, 128.2, 121.6, 119.4, 117.9, 117.2, 115.7, 114.3, 70.1, 67.8, 58.8, 58.0, 56.9, 53.3, 45.4, 44.9, 43.1, 41.9, 41.6, 39.3, 36.6, 35.1, 31.1, 30.0, 26.4, 14.9. HRMS (m/z): [M+H] + calcd. for $C_{61}H_{77}ClN_9O_{10}S$, 1162.5203; found: 1162.5209.

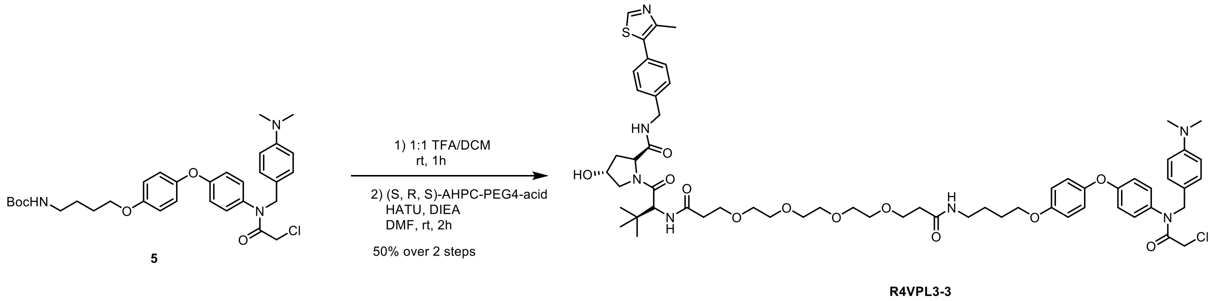


**Preparation of N1-(4-(4-(4-(2-chloro-N-(4-(dimethylamino) benzyl) acetamido) phenoxy) phenoxy) butyl)-N16-( (R)-1-( (2R, 4R)-4-hydroxy-2-( (4- (4-methylthiazol-5-yl) benzyl) carbamoyl) pyrrolidin-1-yl) -3, 3-dimethyl-1-oxobutan-2-yl) -4, 7, 10, 13 tetraoxahexadecanediamide (R4VPL3-3):**

To the solution of **5** (35 mg, 0.06 mmol) in 2mL DCM, TFA(2mL) was added dropwise and stirred the reaction for 1 h at room temperature. After 1 h, the reaction mixture was distilled under reduced pressure. The crude was diluted with DCM and distilled under reduced pressure (3 times). Then to the solution of this crude product in DMF(5mL), (S, R, S)-AHPC-PEG 4-acid (49 mg, 0.07 mmol), HATU (27 mg, 0.07 mmol) and DIPEA (39 mg, 0.3 mmol) were added and continued the reaction for 2h at room temperature and was monitored by TLC. The reaction mixture was diluted with cold water and ethyl acetate. The organic layer was separated, dried over $Na_2SO_4$ and distilled under reduced pressure. Then crude was purified by preparative HPLC with a gradient of 0-60% ACN/$H_2O$ in 30 min system with a reverse phase C18 column to afford **R4VPL3-3** (35 mg, 50%) as a white solid. $^1H$ NMR (400 MHz, $CDCl_3$): δ 9.05(s, 1H), 7.68- 7.61(m, 2H), 7.59- 7.57(m, 1H), 7.52- 7.47(m, 3H), 7.38- 7.36(m,

6H), 7.19- 7.14(1H), 6.99- 6.87(m, 5H), 4.85(s, 2H), 4.70- 4.59(m, 1H), 4.57- 4.51(m, 3H), 3.94(t, 2H, *J* = 6), 3.86(s, 2H), 3.73- 3.70(m, 4H), 3.66- 3.61(m, 13H), 3.32- 3.27(m, 2H), 3.16(s, 6H), 2.55- 2.52(m, 7H), 2.43- 2.36(m, 1H), 2.24- 2.22(m, 1H), 1.79(t, 2H, *J* = 6), 1.68( t, 2H, *J* = 6), 0.96- 0.93(m, 9H). $^{13}C$ NMR (100 MHz, $CDCl_3$): δ 172.9, 172.7, 171.6, 171.2, 166.9, 160.8, 160.4, 159.3, 155.9, 152.0, 148.6, 143.1, 139.2, 137.8, 134.3, 132.6, 132.2, 132.1, 131.0, 129.4, 129.2, 128.8, 128.7, 128.3, 121.6, 120.1, 118.0, 115.7, 70.1, 70.0, 69.9, 67.8, 67.2, 66.9, 58.9, 58.0, 57.0, 53.3, 45.9, 43.1, 41.8, 39.3, 36.4, 36.3, 36.1, 35.1, 26.5, 26.4, 26.0, 14.6. HRMS (m/z): [M+H] + calcd. for $C_{61}H_{81}ClN_7O_{12}S$, 1170.5352; found: 1170.5333.

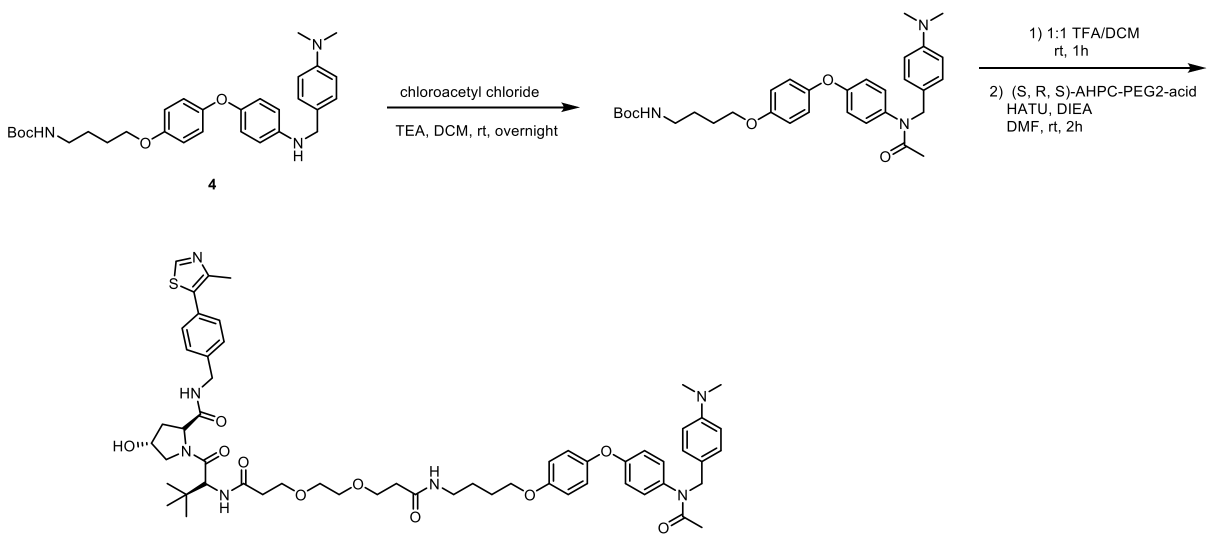


PEG 2 control PROTAC

**Preparation of inactive R4VP L3-1:** $^1H$ NMR (400 MHz, $CDCl_3$): δ 9.19(s, 1H), 7.56- 7.54(m, 2H), 7.51- 7.46(m, 5H), 7.37- 7.36(m, 4H), 7.07- 7.05(m, 2H), 6.97-6.95(m, 3H), 4.95(s, 2H), 4.77- 4.70(m, 2H), 4.65- 4.63(m, 2H), 4.47- 4.41(m, 1H), 4.22- 4.20(m, 1H), 4.03(t, 2H, *J* = 6), 3.81- 3.76(m, 4H), 3.75- 3.69(m, 4H), 3.38- 3.37(m, 2H), 3.27(s, 6H), 2.65-2.42(m, 8H), 2.33- 2.27(m, 1H), 2.01(s, 3H), 1.89- 1.85(m, 2H), 1.77- 1.73(m, 2H), 1.08(m, 9H). $^{13}C$ NMR (100 MHz, $CDCl_3$): δ 173.1, 172.9, 171.9, 171.6, 171.3, 160.8, 160.4, 158.7, 155.7, 152.2, 149.0, 145.3, 142.5, 139.4, 139.0, 136.1, 130.9, 129.4, 129.0, 128.3, 121.5, 120.3, 118.0, 116.9, 115.7, 114.0, 70.2, 70.0, 67.9, 67.1, 66.8, 59.1, 58.1, 57.1, 52.6, 46.2, 43.1, 39.4, 36.7, 36.3, 35.9, 35.1, 26.3, 22.3, 14.3.

**Preparation of Biotin-R4VPL3-1:**

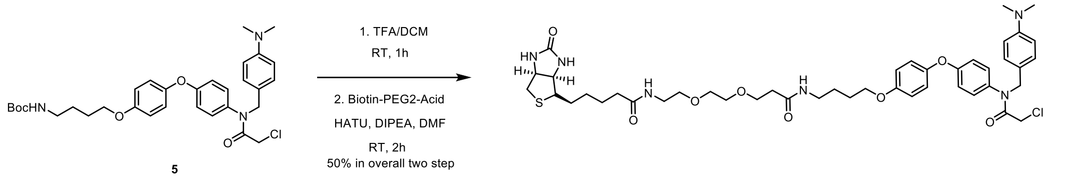

### R4VP used in Fig 1C, F

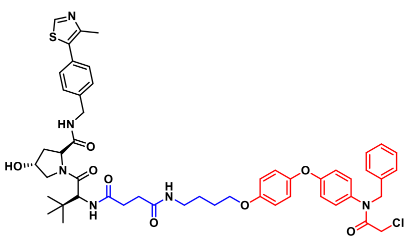

R4VP

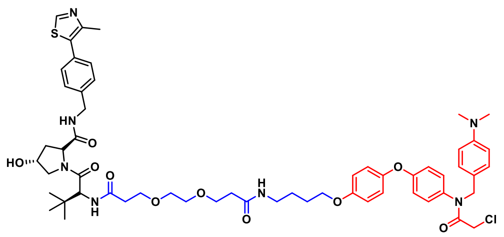

R4VP L3-1

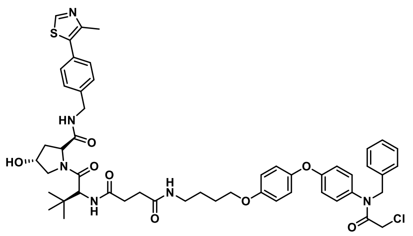

R4VP

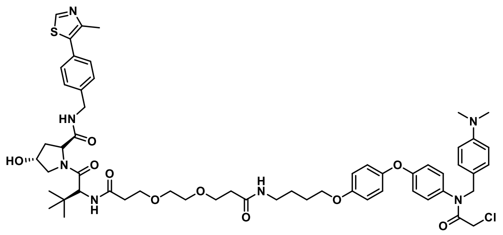

R4VP L3-1

Supp Fig. 2A a - Recruiters structure

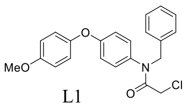

L1

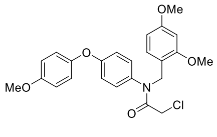

L2

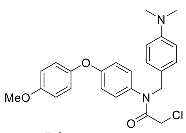

L3

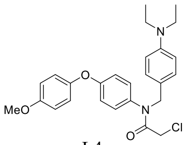

L4

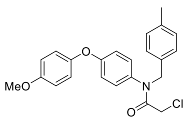

L5

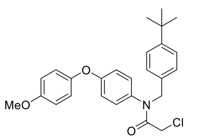

L6

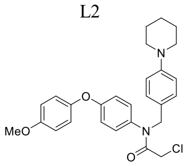

L7

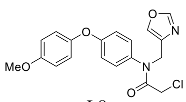

L8

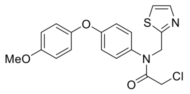

L9

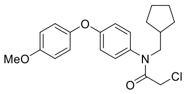

L10

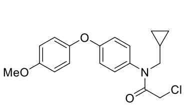

L11

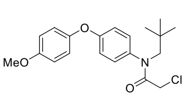

L12

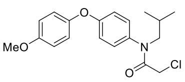

L13

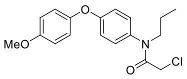

L14

**Preparation of Biotin-R4VPL3-1:** To a solution of **5** (100mg, 0.17 mmol) in DCM(5mL), TFA (5 mL) was added dropwise and stirred the reaction for 1 h at room temperature. After 1 h, the reaction mixture was distilled under reduced pressure. The crude was diluted with dichloromethane and distilled under reduced pressure (3 times). Then to the solution of this crude product in DMF (5 mL), **Biotin-2PEG- acid** (70mg, 0.17 mmol), HATU (76 mg, 0.20 mmol) and DIEA (148 µL, 0.85mmol) were added and continued the reaction for 2h at room. Then little amount of crude was purified by analytical HPLC in ACN/$H_2O$ system with a reverse phase C18 column followed by preparative to afford the **Biotin attached R4VP L3-1**. 1H NMR (400 MHz, CDCl3): δ 7.32-7.327(m, 3H), 7.02-6.95(m, 2H), 6.92-6.84(m, 5H), 6.54-6.43(m, 2H), 4.84(s, 2H), 4.56-4.53(m,1H), 4.39-4.36(m, 1H), 3.97(t, 2H, *J* = 4), 3.85(s, 2H), 3.77(t, 2H, *J* = 4),3.65-3.51(m, 4H), 3.47-3.39(m, 2H), 3.36-3.30(m, 2H), 3.21-3.15(m, 1H), 3.12-3.08(m, 4H), 2.98-2.90(m, 2H), 2.78-2.72(m, 1H), 2.49(t, 2H, *J* = 4)2.29-2.21(m, 2H), 1.85-1.77(m, 2H), 1.76-1.62(m, 4H), 1.49-1.39(m, 3H), 1.25(s, 2H), 1.10(s, 2H). HRMS (m/z): calcd. for $C_{44}H_{60}ClN_6O_8S$, 867.3882. found:867.3857

**$^1$H NMR of Amine 1:**

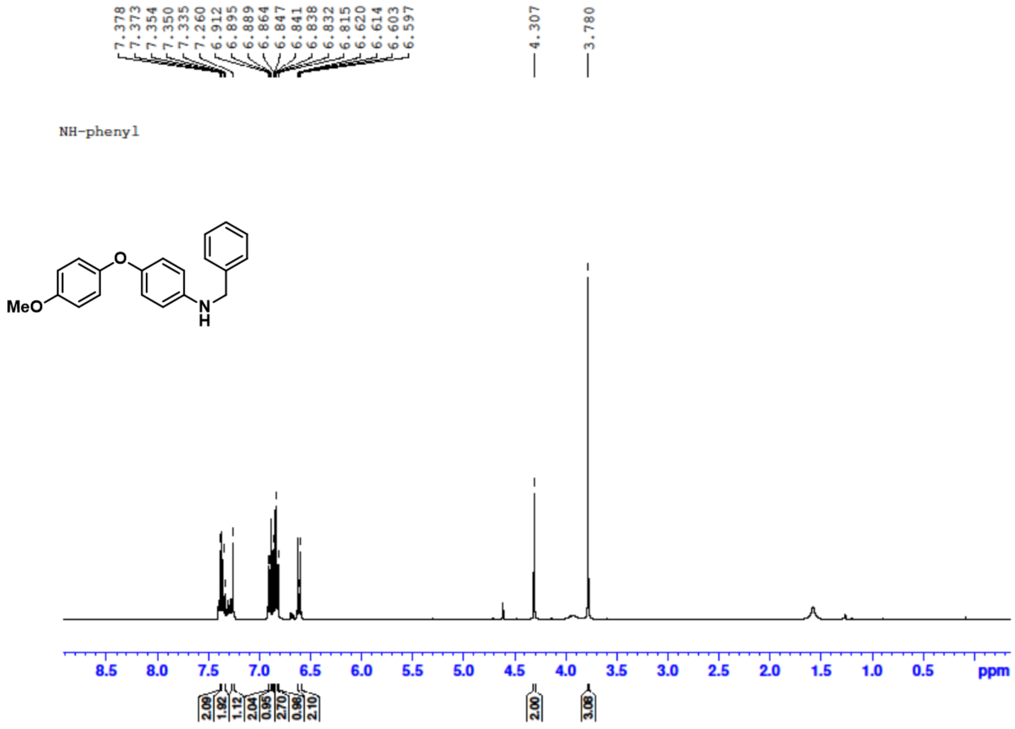

**[13]C NMR of Amine 1:**

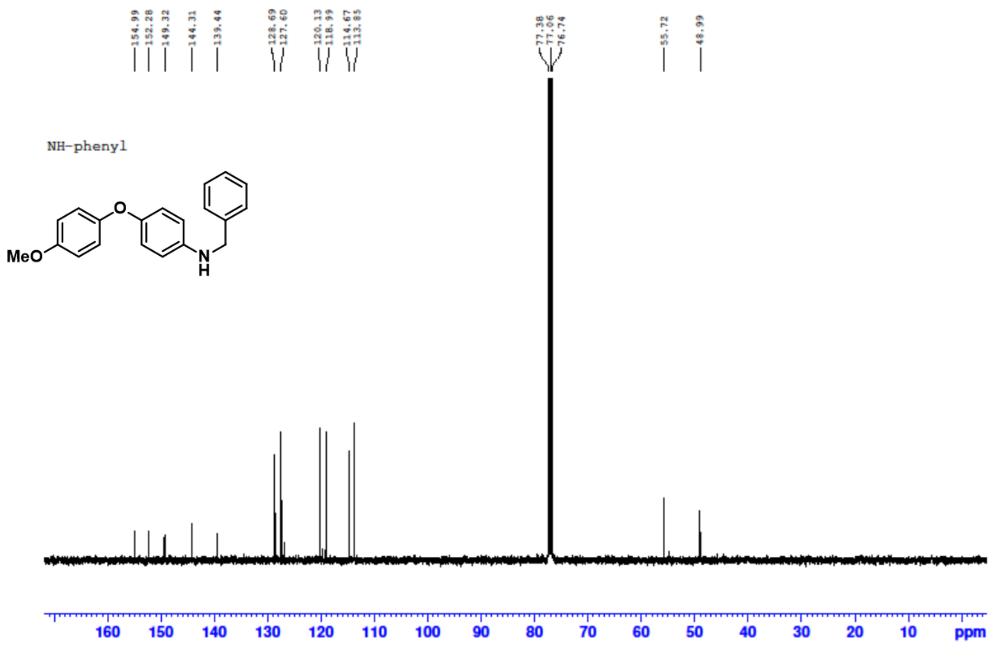


**[1]H NMR of Ligand 1:**

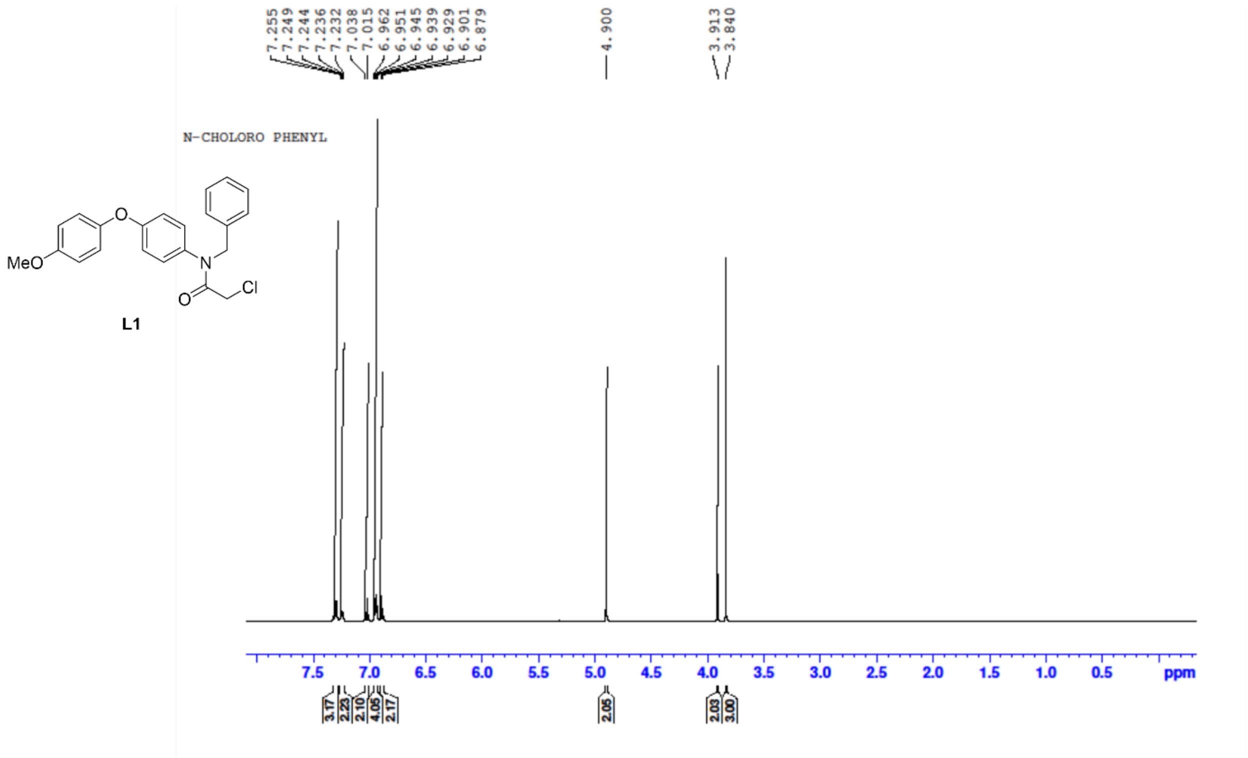

#### $^{13}$C NMR of Ligand 1:

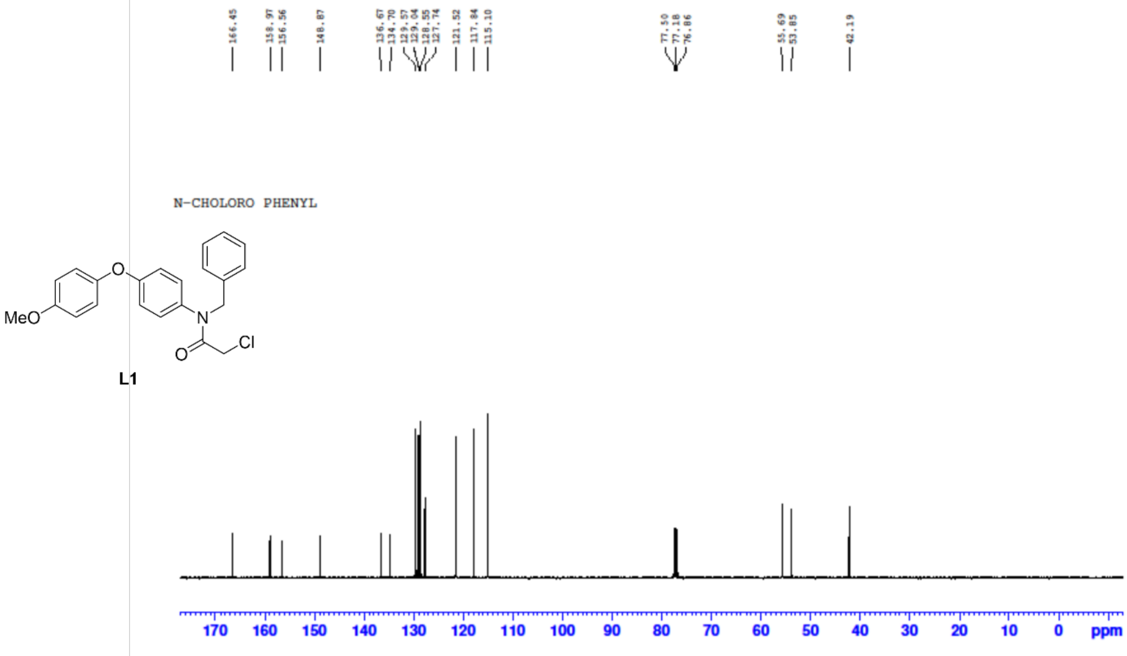


#### $^{1}$H NMR of Amine 2:

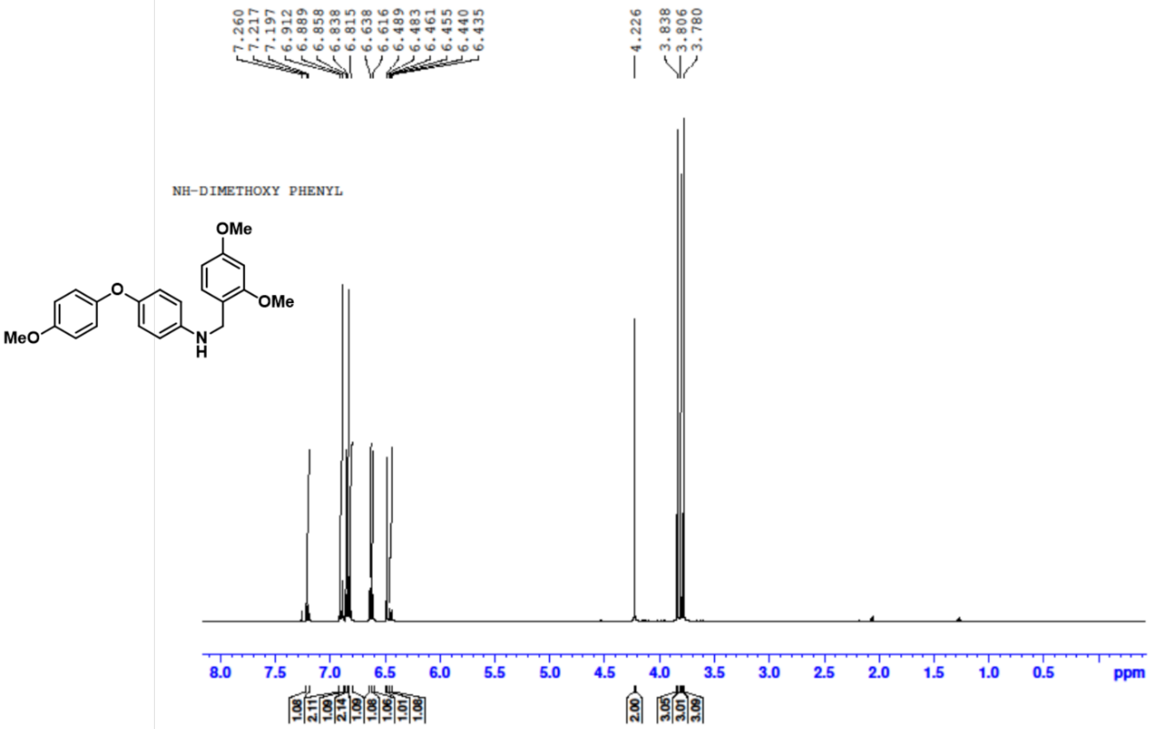

[13]C NMR of Amine 2:

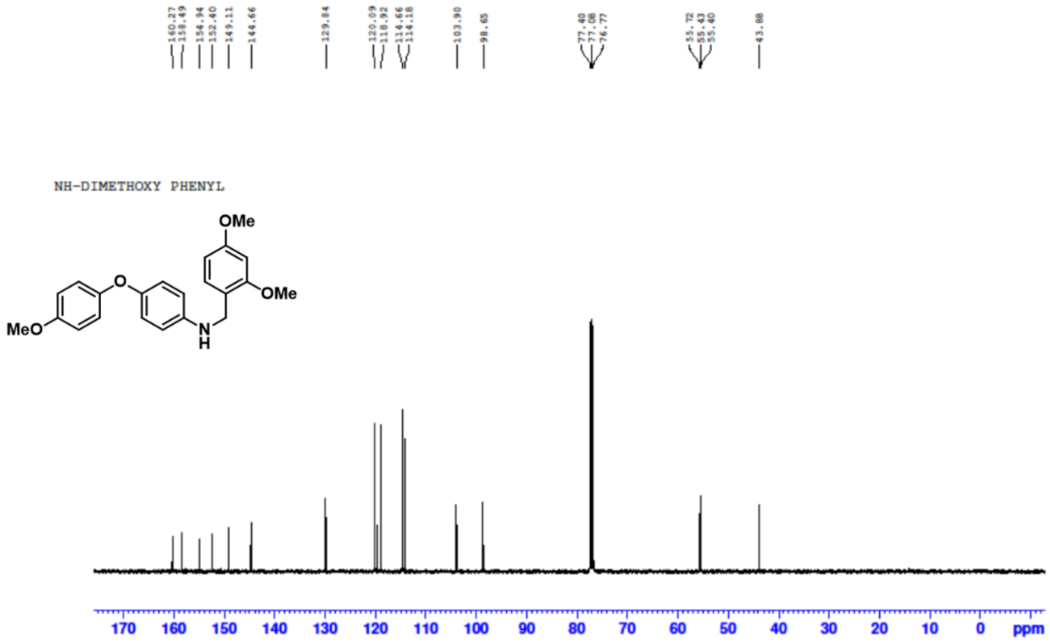


[1]H NMR of Ligand 2:

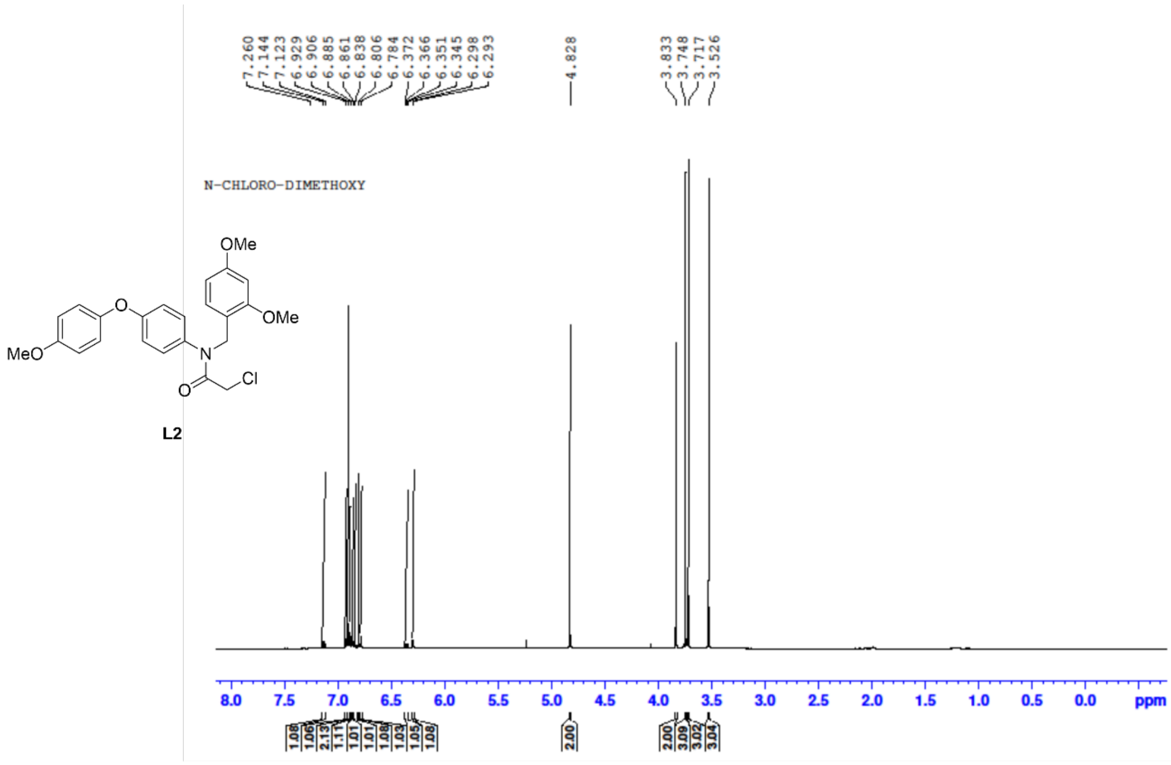

#### [13]C NMR of Ligand 2:

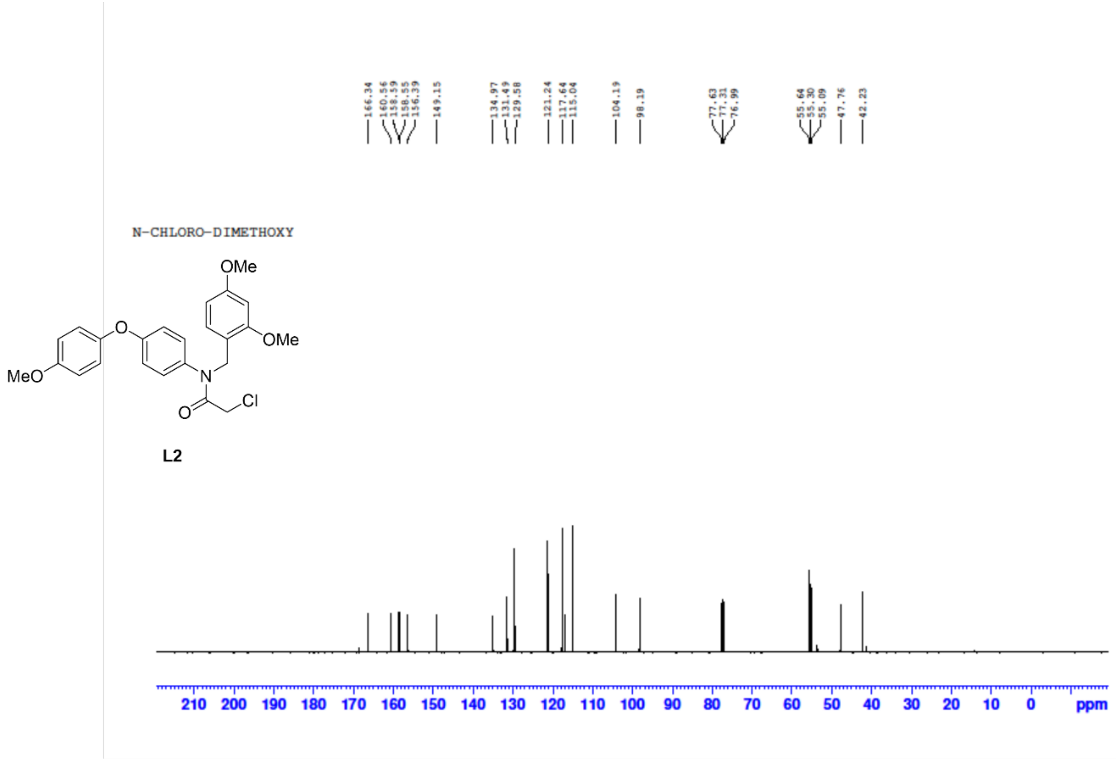


#### [1]H NMR of Amine 3:

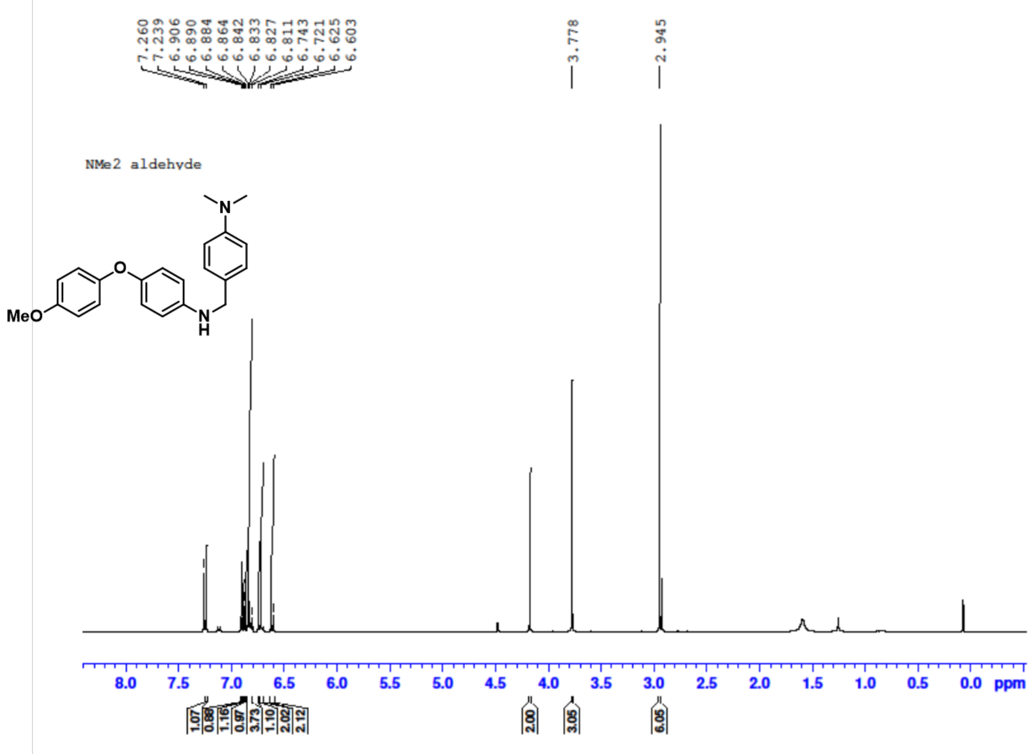

**$^{13}$C NMR of Amine 3:**

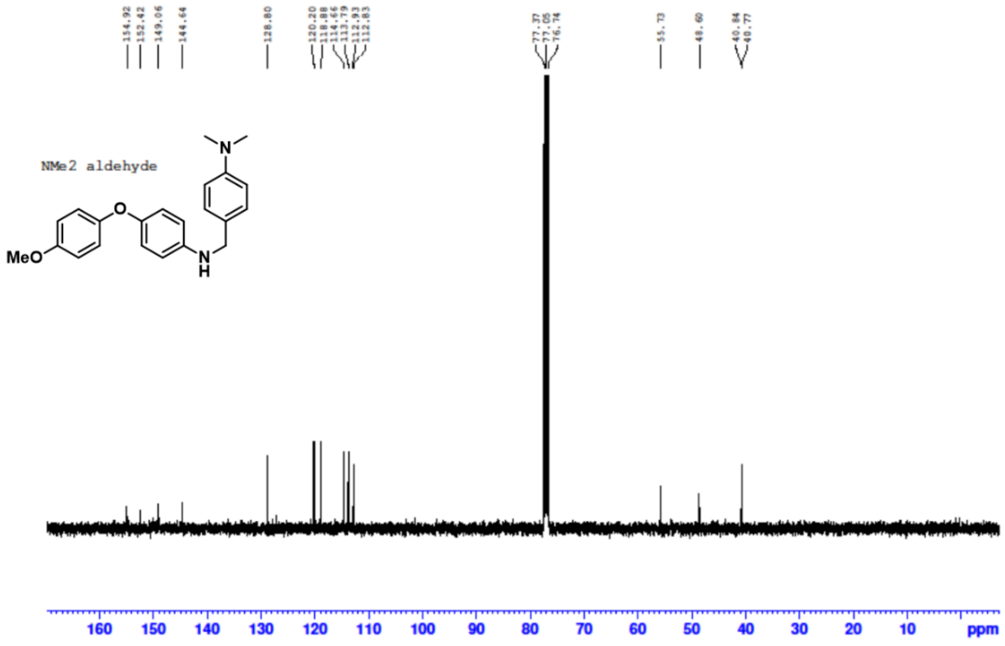


**$^{1}$H NMR of Ligand 3:**

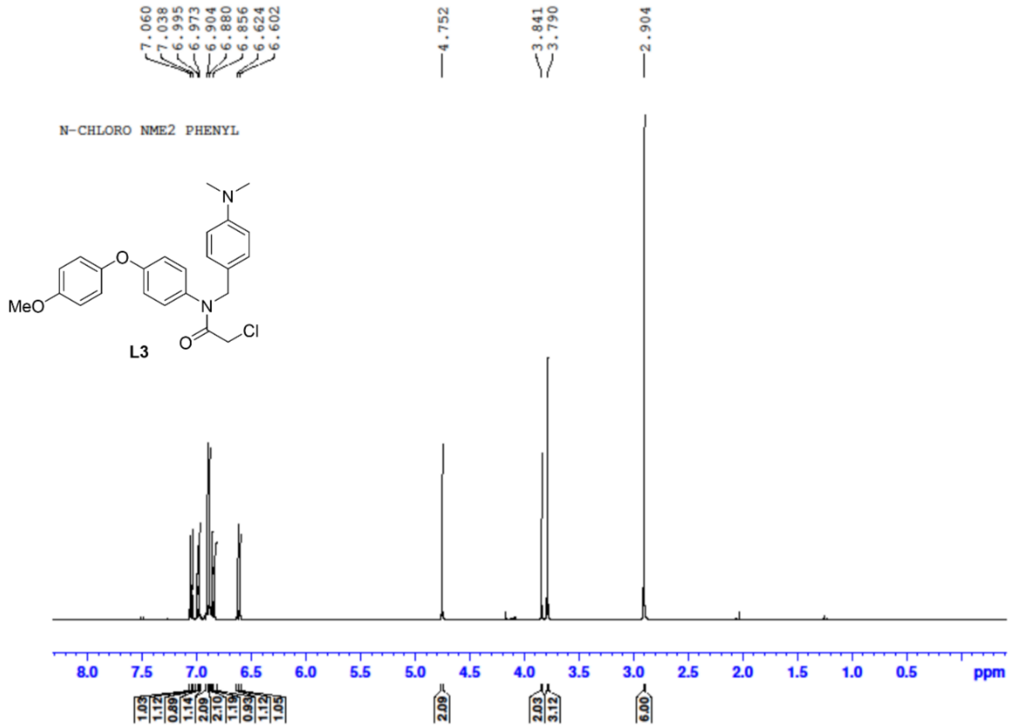

$^{13}$C NMR of Ligand 3:

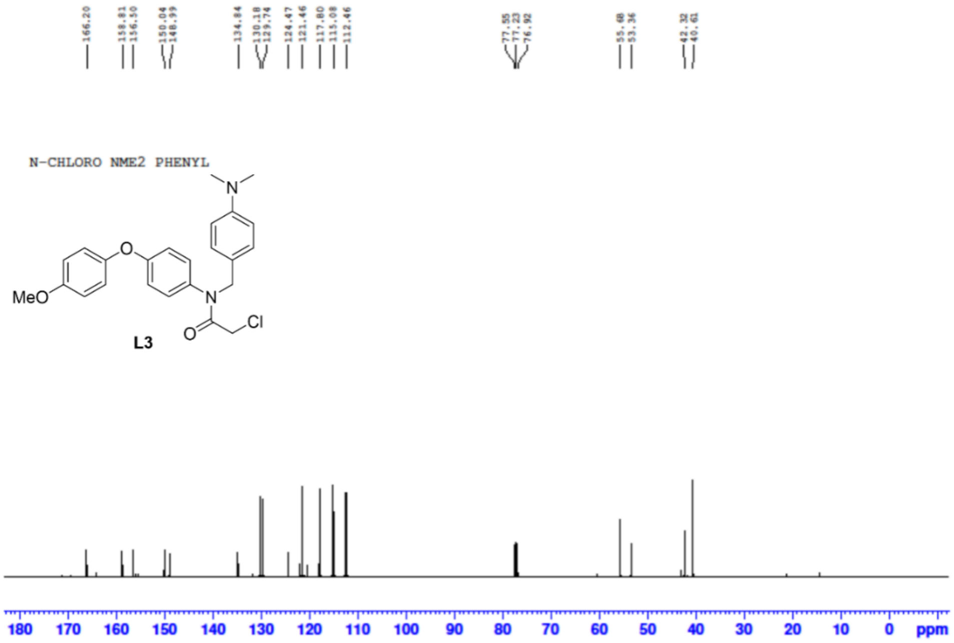


$^{1}$H NMR of Amine 4:

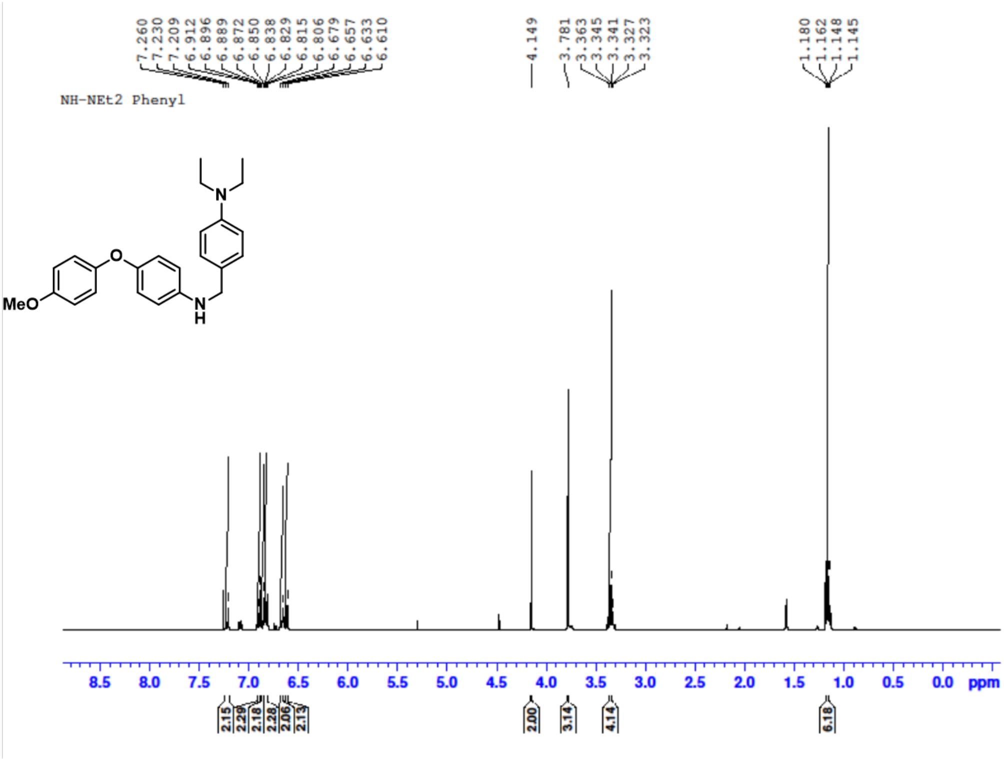

#### [13]C NMR of Amine 4:

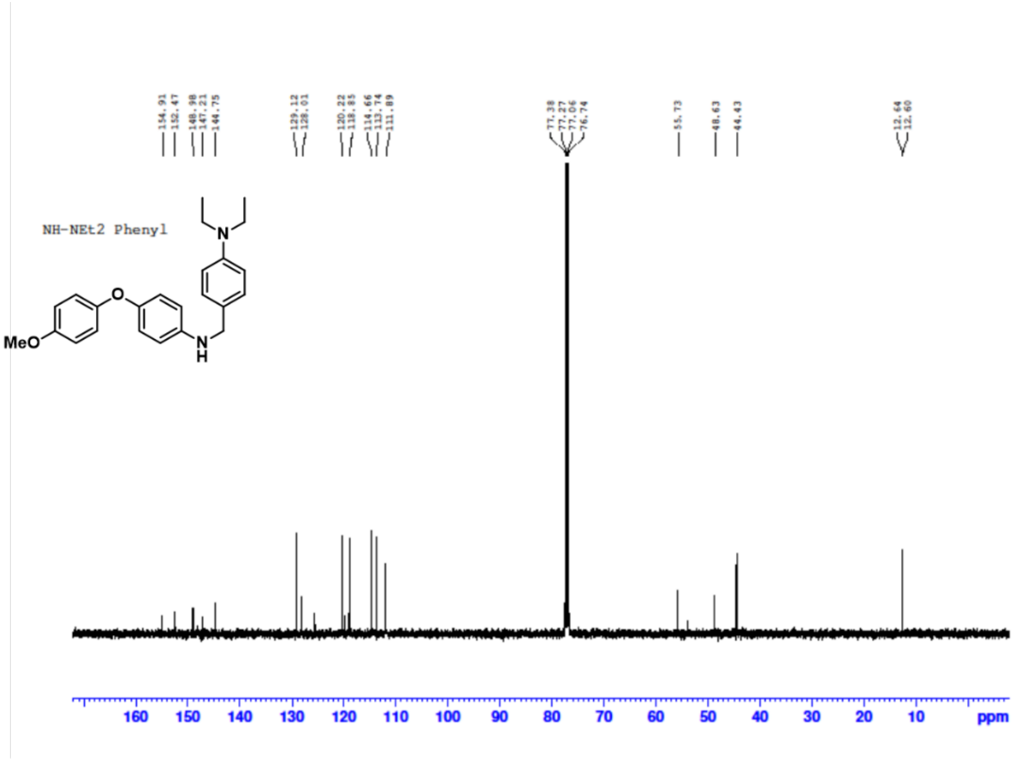


#### [1]H NMR of Ligand 4:

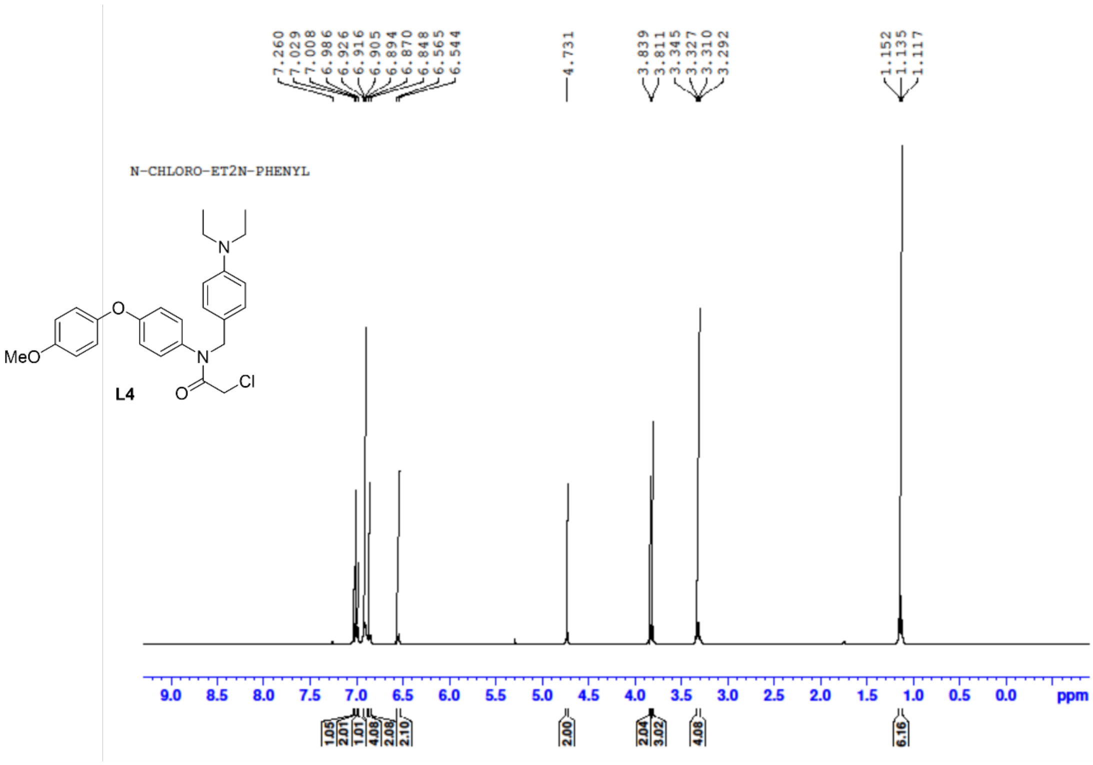

#### $^{13}$C NMR of Ligand 4:

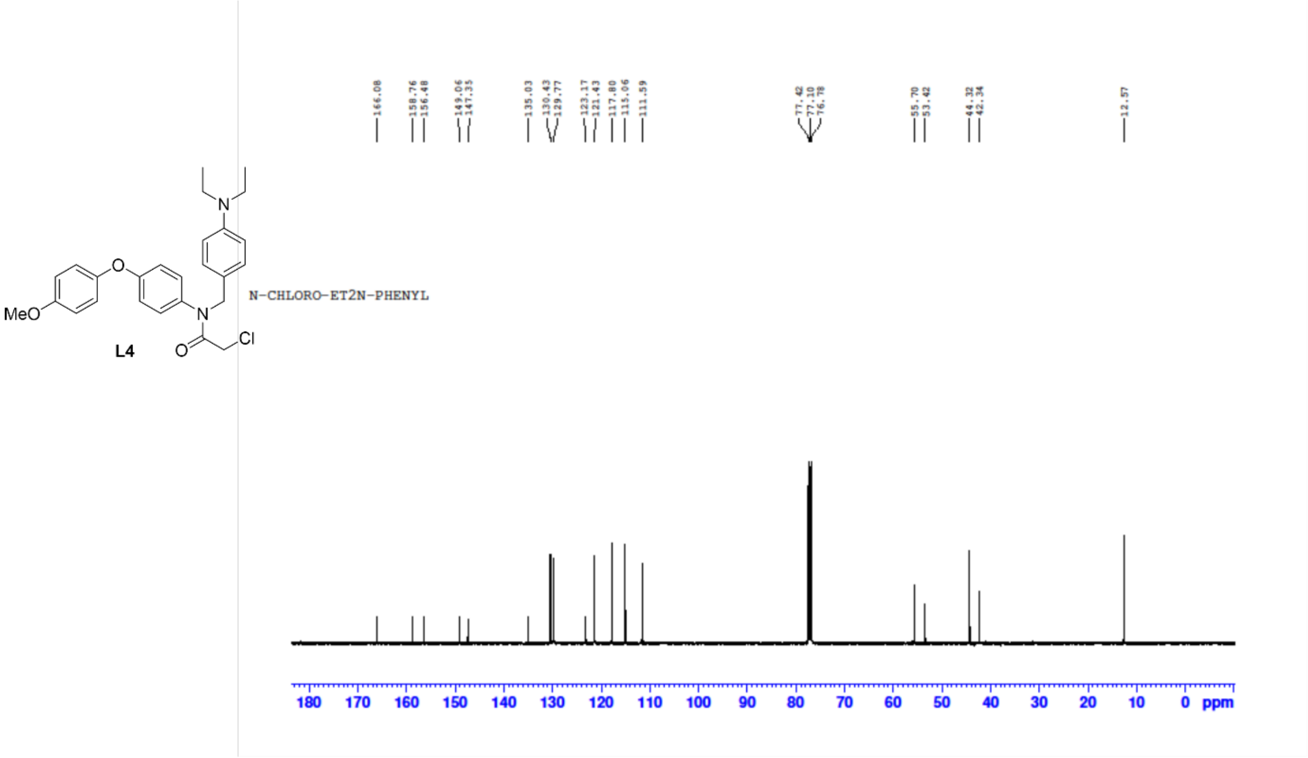


#### $^{1}$H NMR of Amine 5:

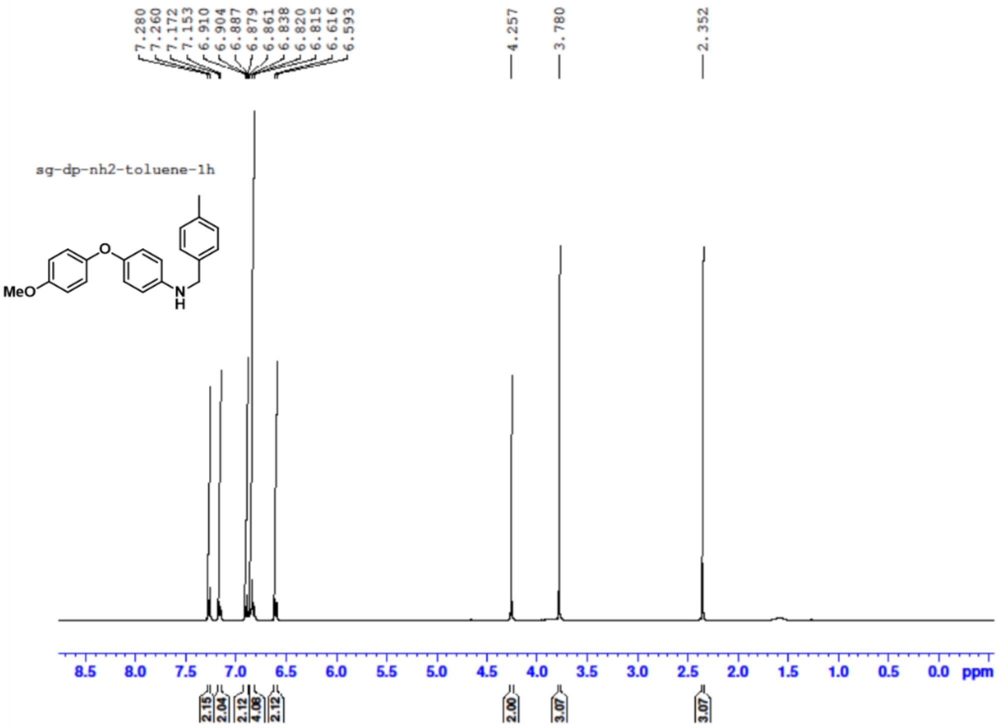

#### $^{13}$C NMR of Amine 5:

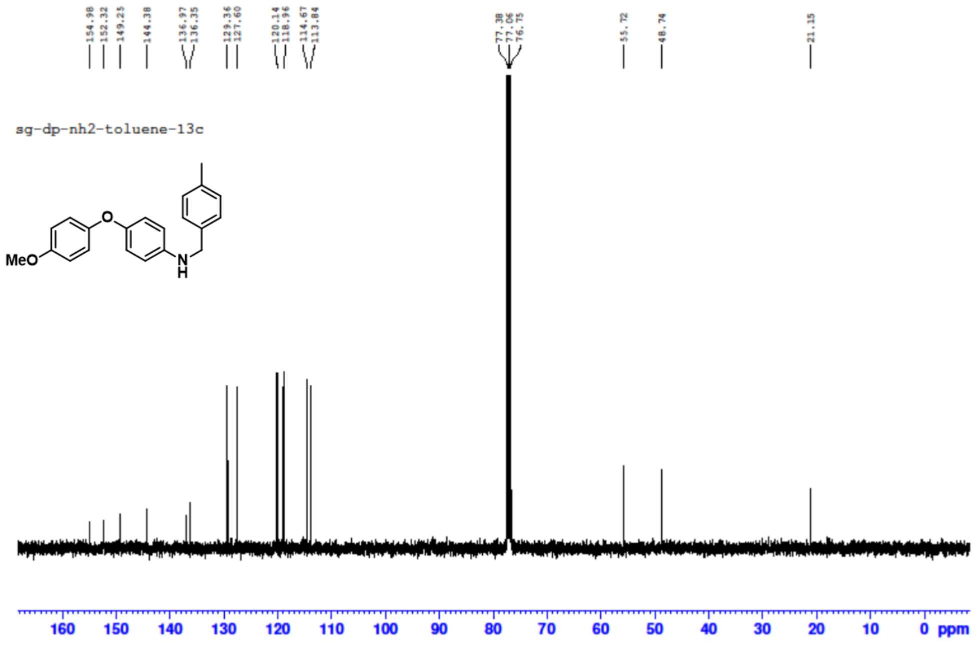


#### $^{1}$H NMR of Ligand 5:

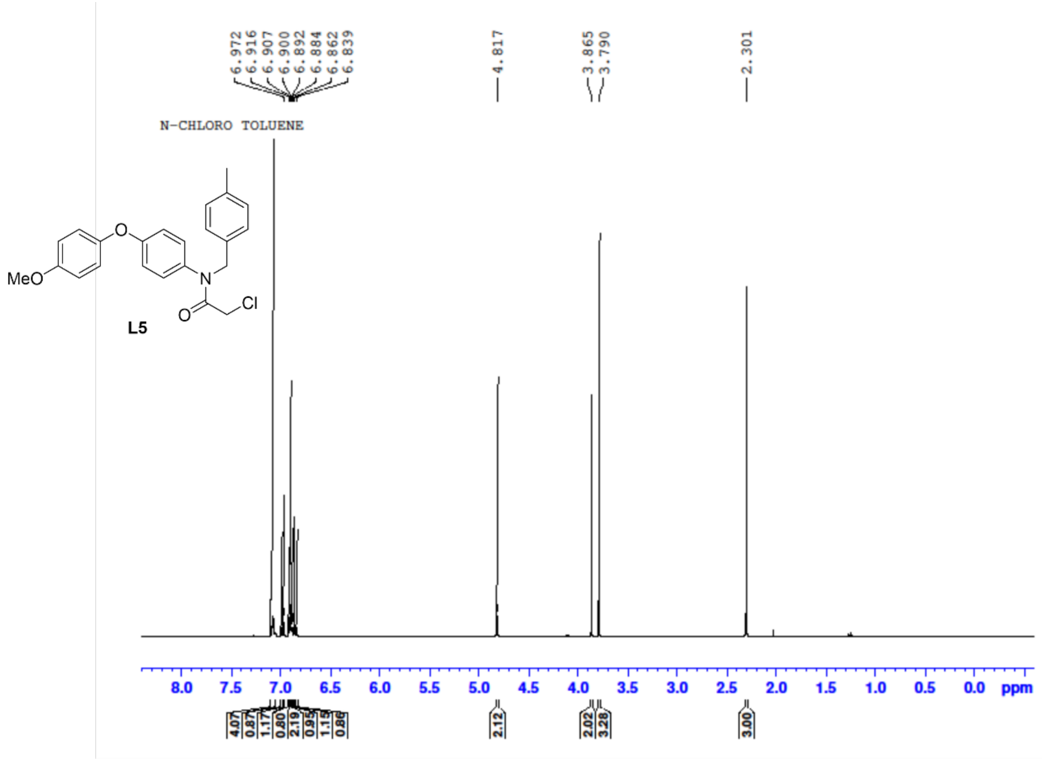

#### $^{13}$C NMR of Ligand 5:

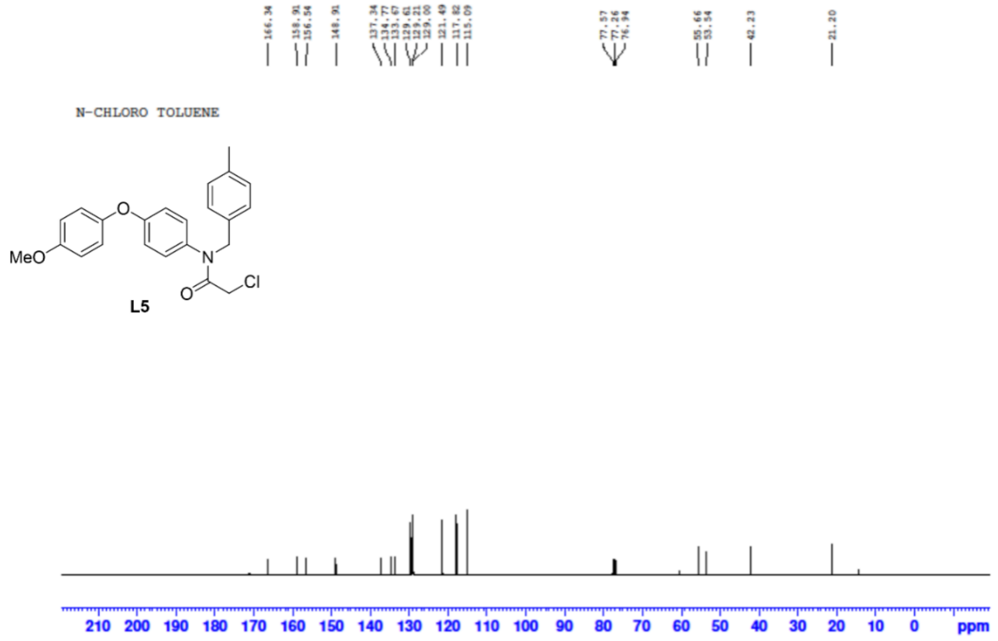


#### $^{1}$H NMR of Amine 6:

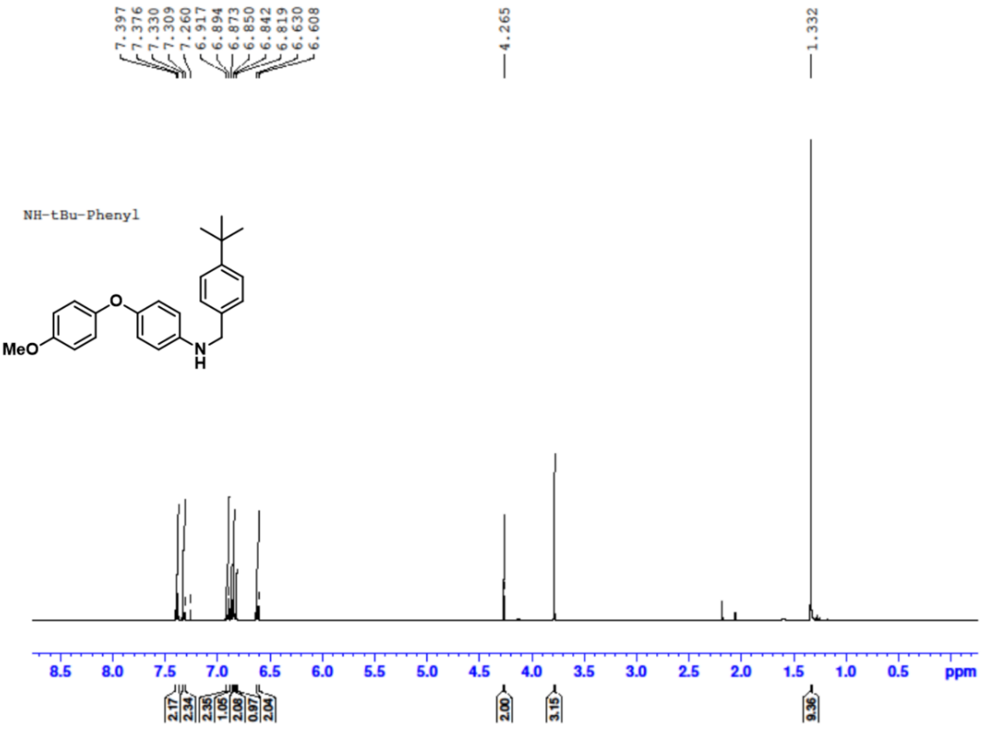

#### [13]C NMR of Amine 6:

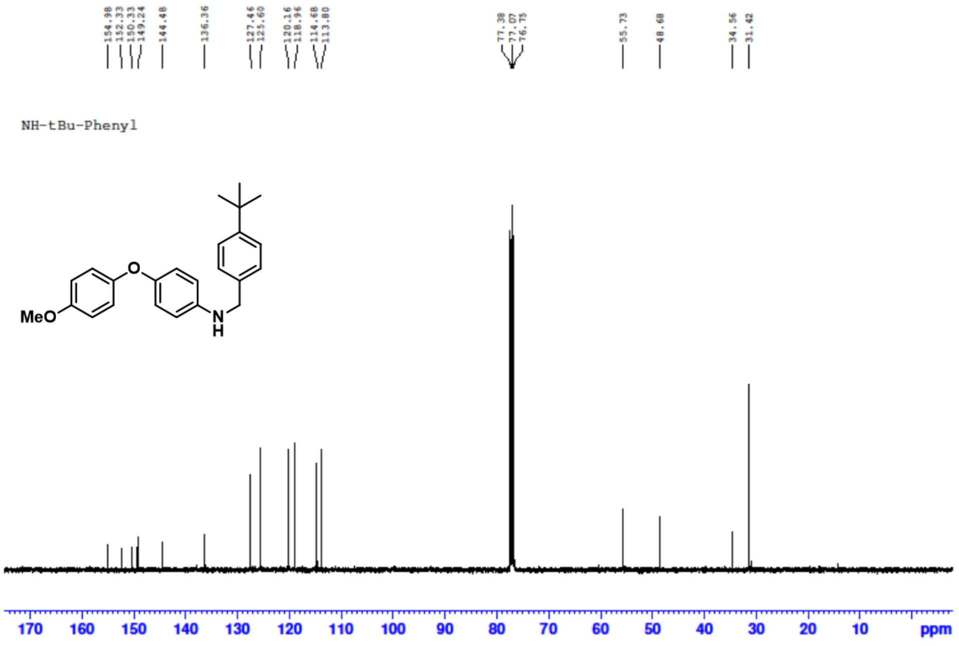


#### [1]H NMR of Ligand 6:

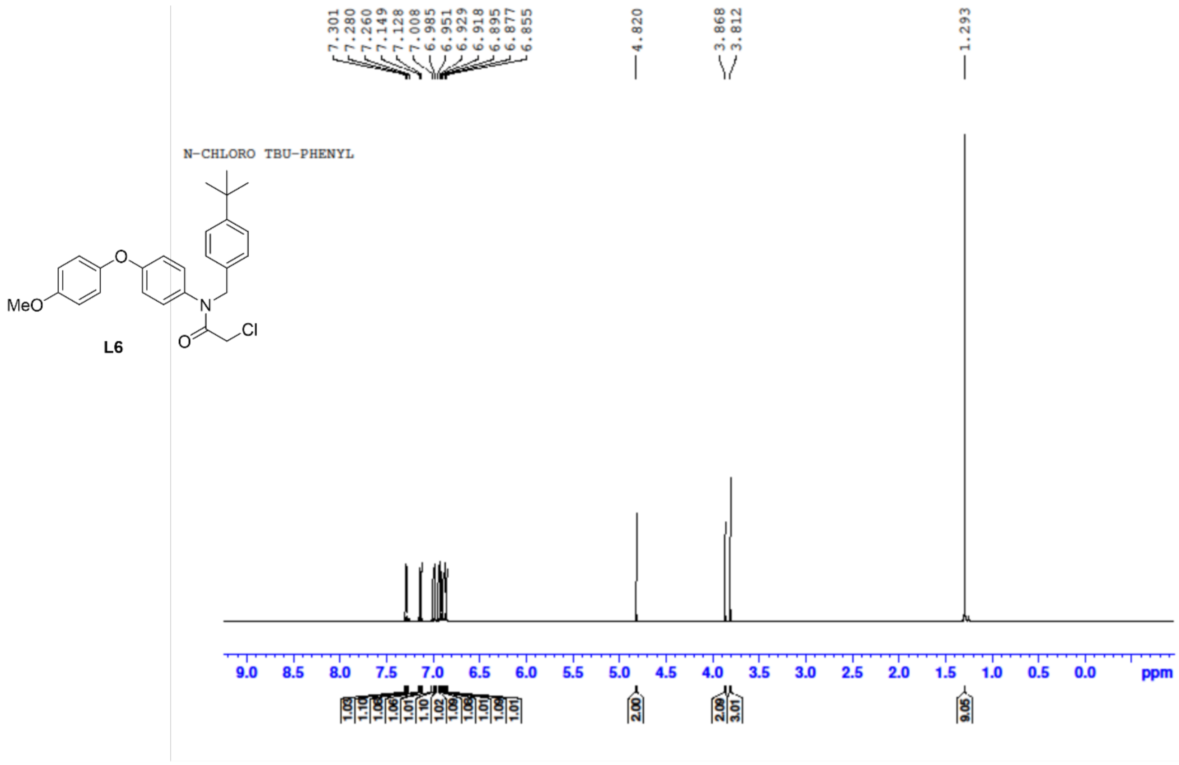

#### $^{13}$C NMR of Ligand 6:

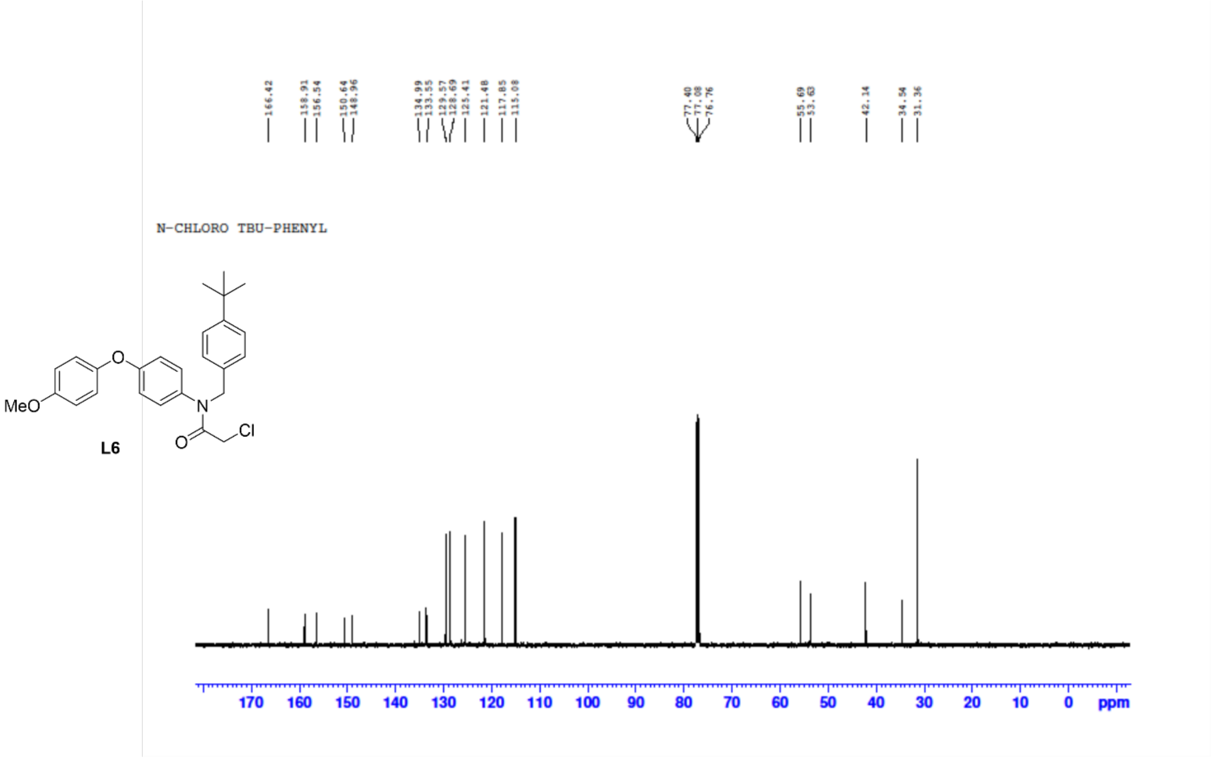


#### $^{1}$H NMR of Amine 7:

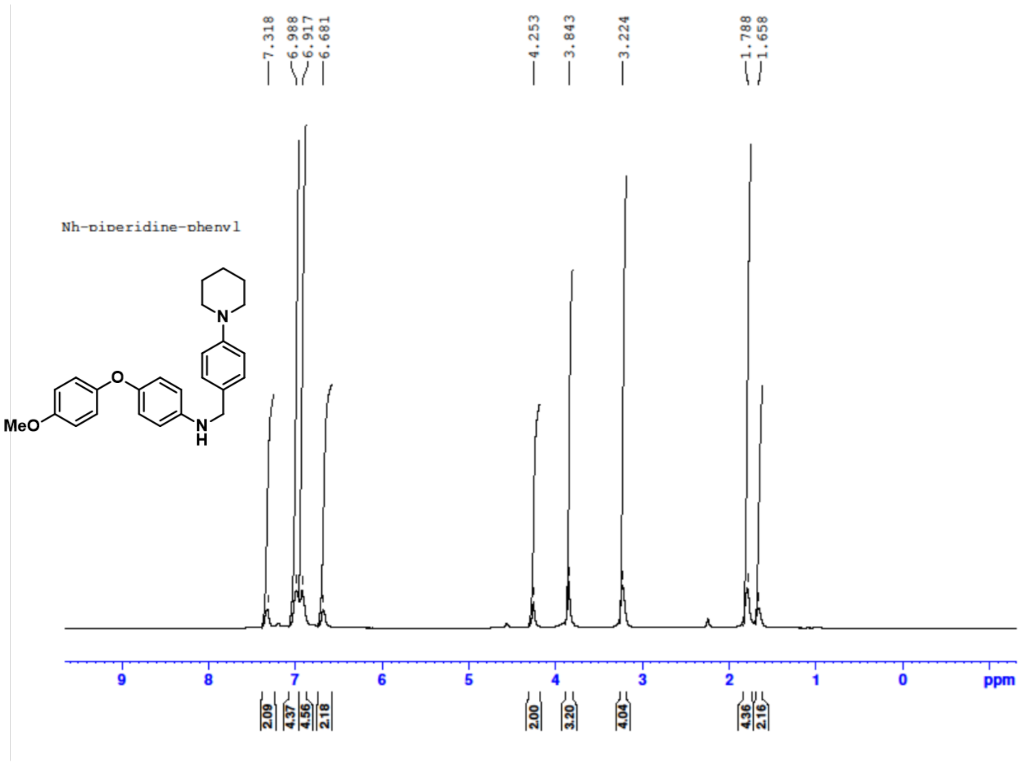

#### [13]C NMR of Amine 7:

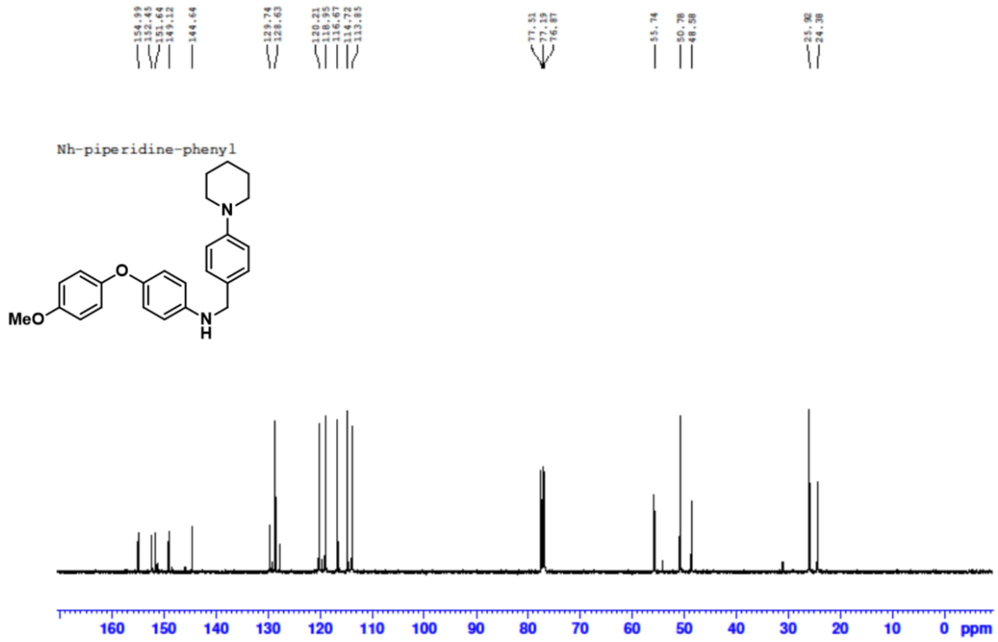


#### [1]H NMR of Ligand 7:

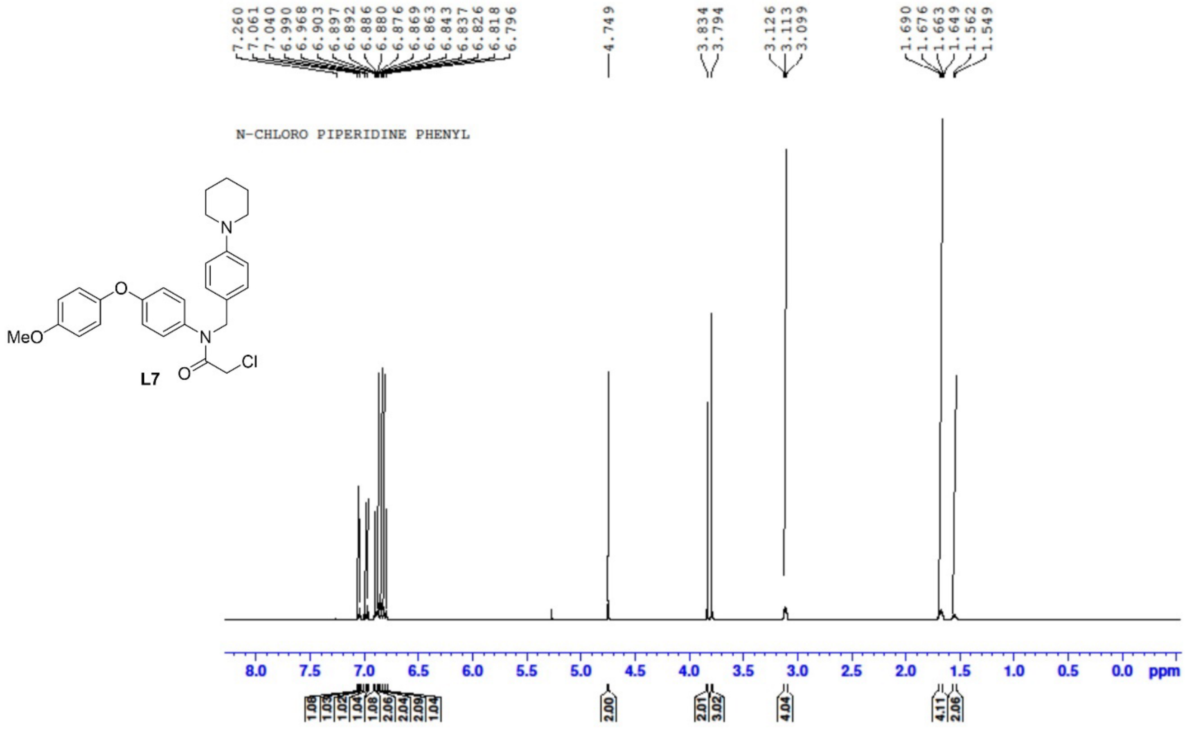

#### [13]C NMR of Ligand 7:

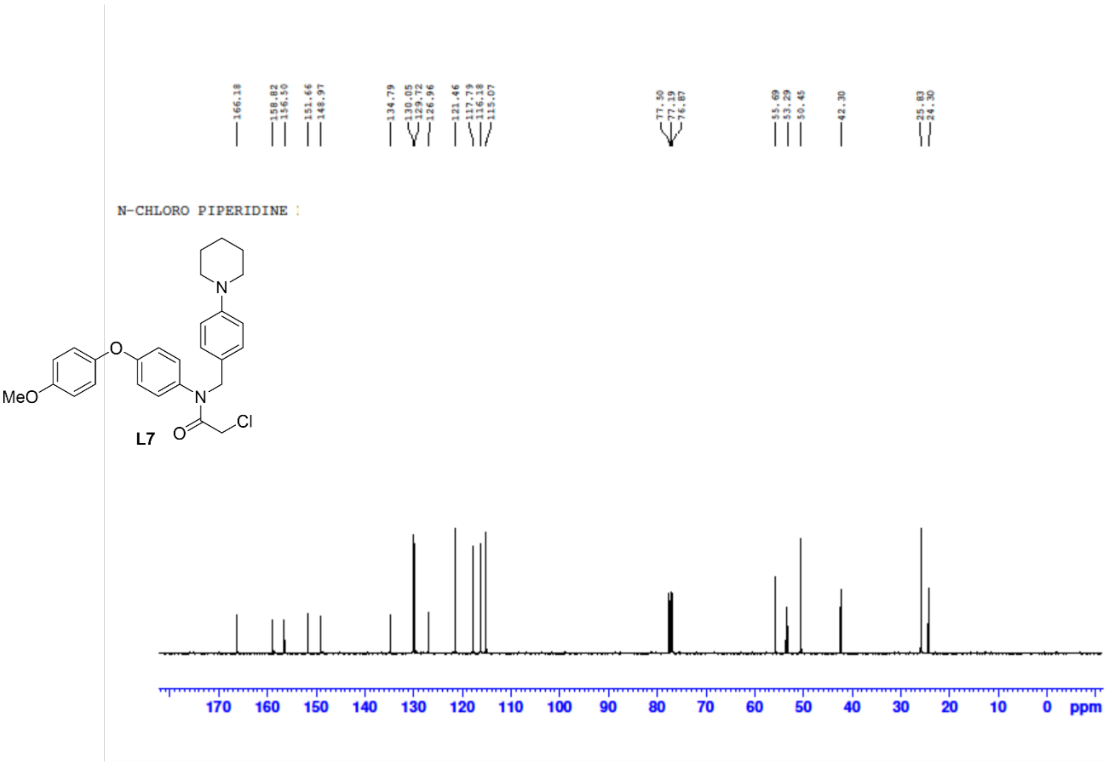


#### [1]H NMR of Amine 8:

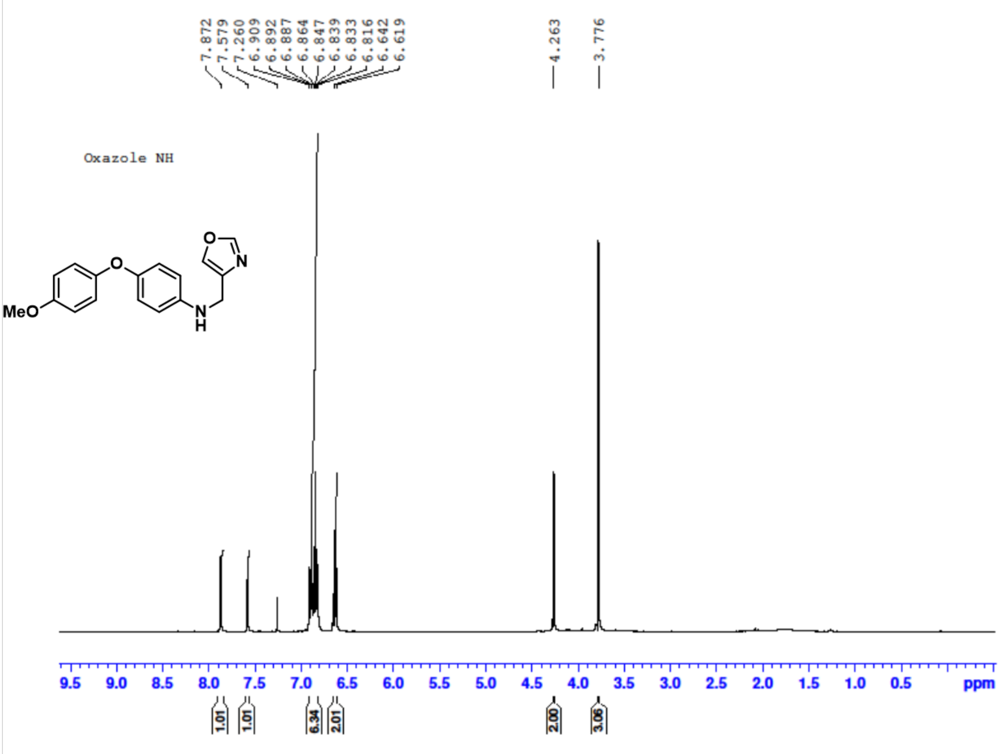

$^{13}$C NMR of Amine 8:

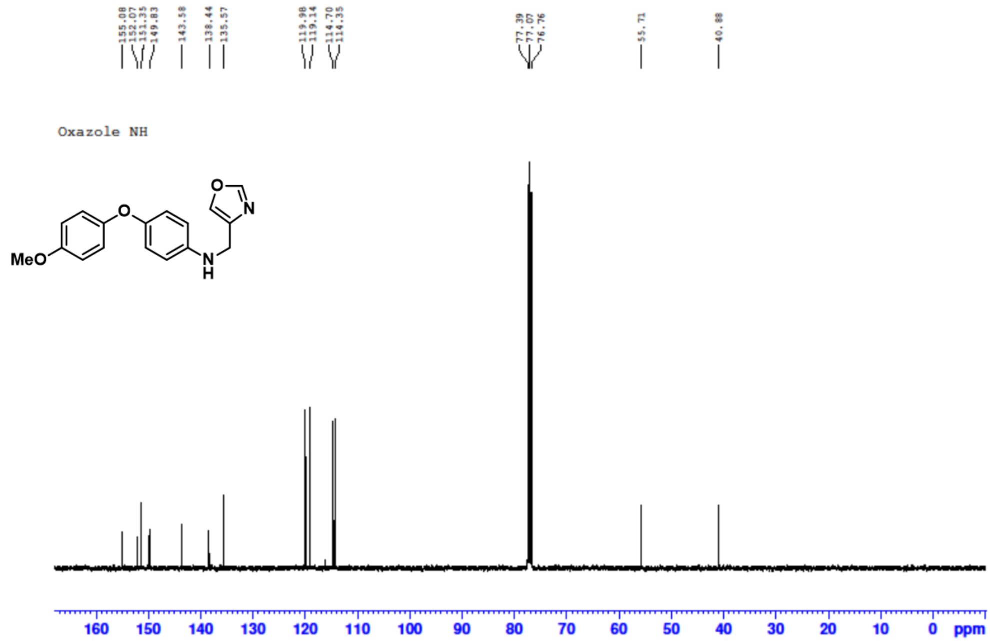


$^{1}$H NMR of Ligand 8:

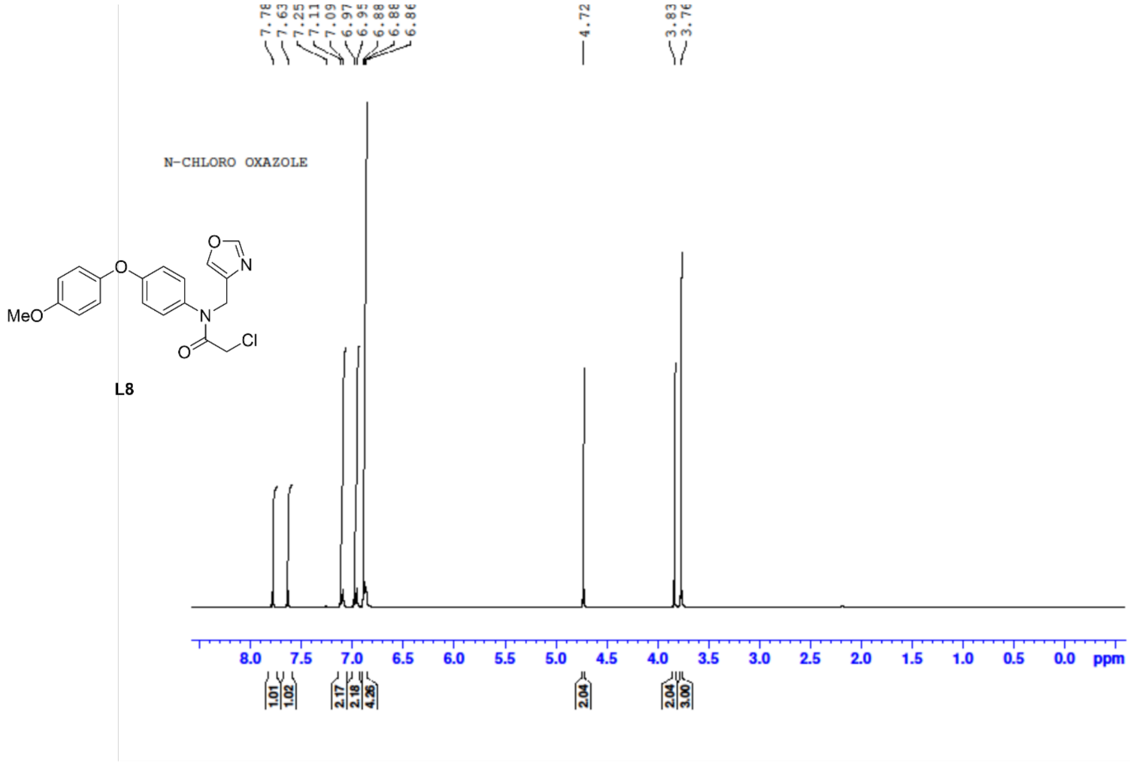

#### [13]C NMR of Ligand 8:

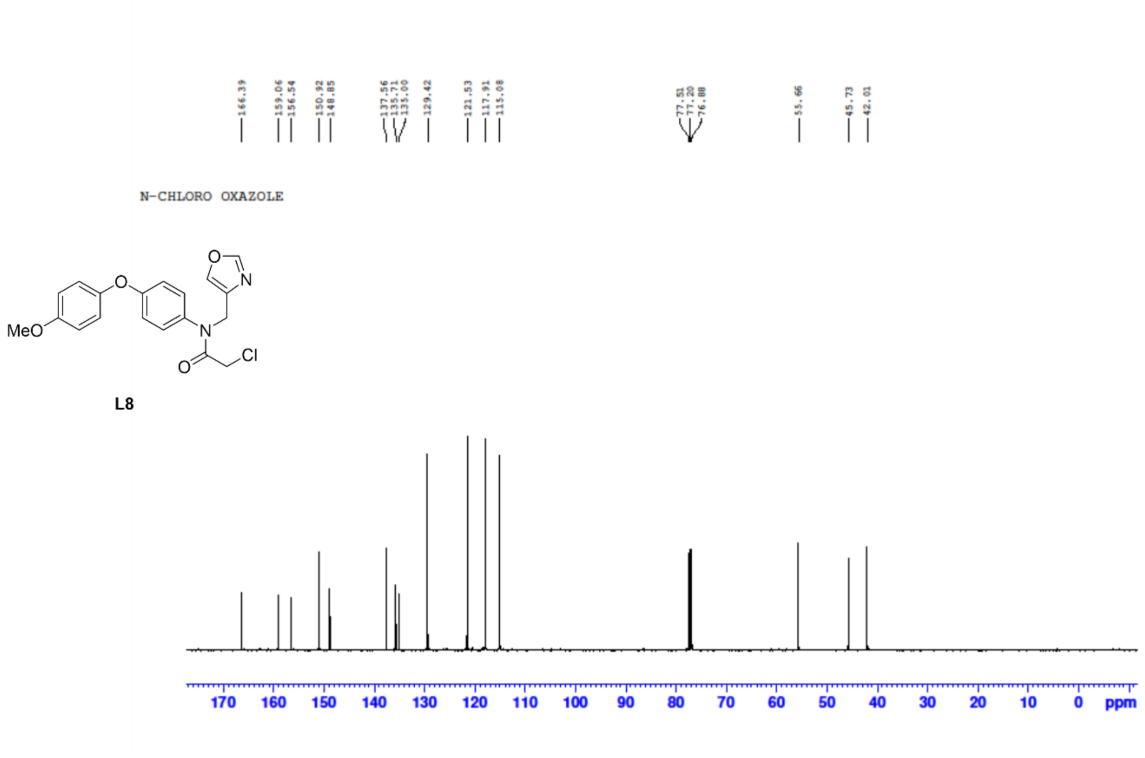


#### [1]H NMR of Amine 9:

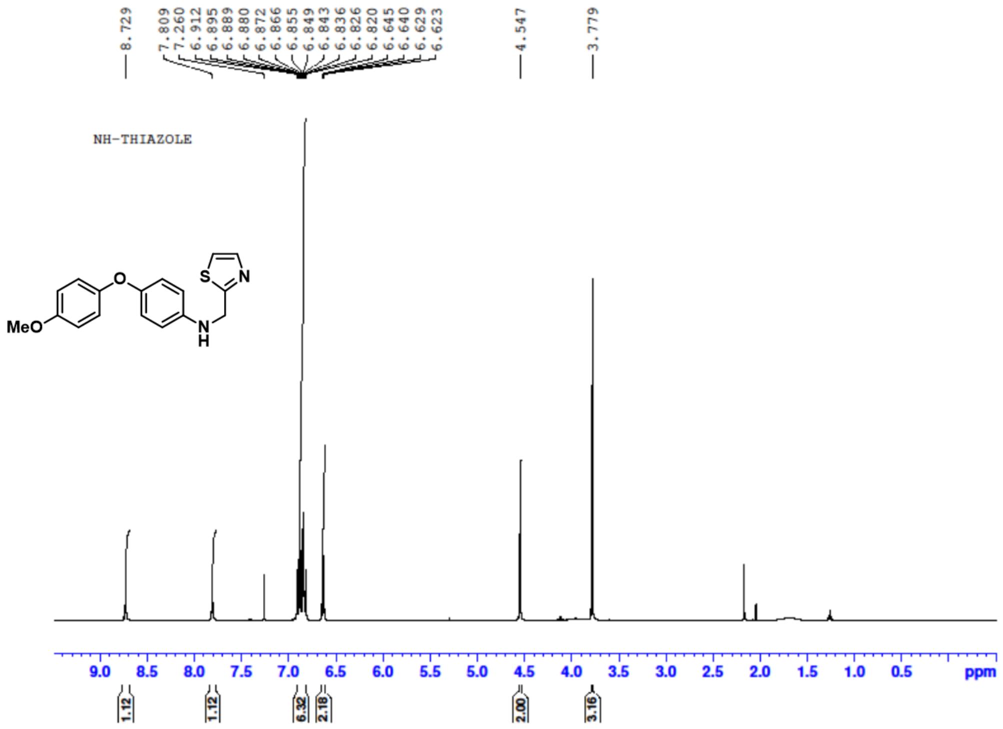

#### [13]C NMR of Amine 9:

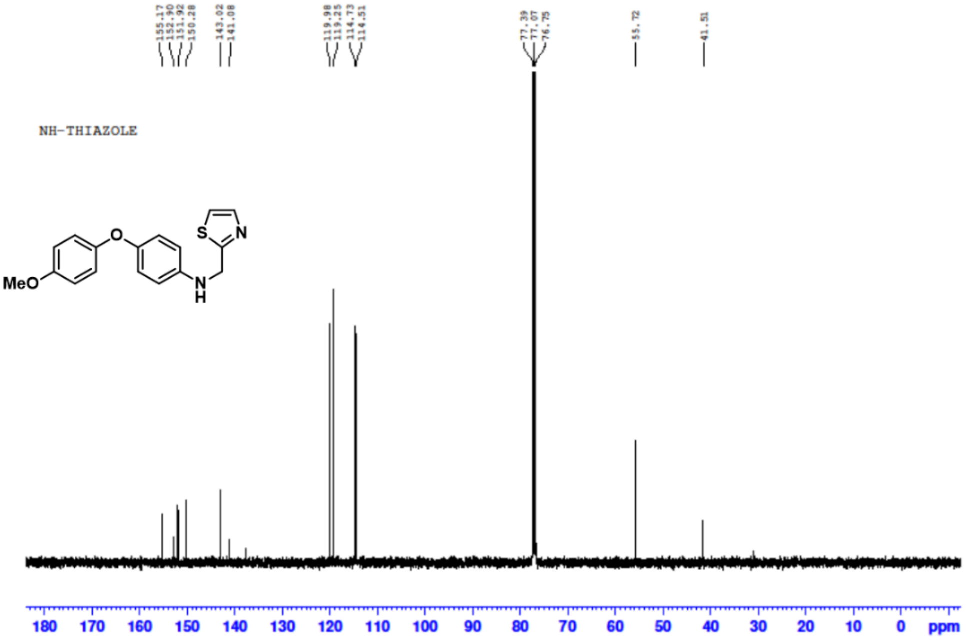


#### [1]H NMR of Ligand 9:

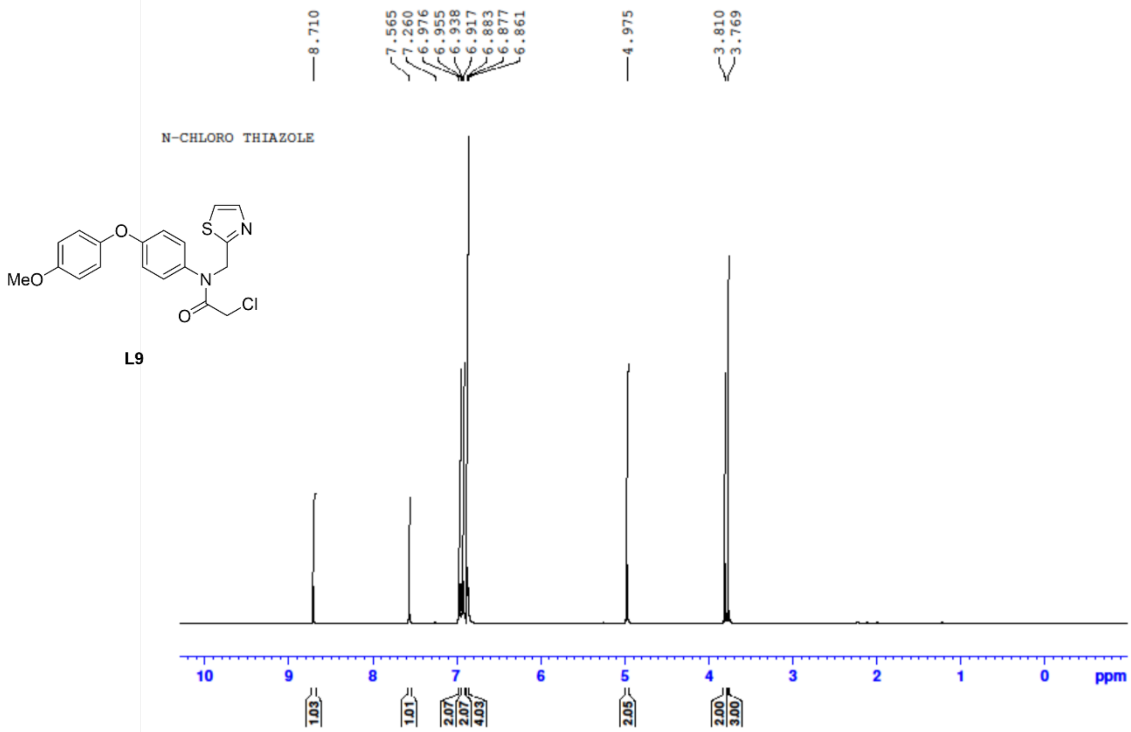

#### $^{13}C$ NMR of Ligand 9:

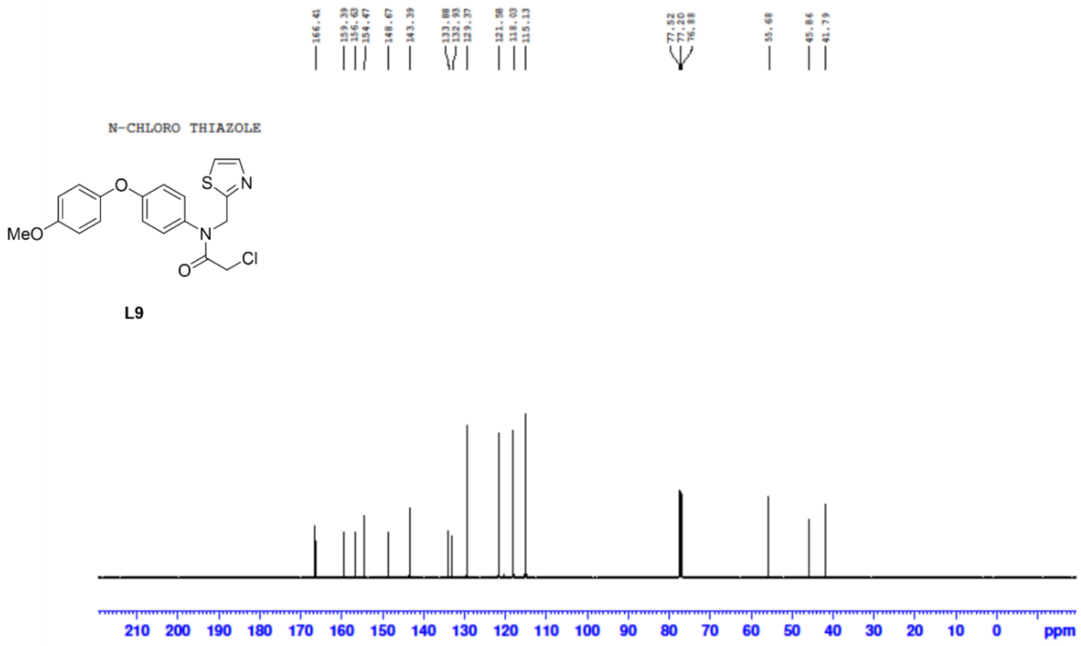


#### $^{1}H$ NMR of Amine 10:

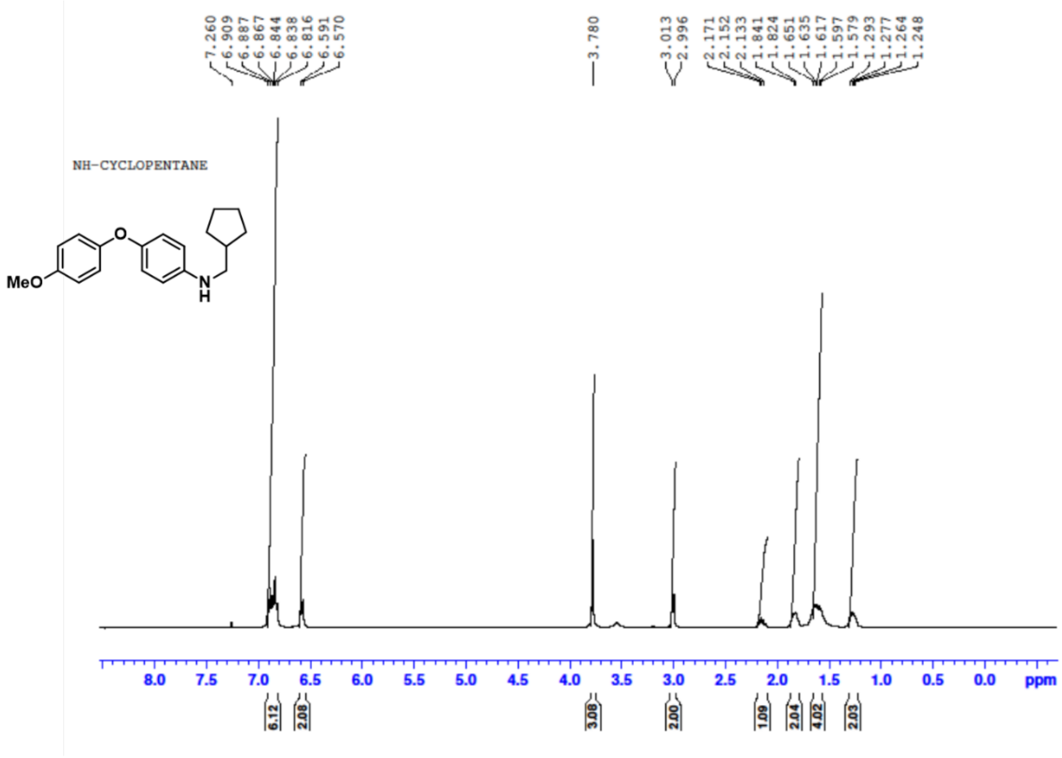

$^{13}$C NMR of Amine 10:

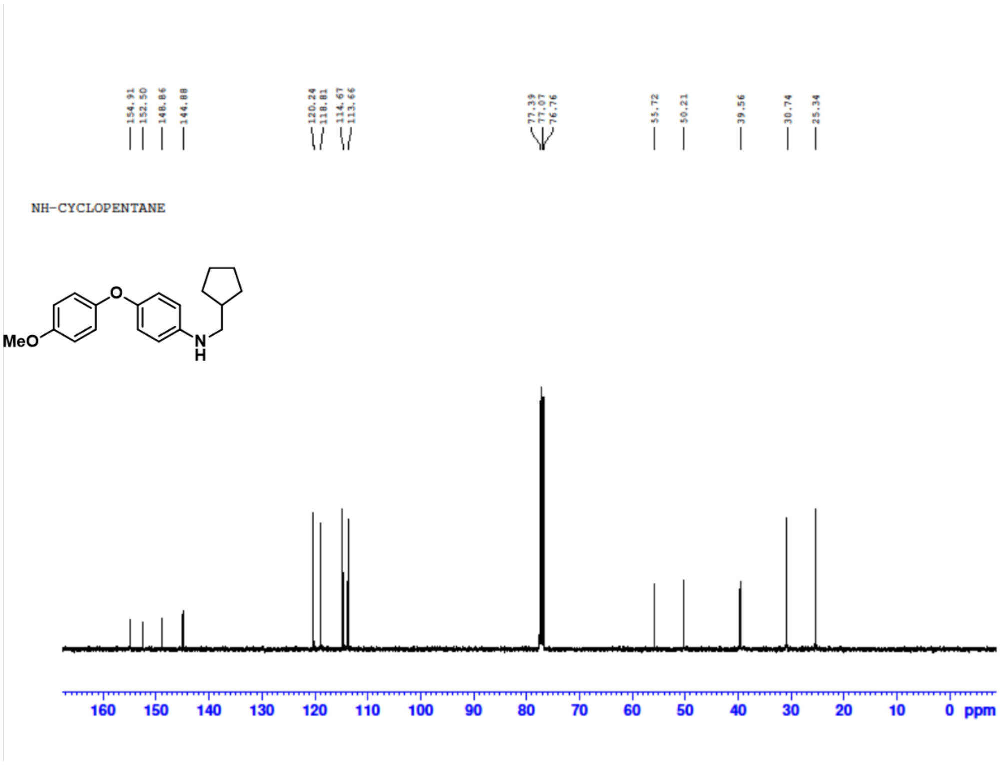


$^{1}$H NMR of Ligand 10:

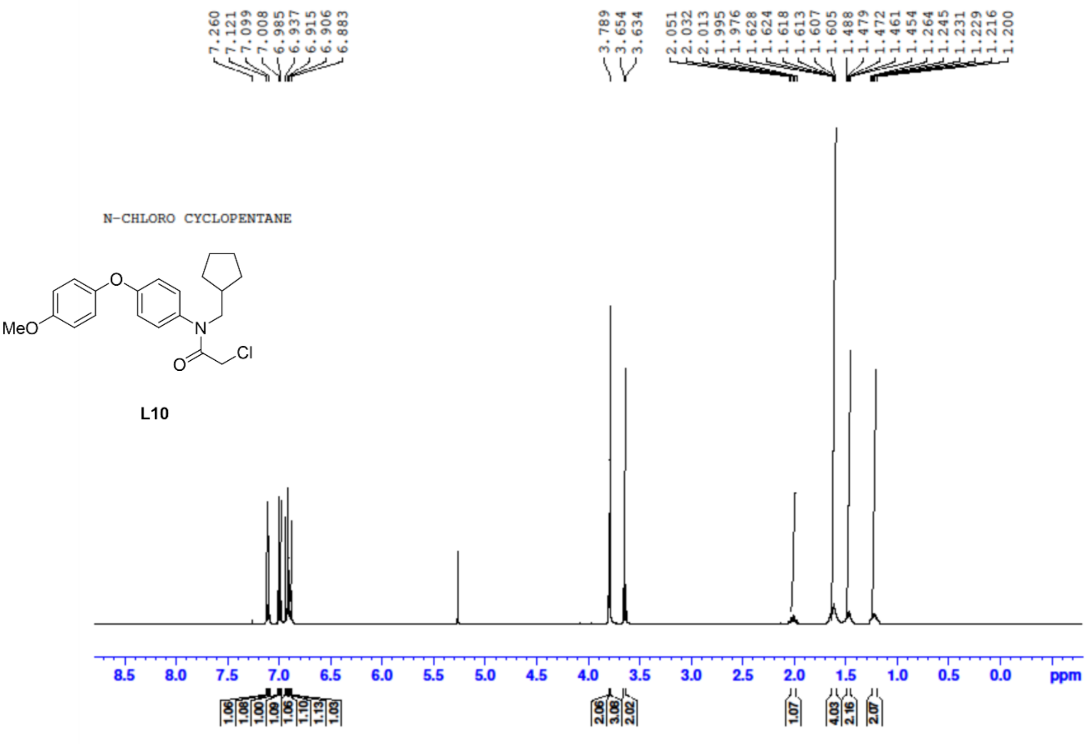

$^{13}$C NMR of Ligand 10:

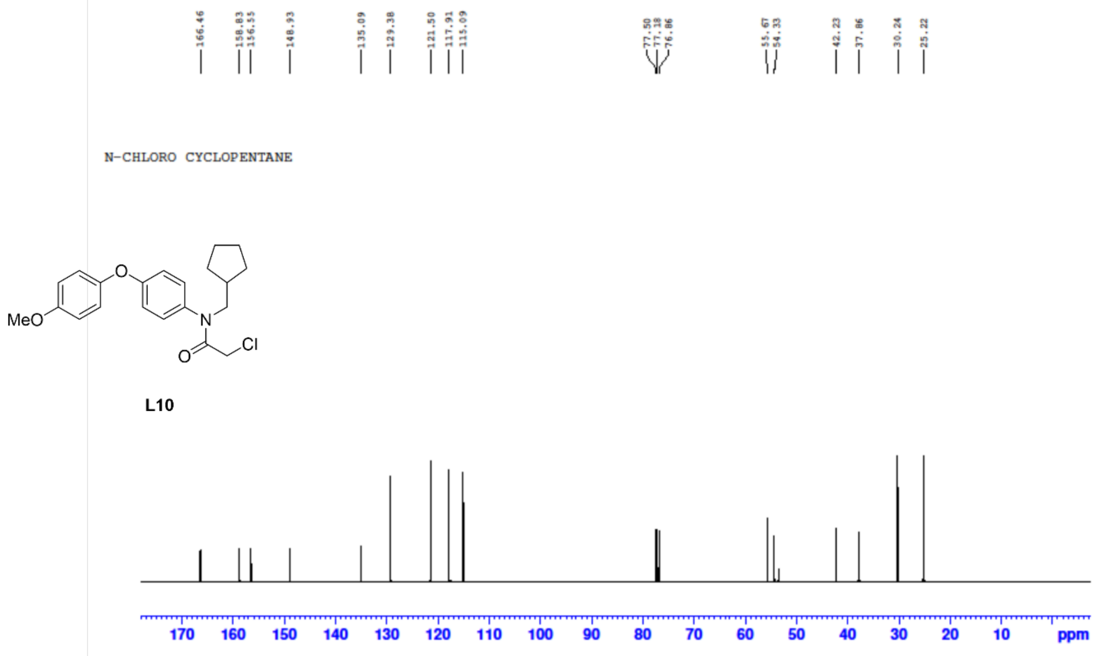


$^{1}$H NMR of Amine 11:

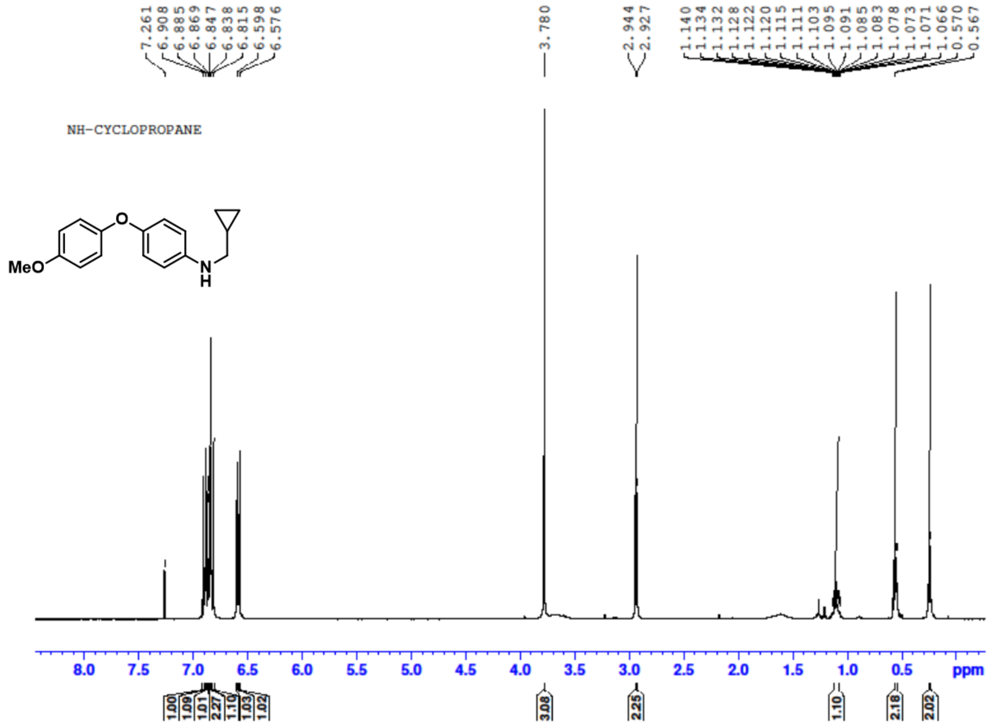

#### $^{13}$C NMR of Amine 11:

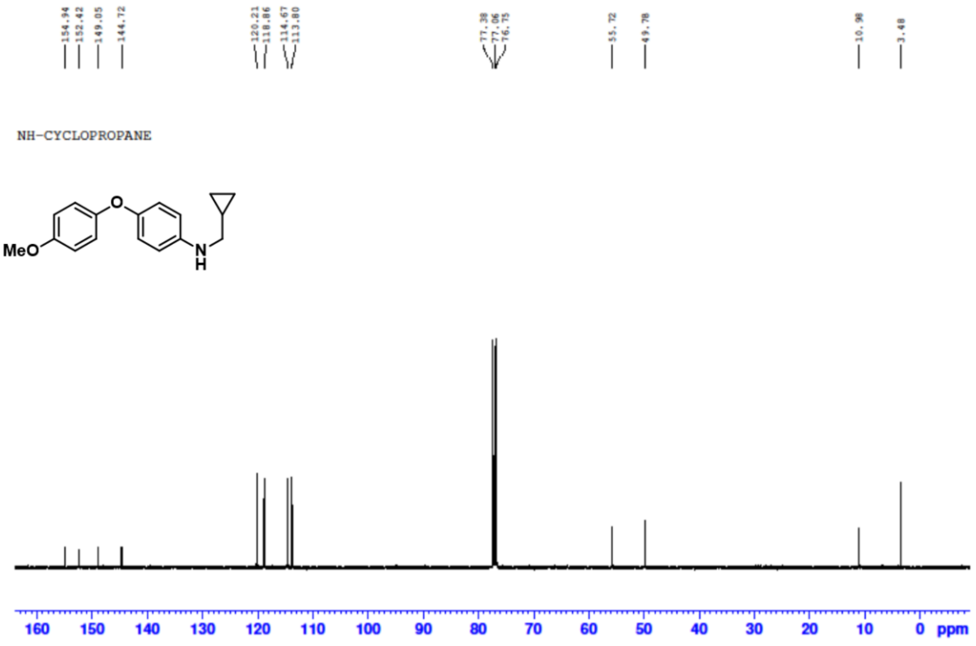


#### $^{1}$H NMR of Ligand 11:

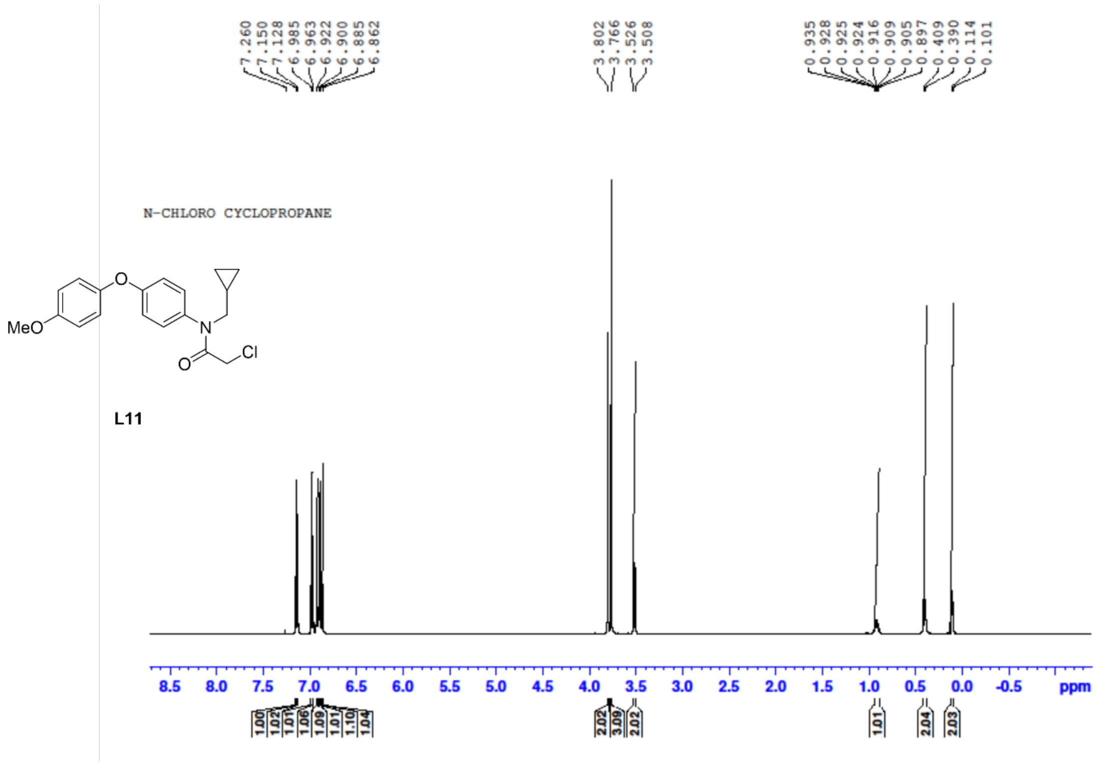

#### $^{13}C$ NMR of Ligand 11:

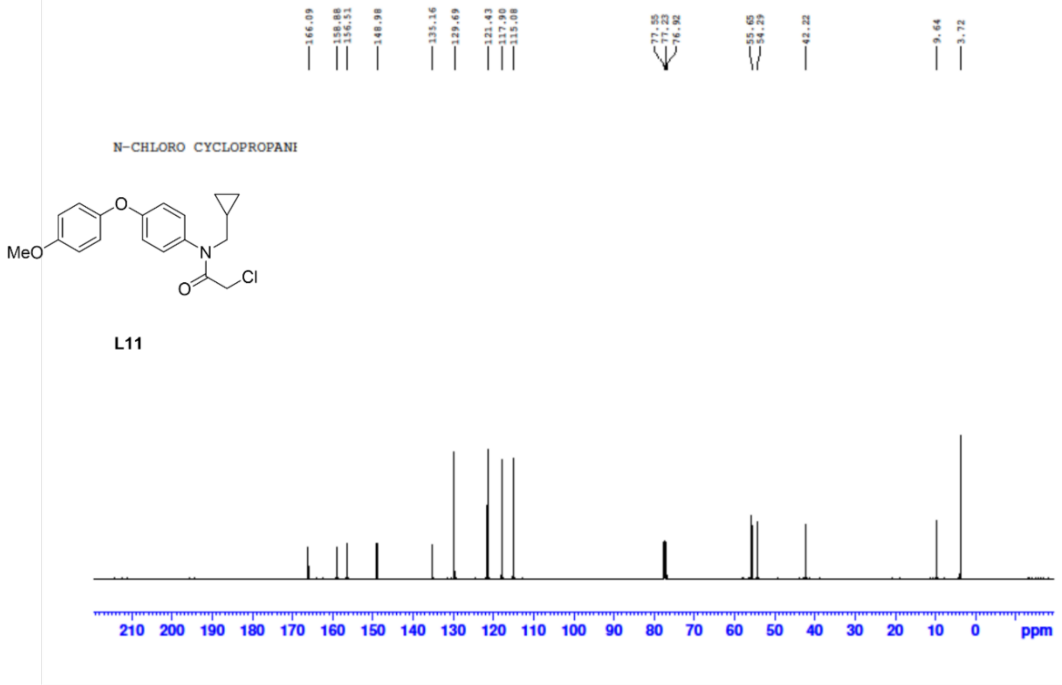


#### $^{1}H$ NMR of Amine 12:

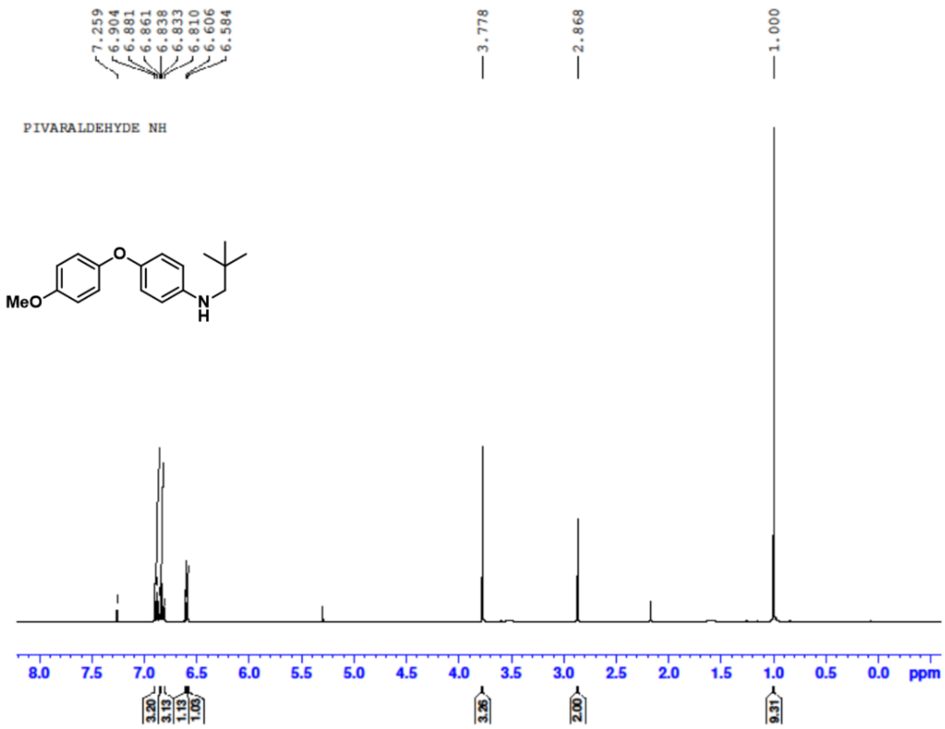

**[13]C NMR of Amine 12:**

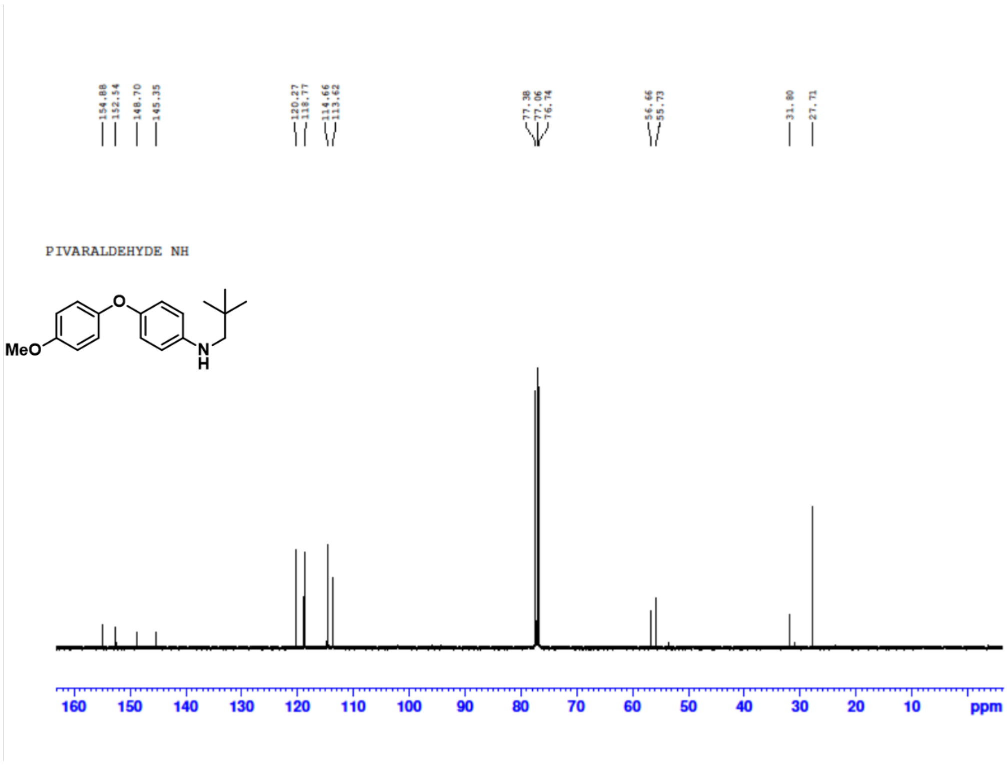


**[1]H NMR of Ligand 12:**

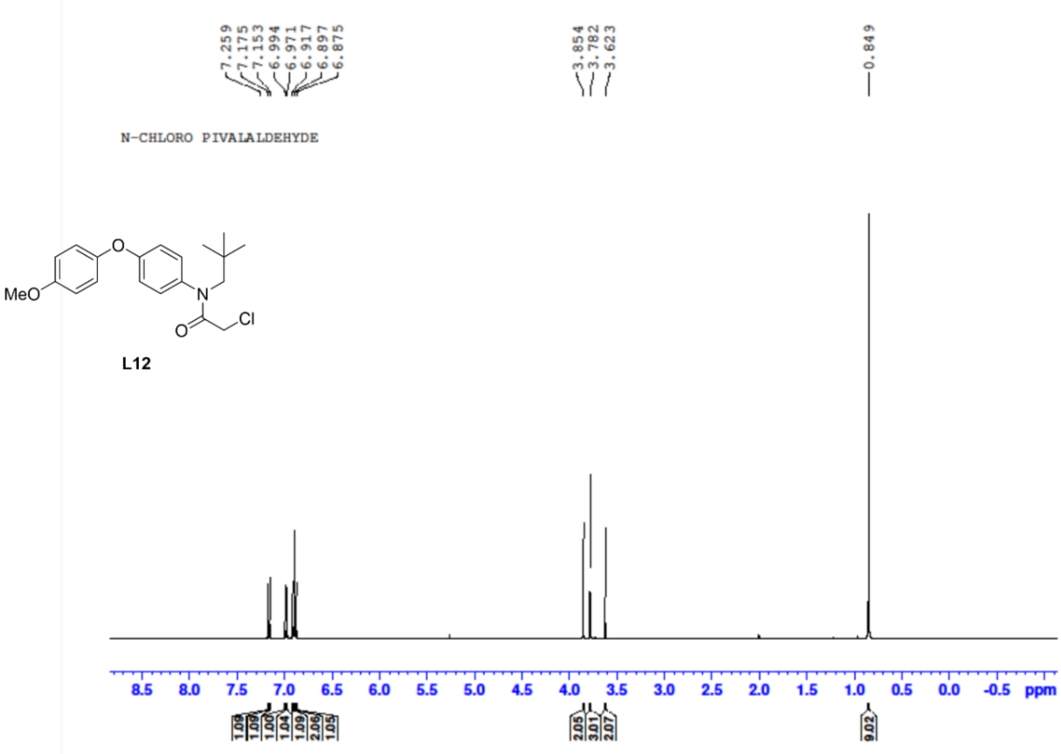

**$^{13}$C NMR of Ligand 12:**

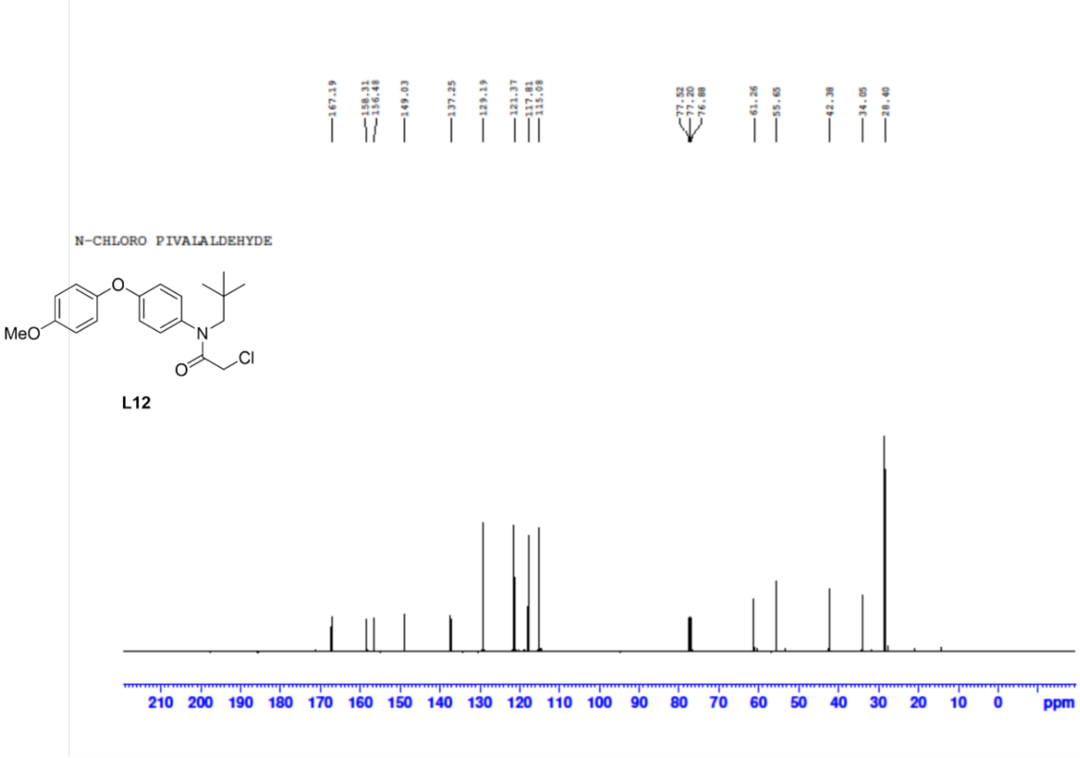


**$^{1}$H NMR of Amine 13:**

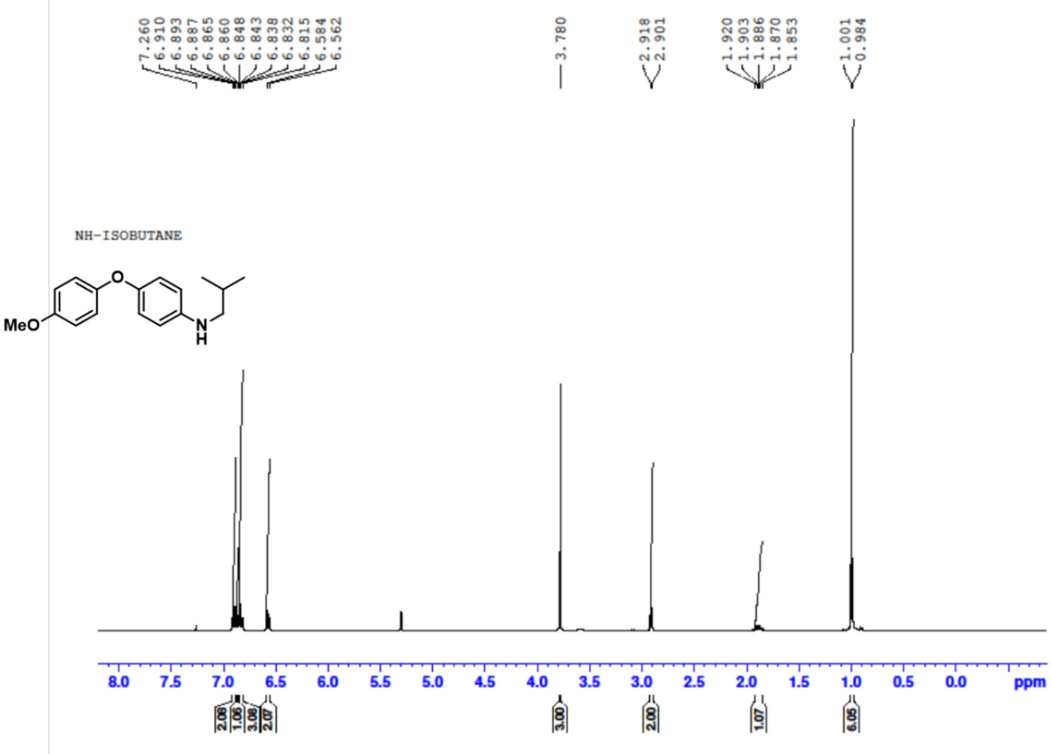

[13]C NMR of Amine 13:

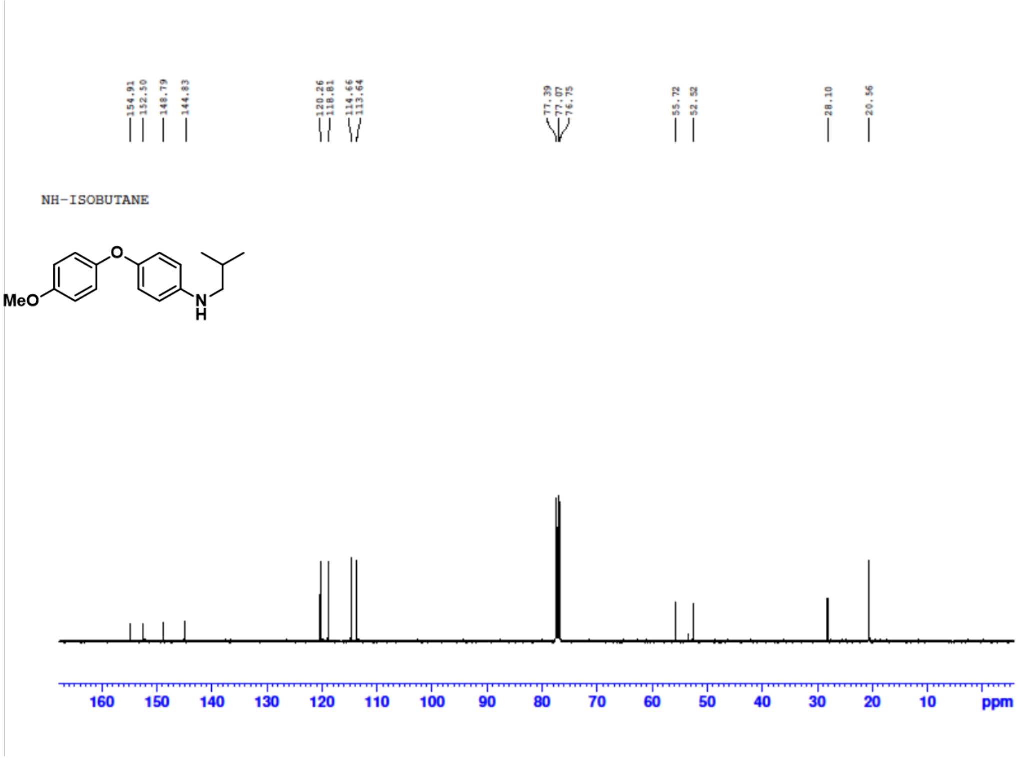


[1]H NMR of Ligand 13:

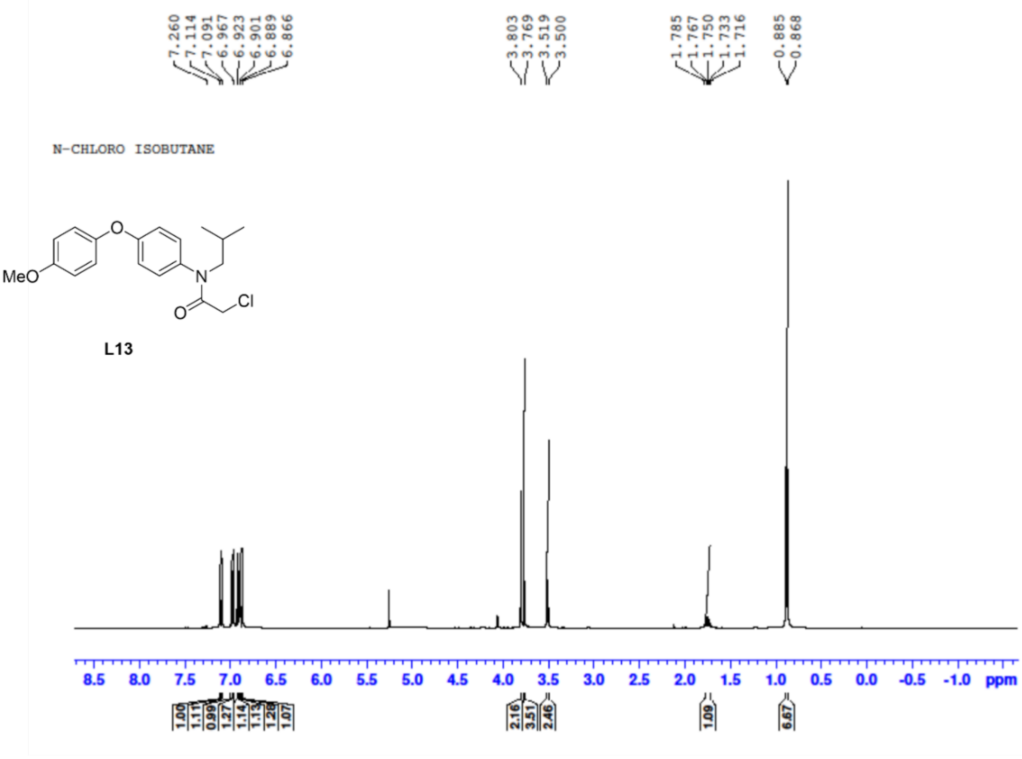

**$^{13}$C NMR of Ligand 13:**

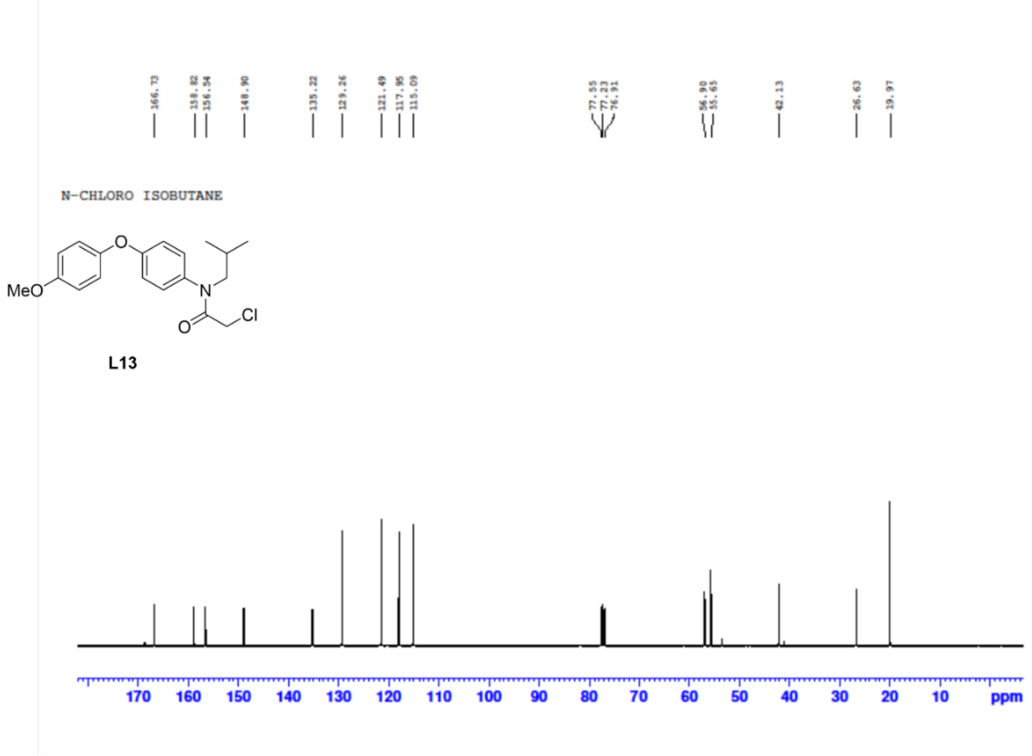


**$^{1}$H NMR of Amine 14:**

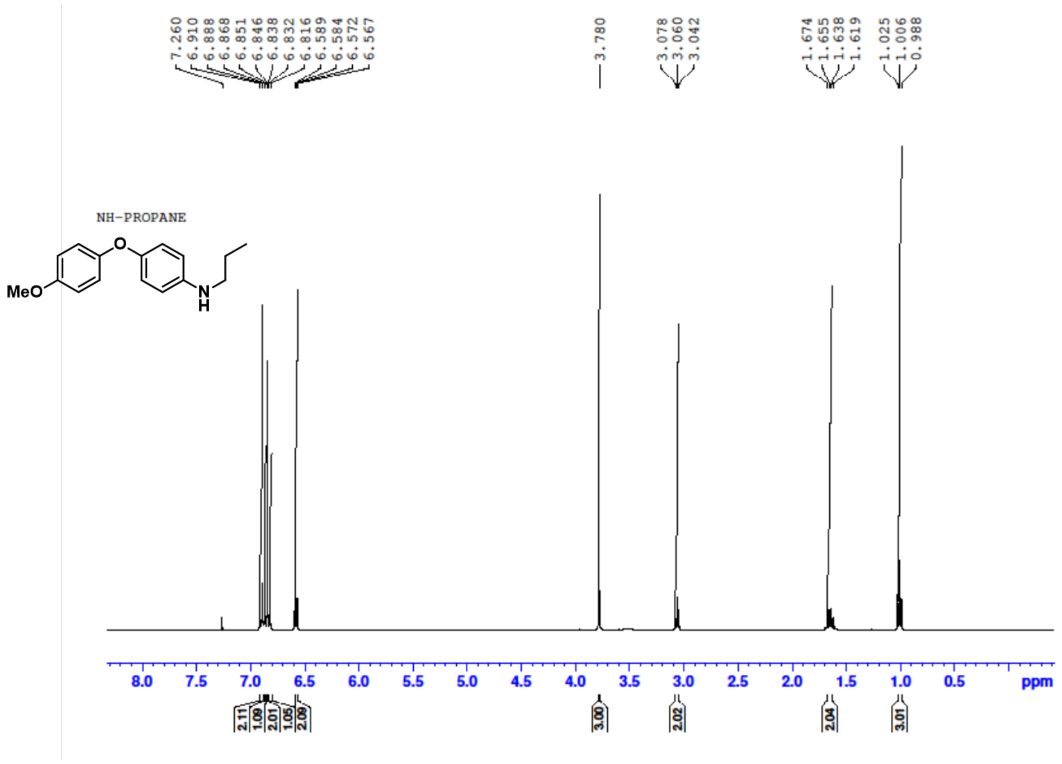

#### $^{13}$C NMR of Amine 14:

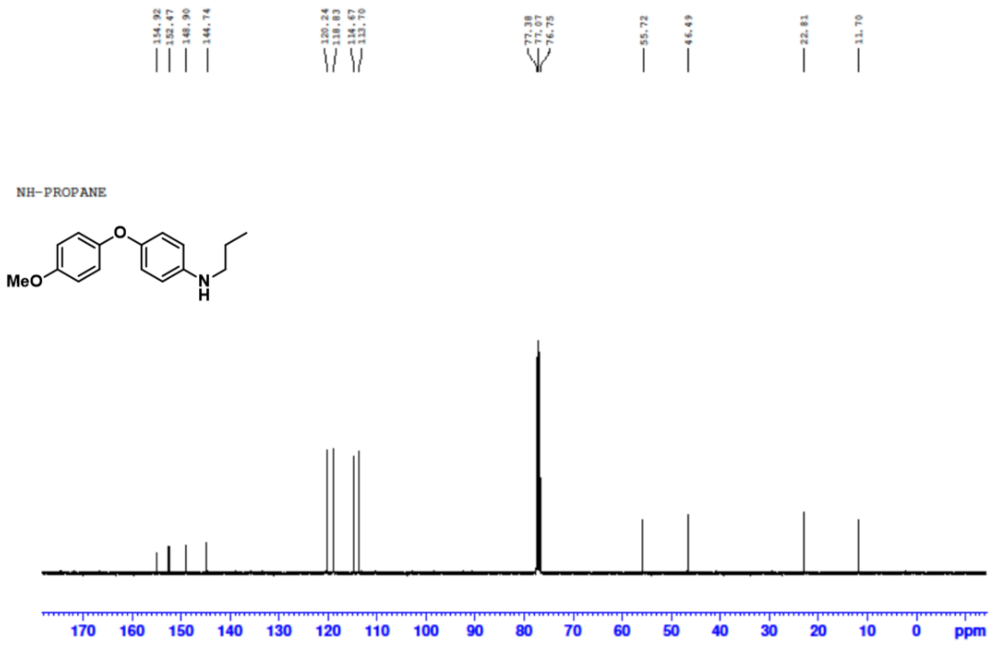


#### $^{1}$H NMR of Ligand 14:

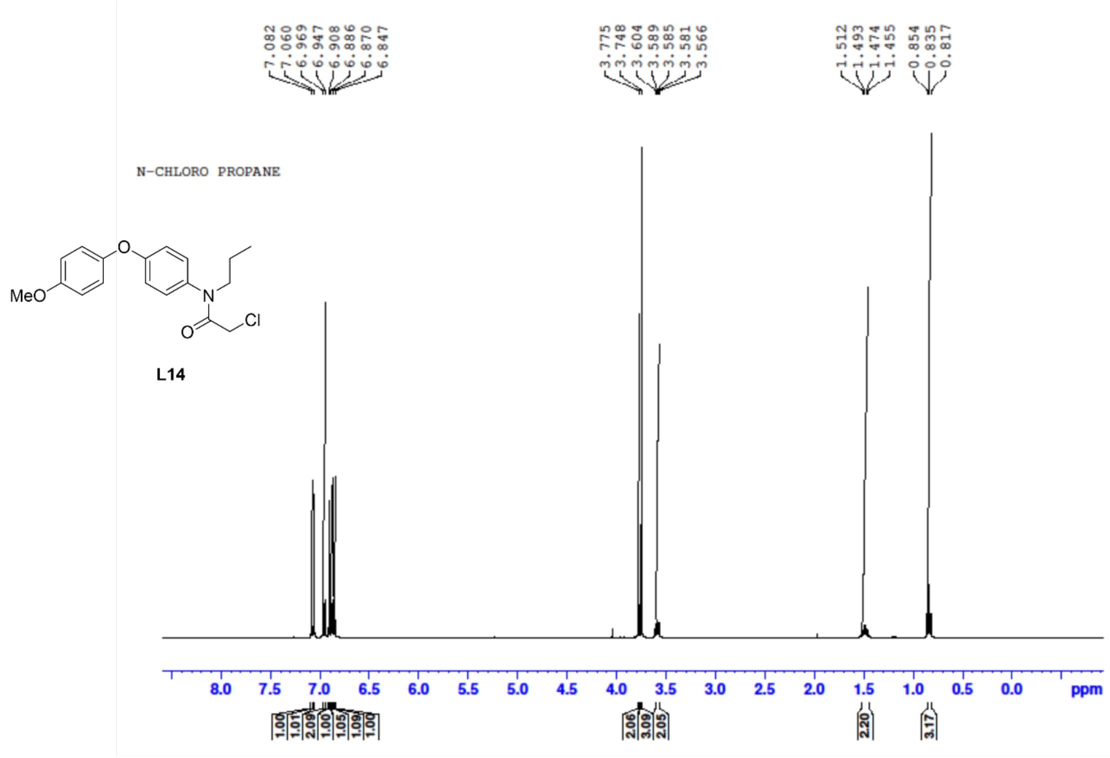

**$^{13}$C NMR of Ligand 14:**

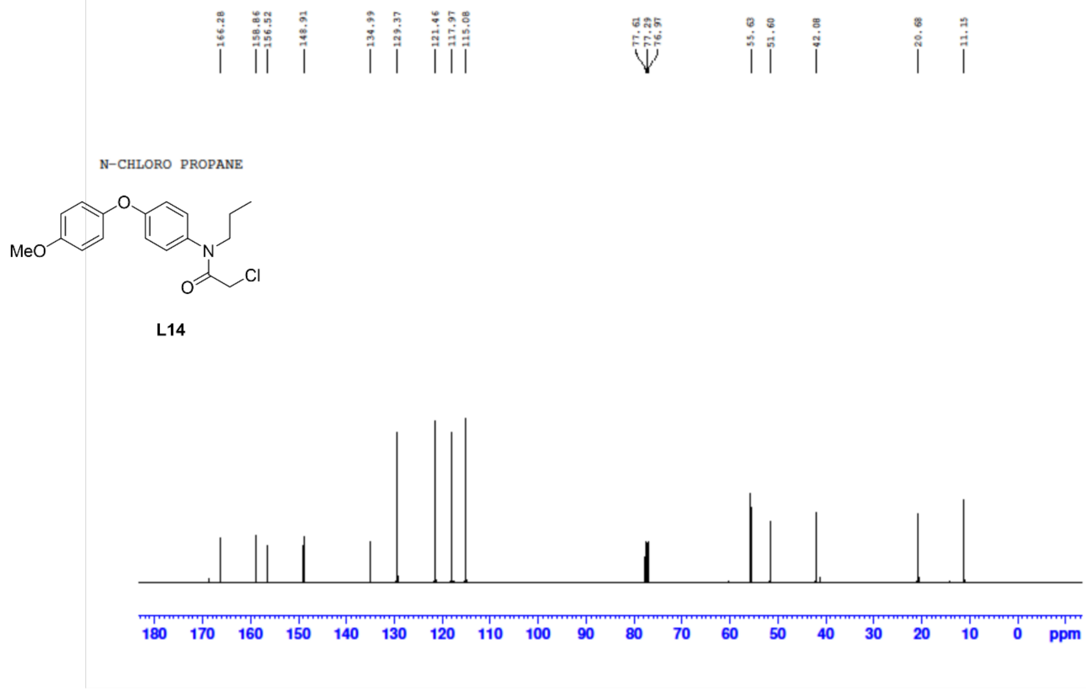


**$^{1}$H NMR of Inactive Ligand:**

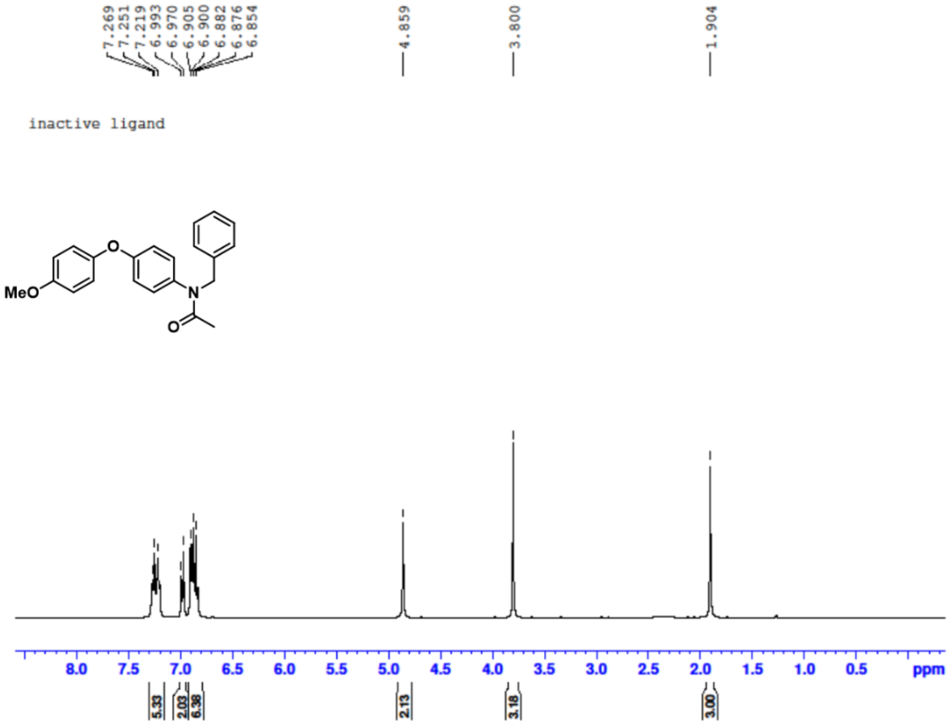


**$^{13}$C NMR of Inactive Ligand:**

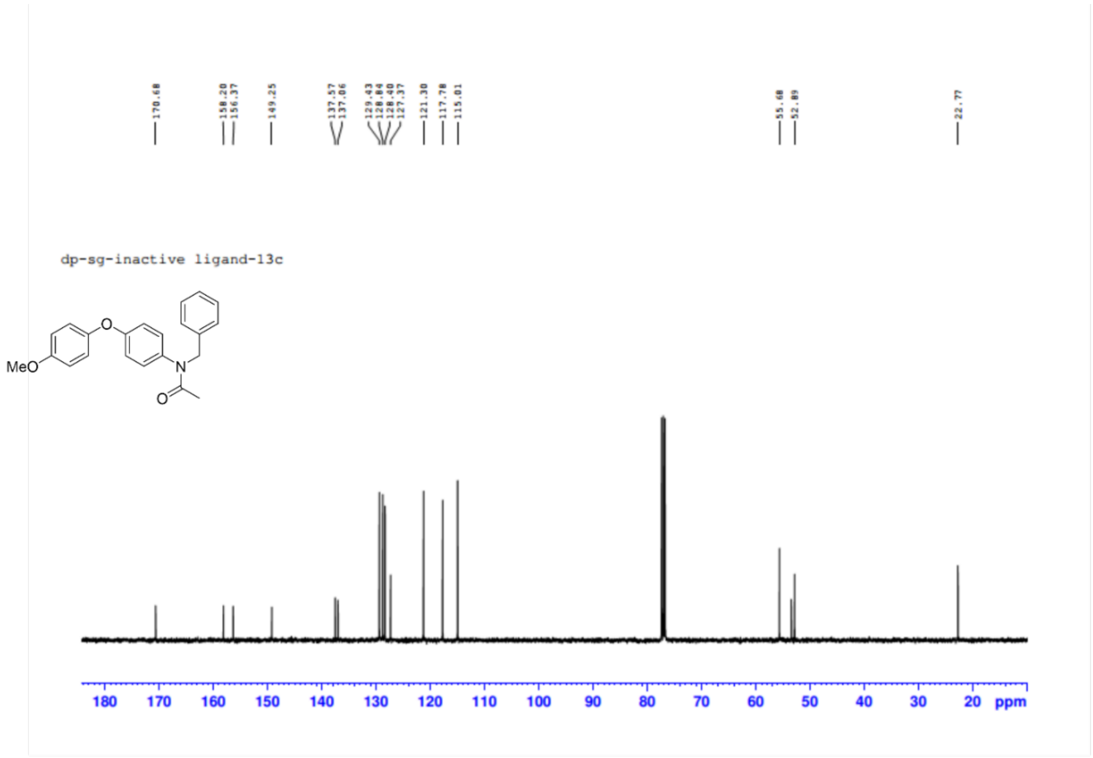


**$^{1}$H NMR of VHL ligand:**

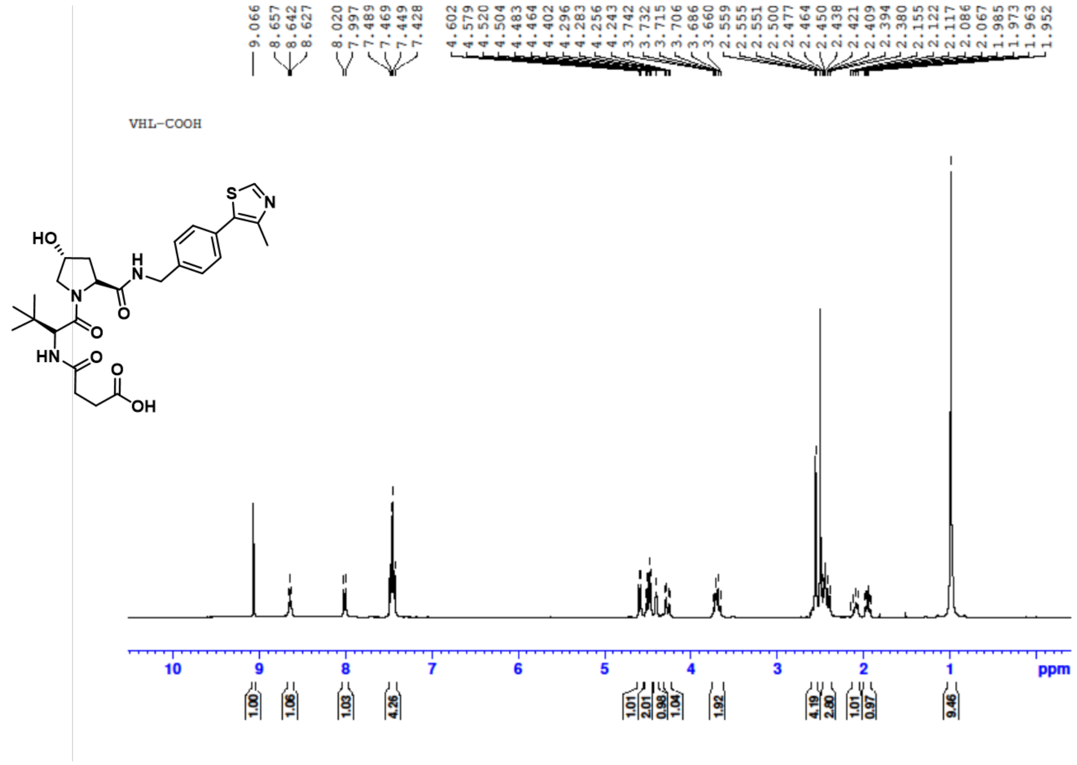

#### $^{13}$C NMR of VHL ligand:

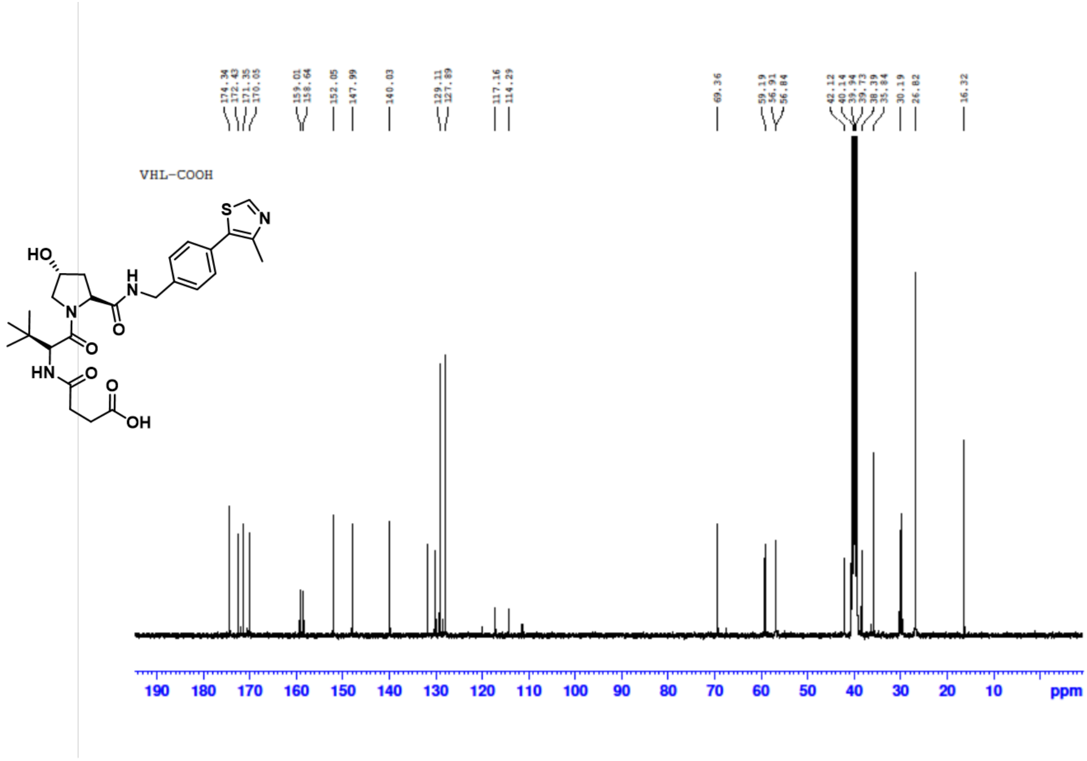


#### HRMS of VHL ligand:

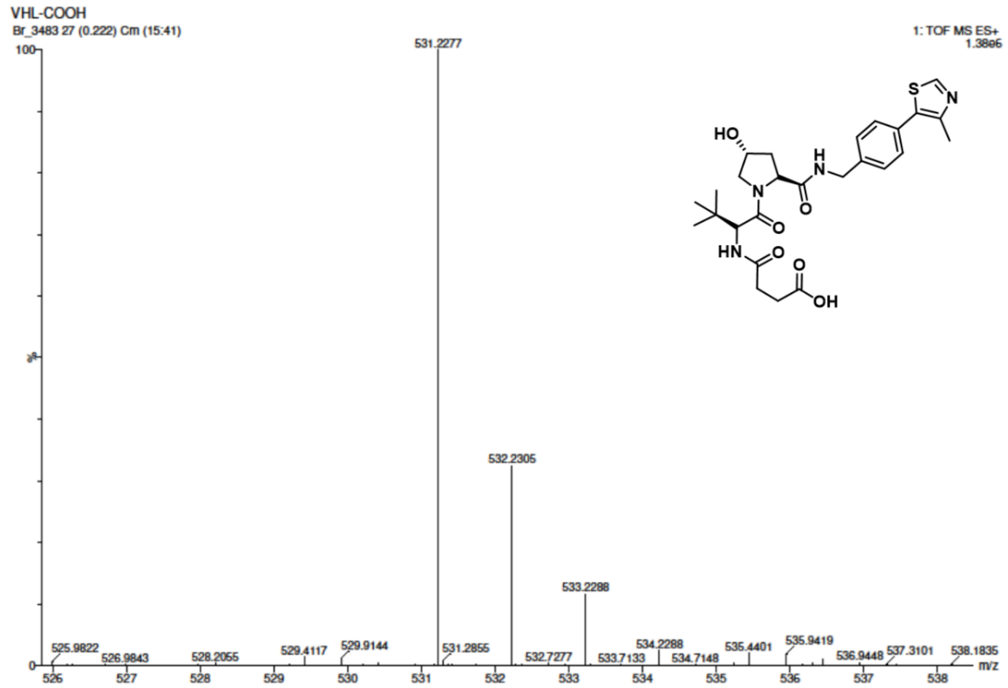

#### 1H NMR of R4VP:

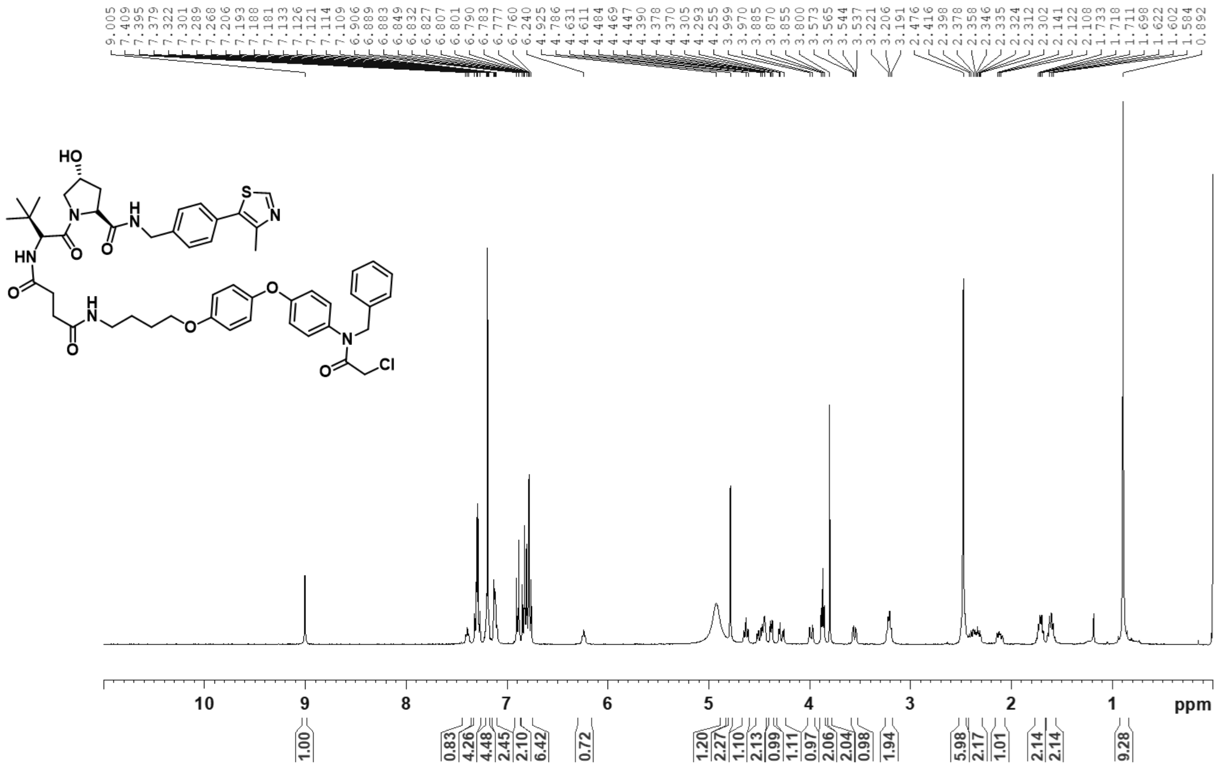


#### 13C NMR of R4VP:

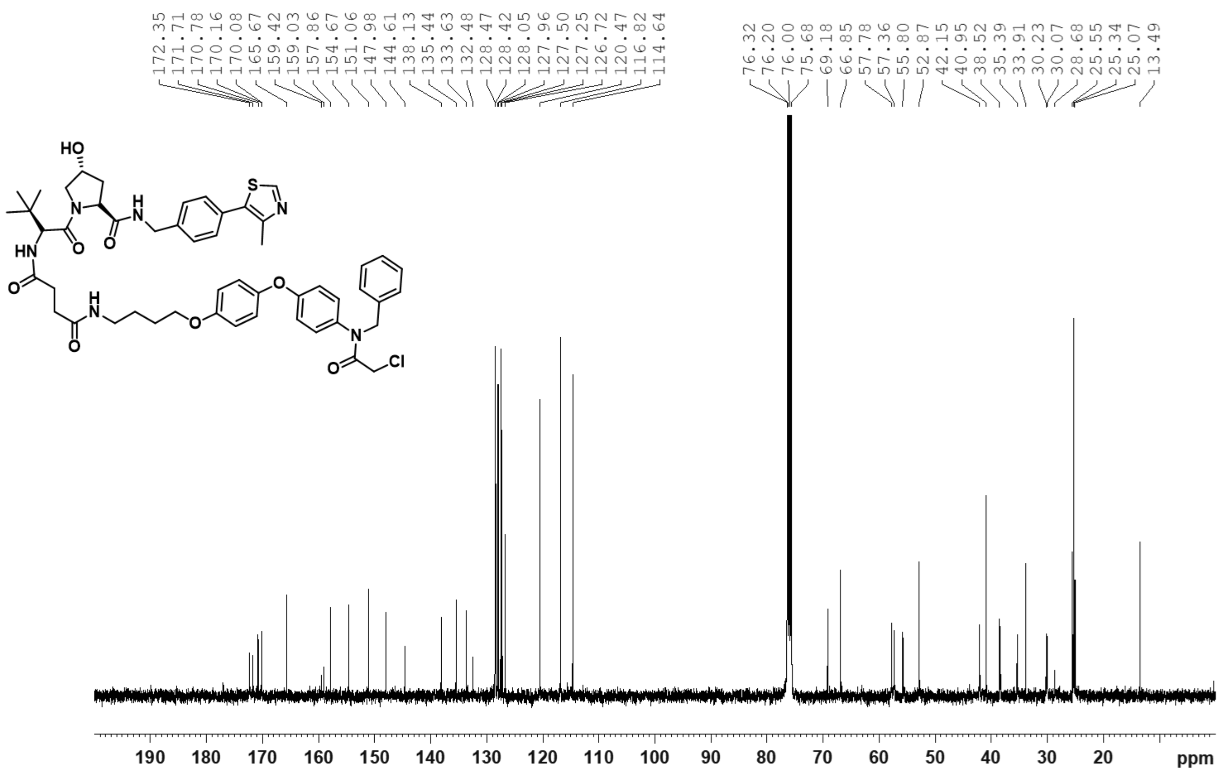

**HRMS of R4VP:**

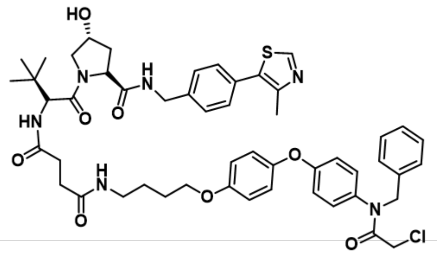

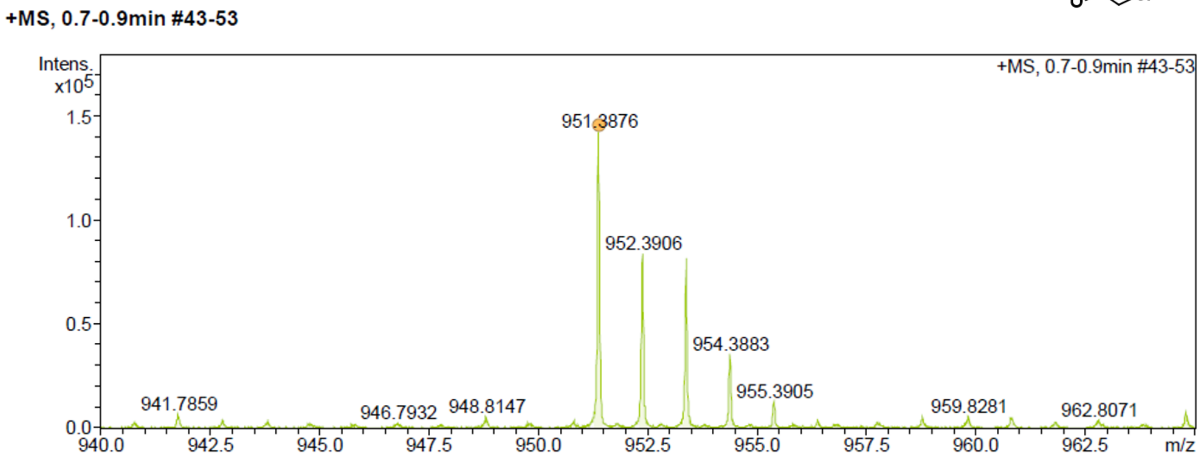


**[1]H NMR of Compound 3:**

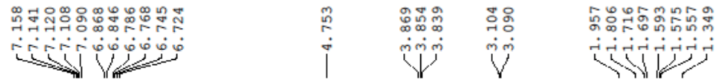

Control NH Boc

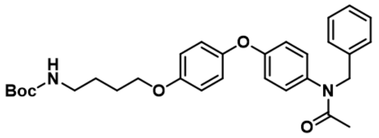

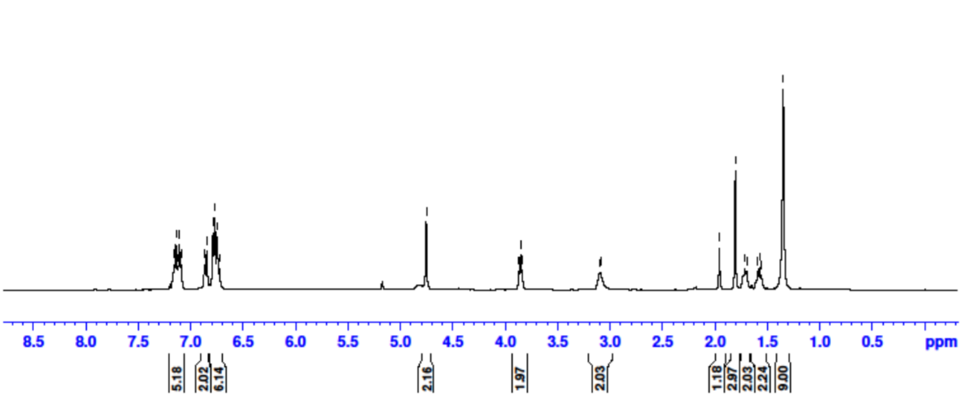

#### [13]NMR of Compound 3:

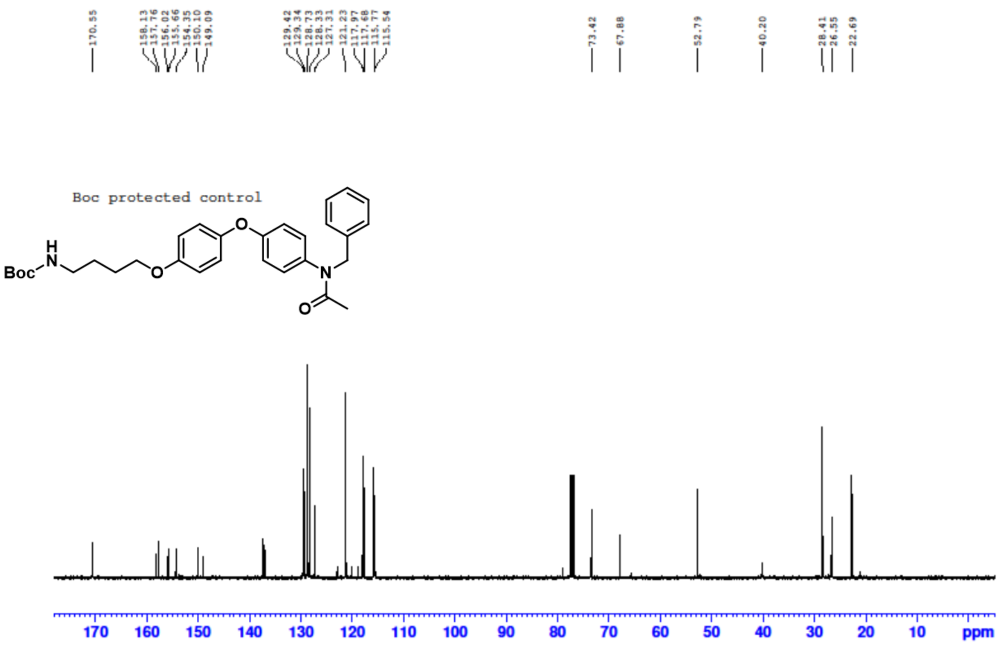


#### HRMS of Compound 3:

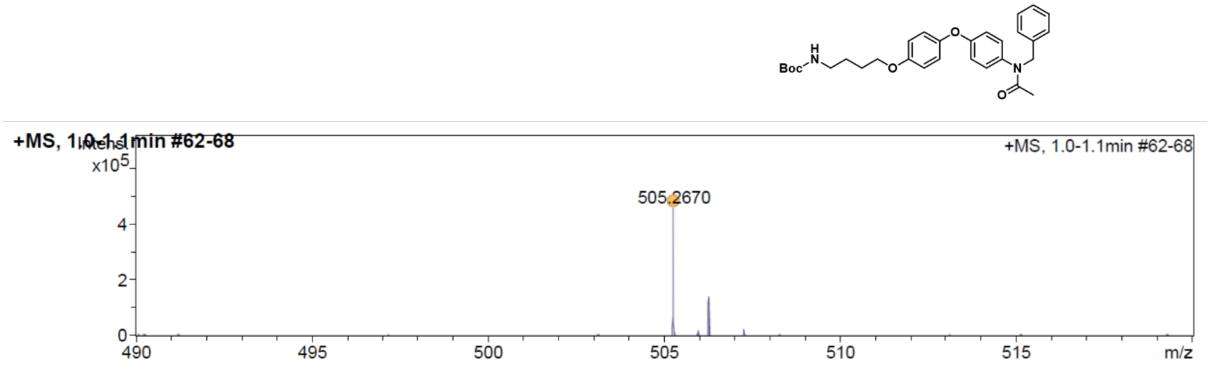

##### $^{1}$H NMR of inactive R4VP:

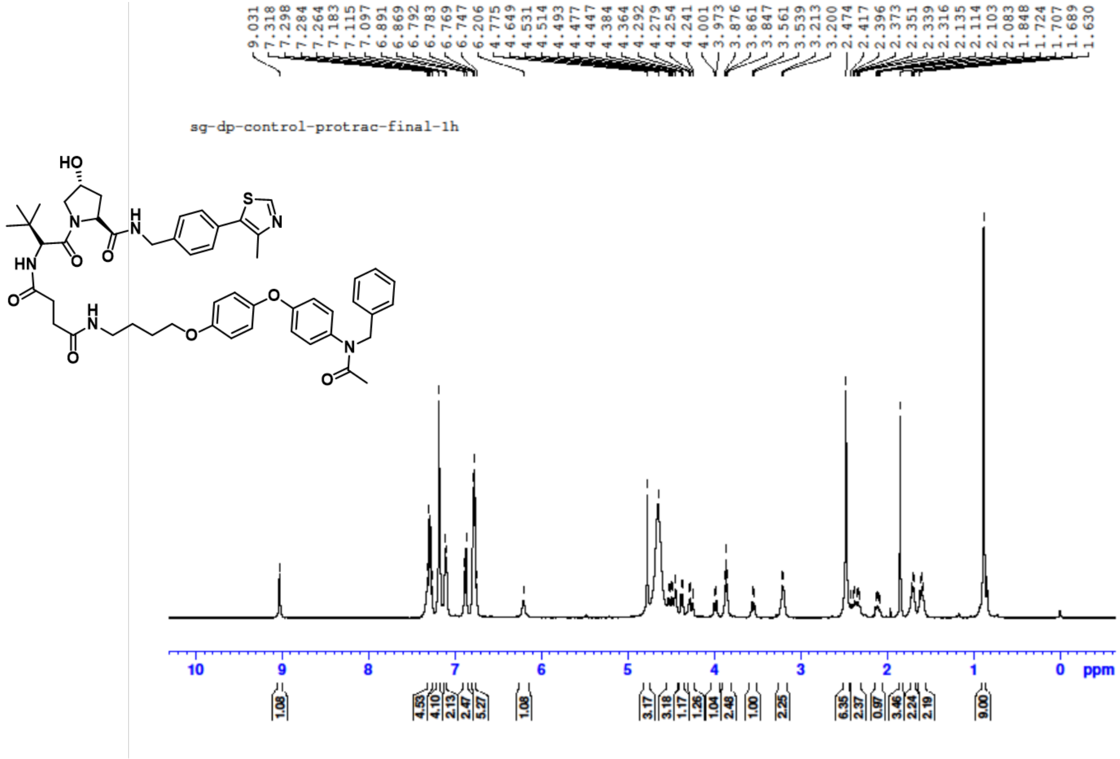


##### $^{13}$C NMR of inactive R4VP:

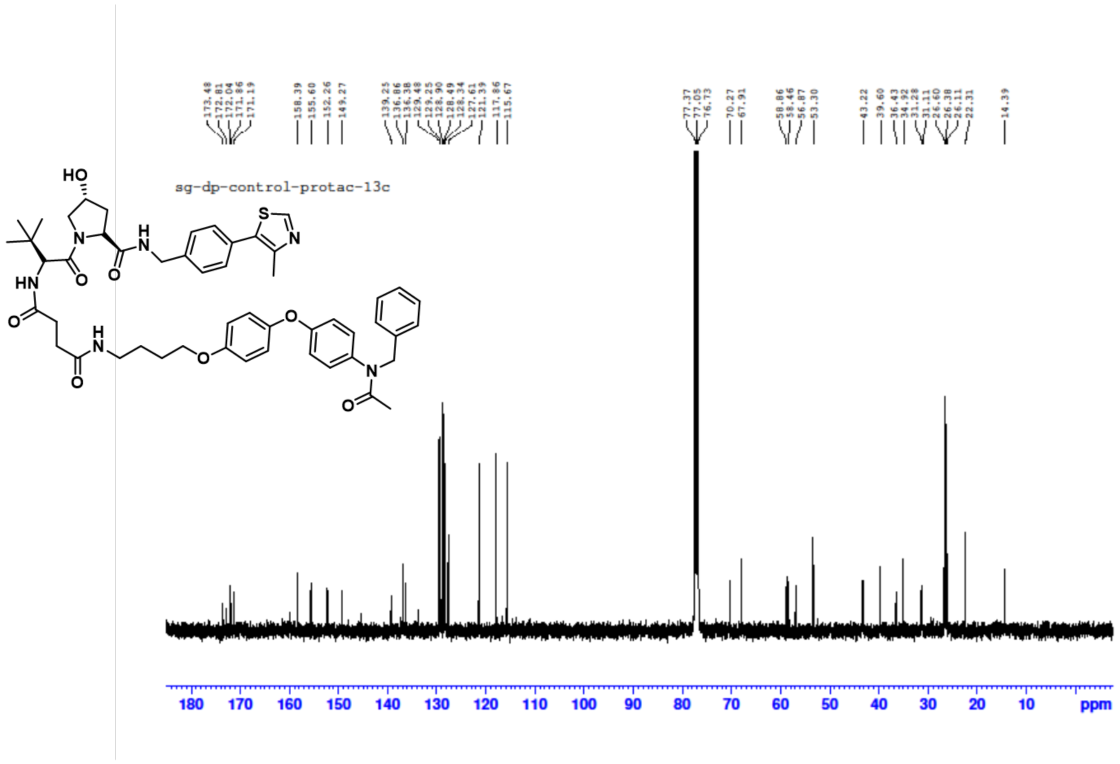

**HRMs inactive R4VP:**

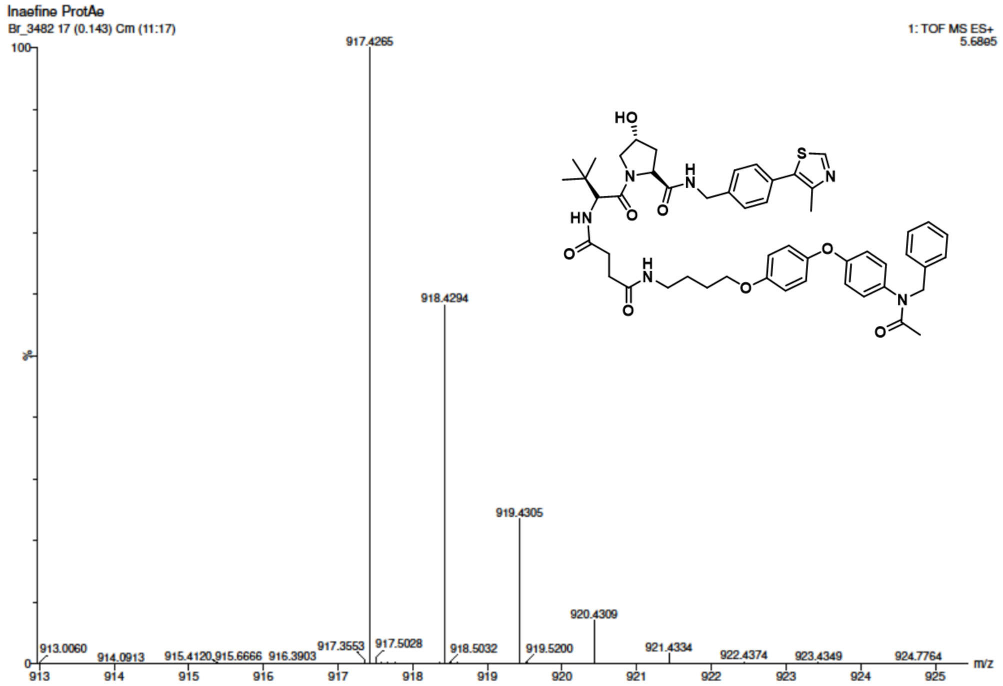


**$^{1}$H NMR of compound 2:**

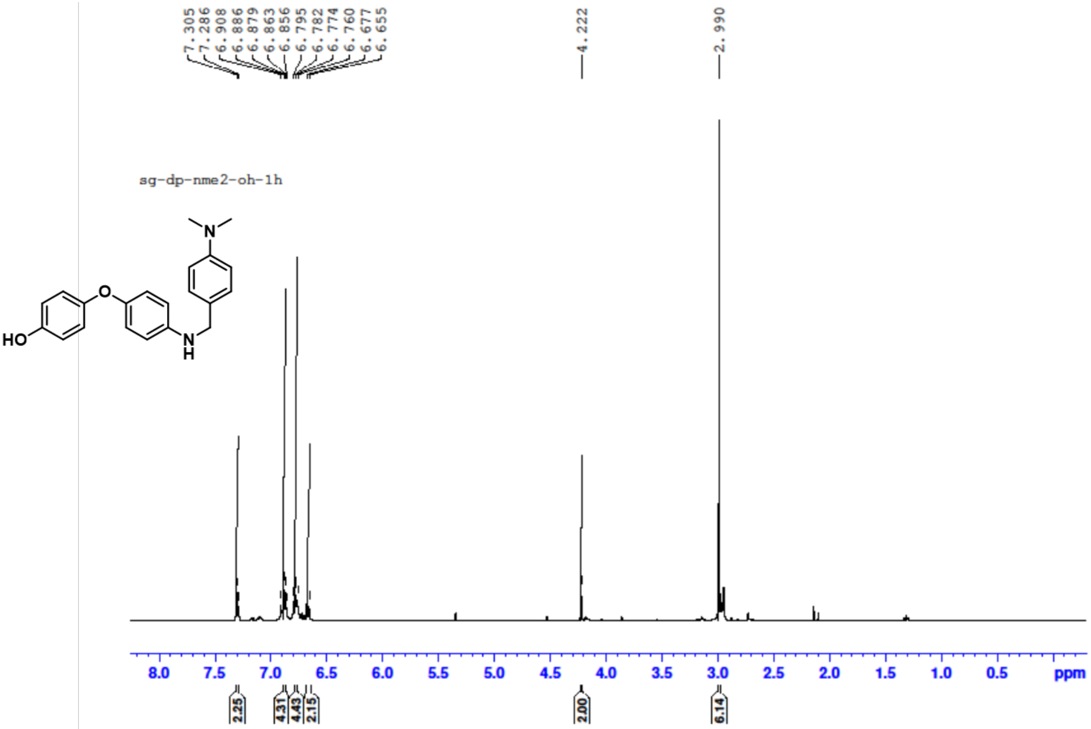

[13]C NMR of compound 2:

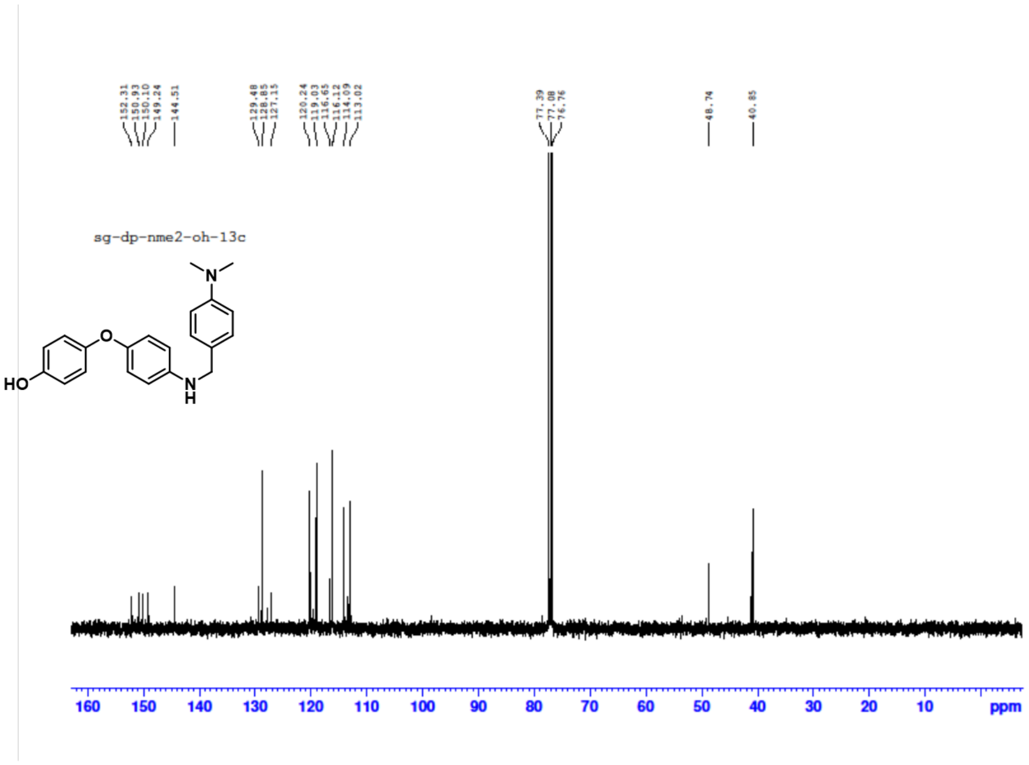


[1]H NMR of N-boc N-Me2:

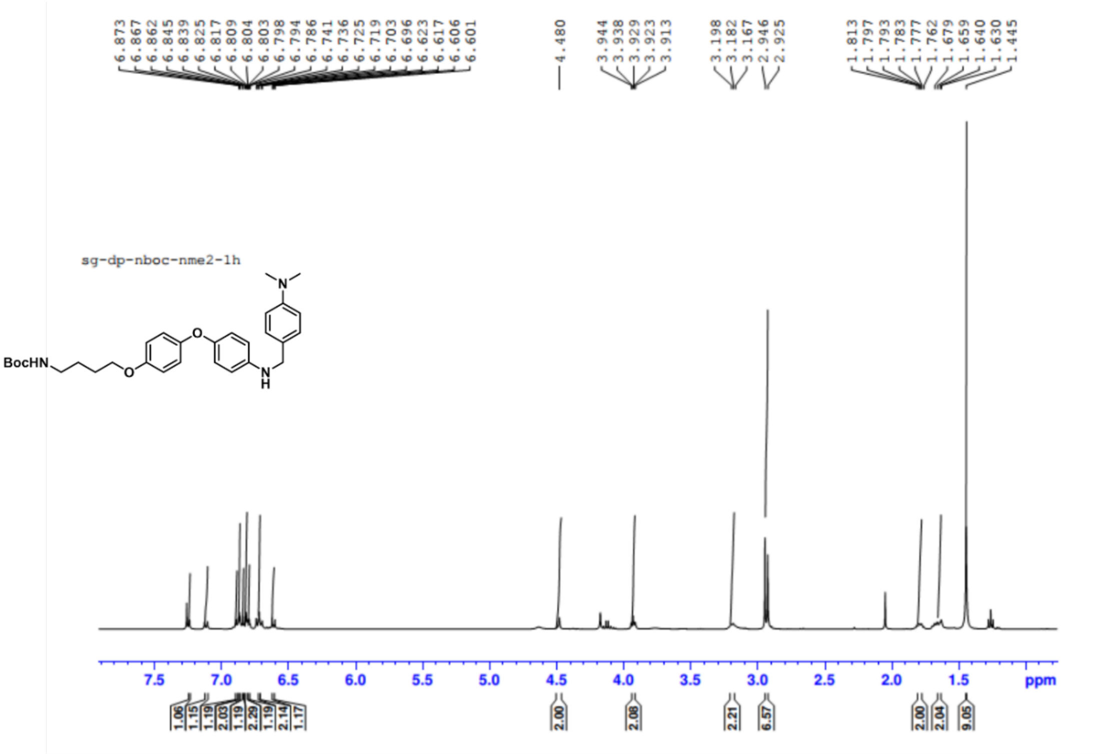

$^{13}$C NMR of N-boc N-Me2:

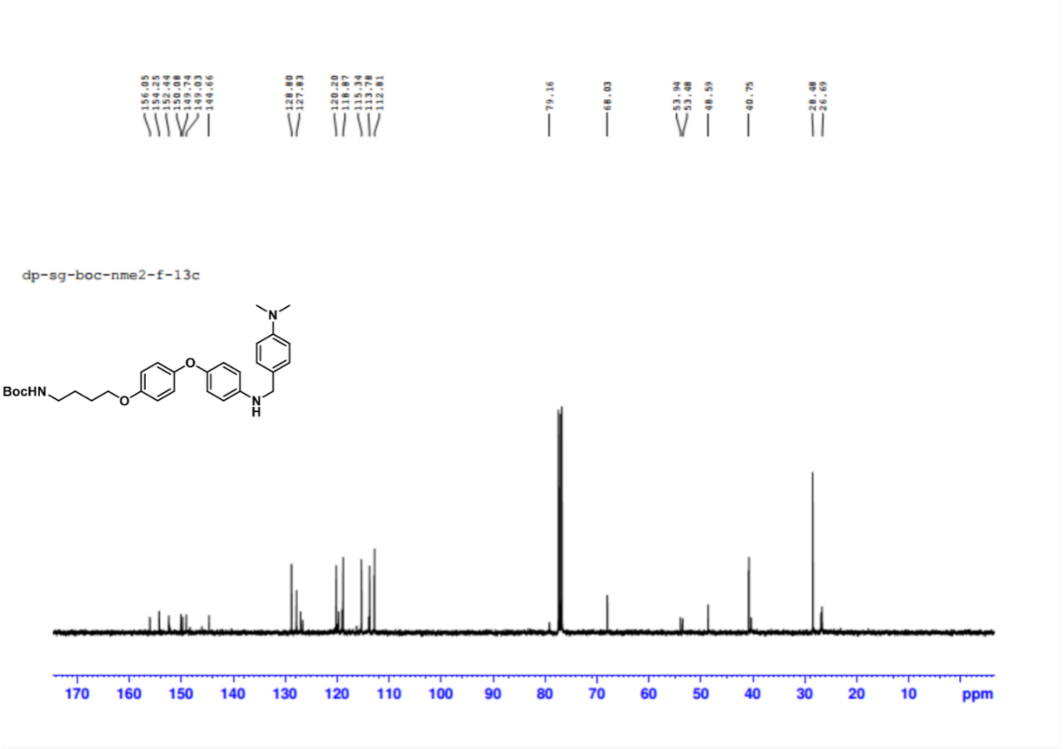


HRMS of N-boc N-Me2:

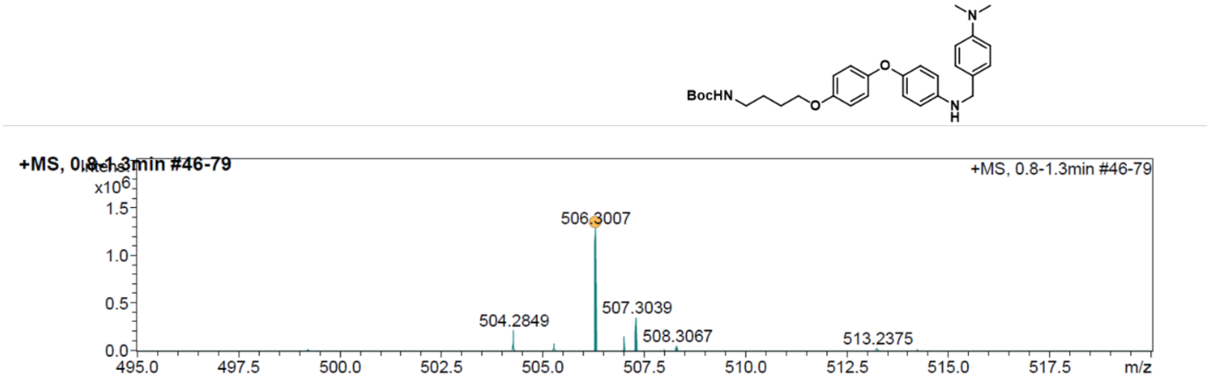

**$^{1}$H NMR of N-Chloro boc N-Me2:**

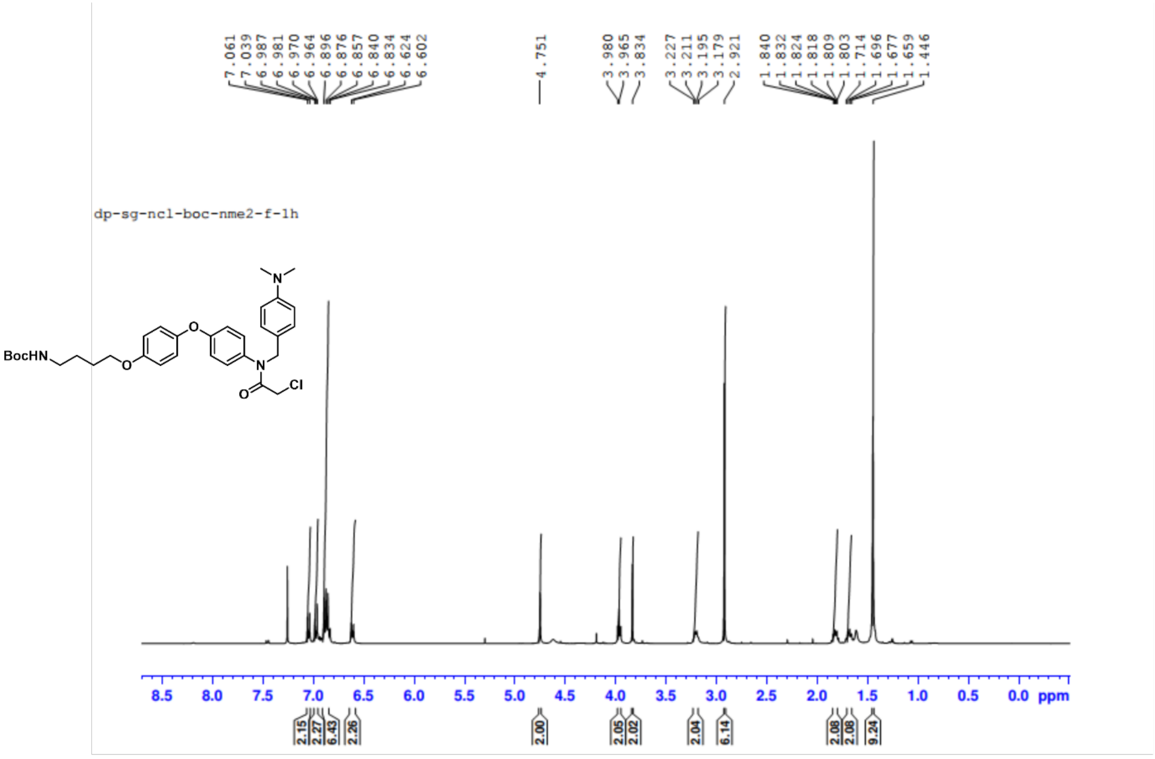


**$^{13}$C NMR of N-Chloro boc N-Me2:**

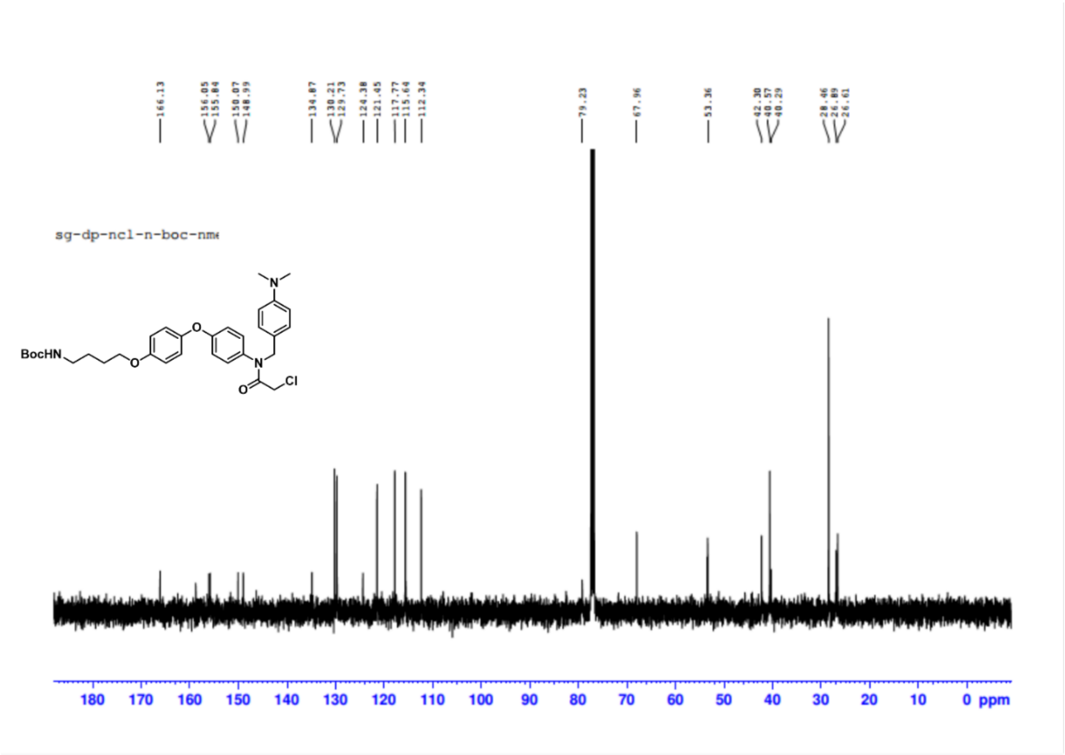

**HRMS of N-Chloro boc N-Me2:**

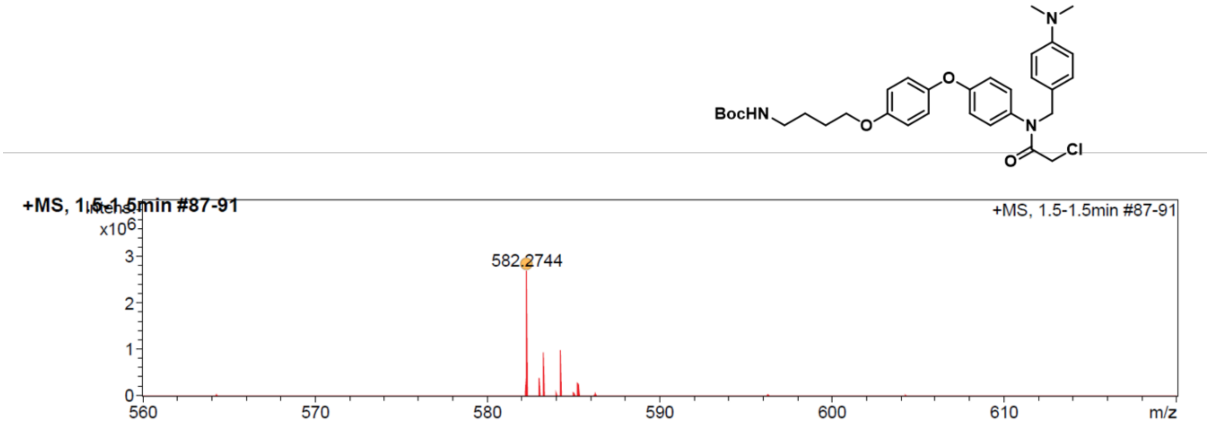


**[1]H NMR of R4VPL3:**

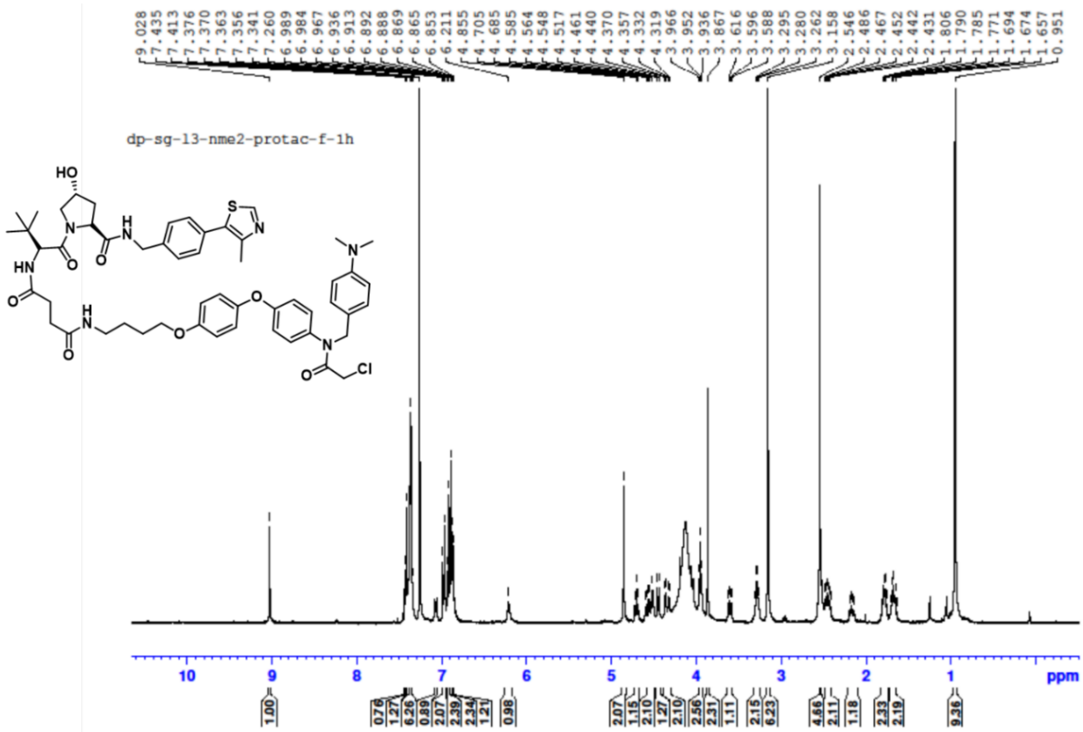

##### $^{13}C$ NMR of R4VPL3:

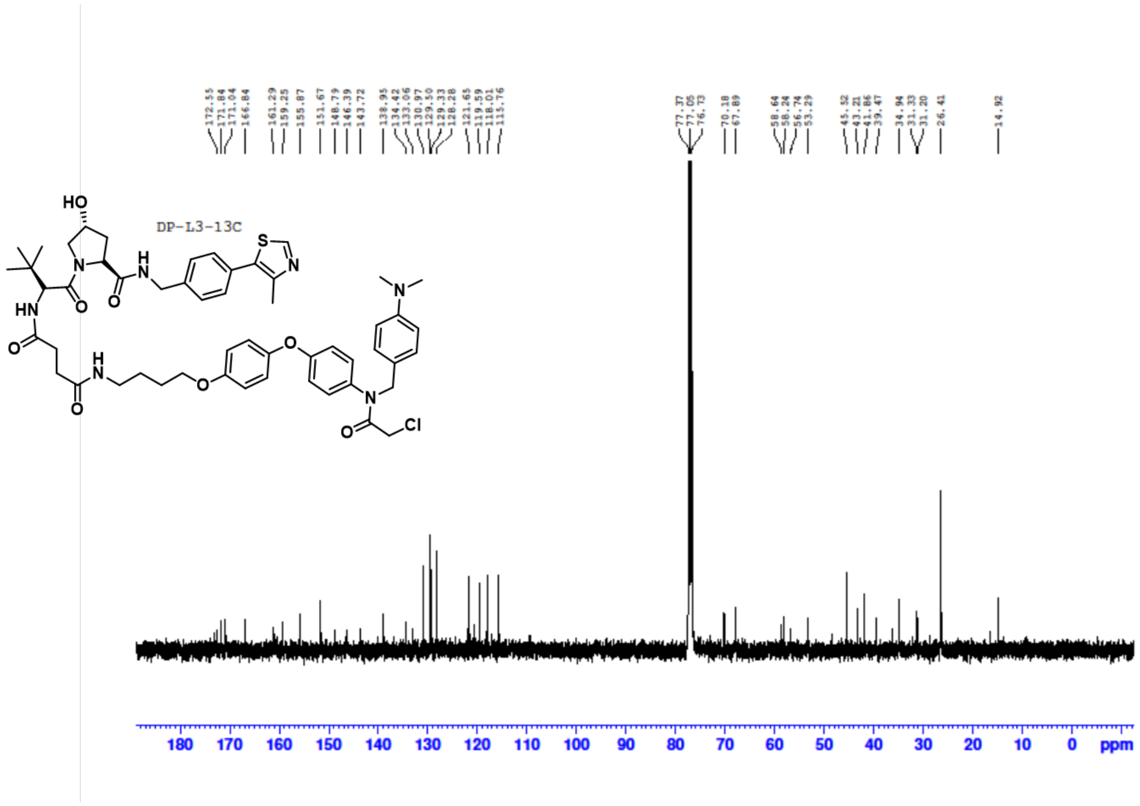

##### HRMs of R4VPL3:

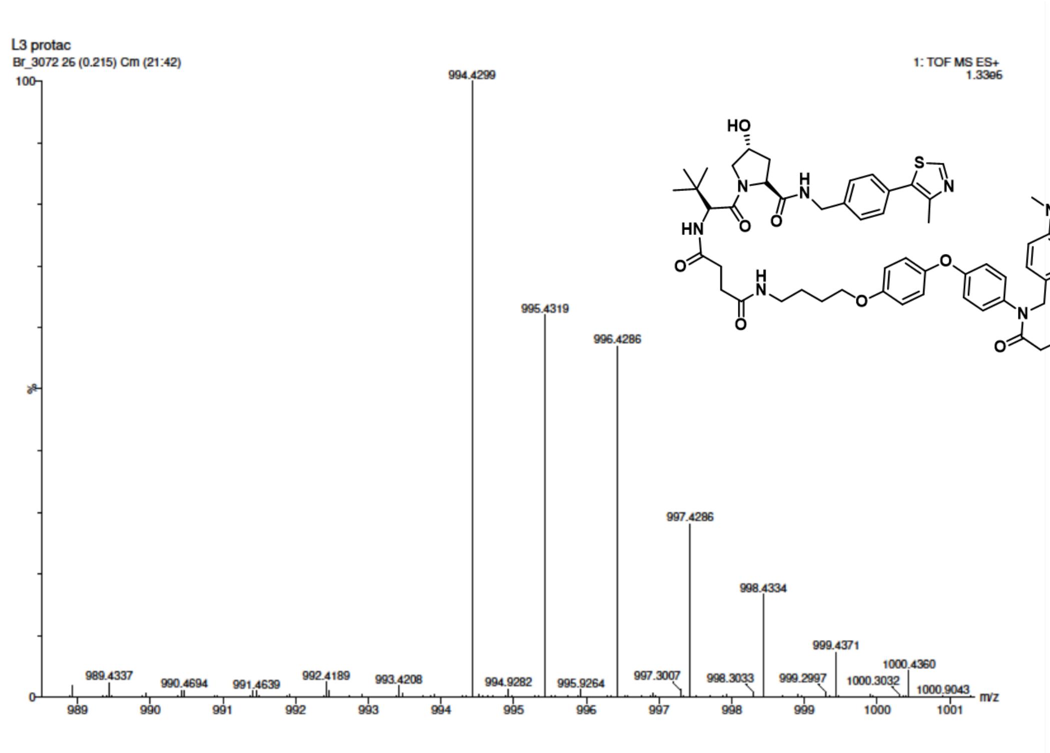

**$^{1}$H NMR of R4VP L3-1:**

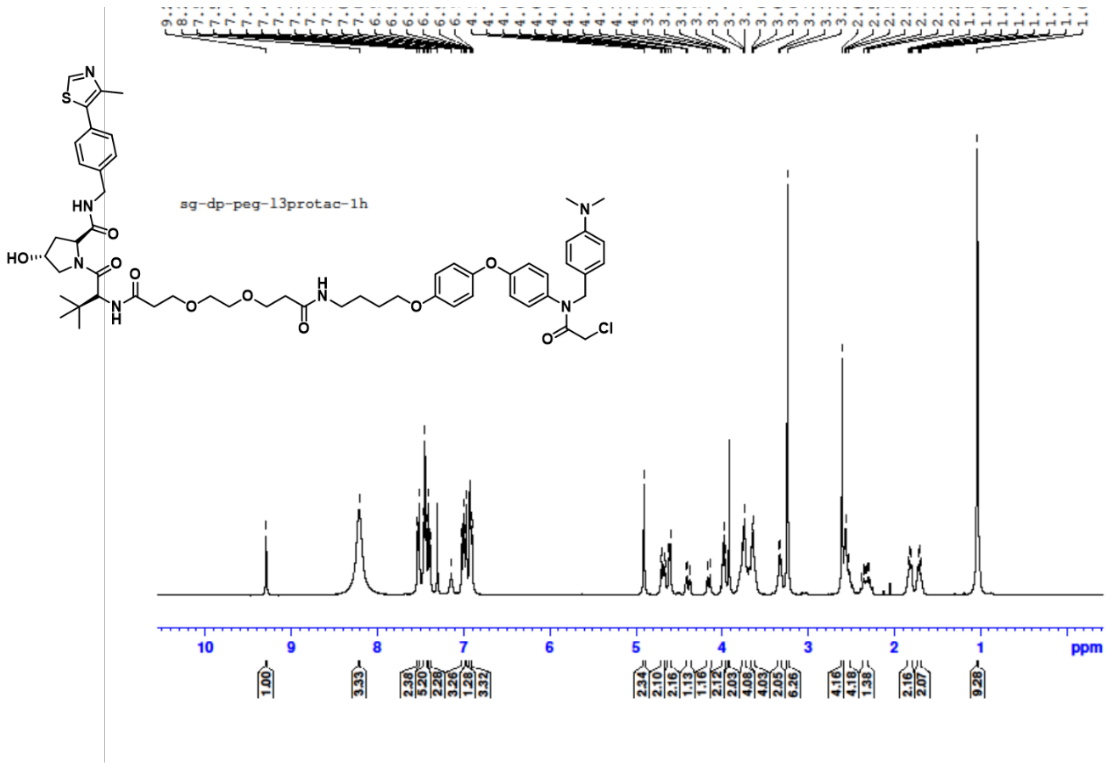


**$^{13}$C NMR of R4VP L3-1:**

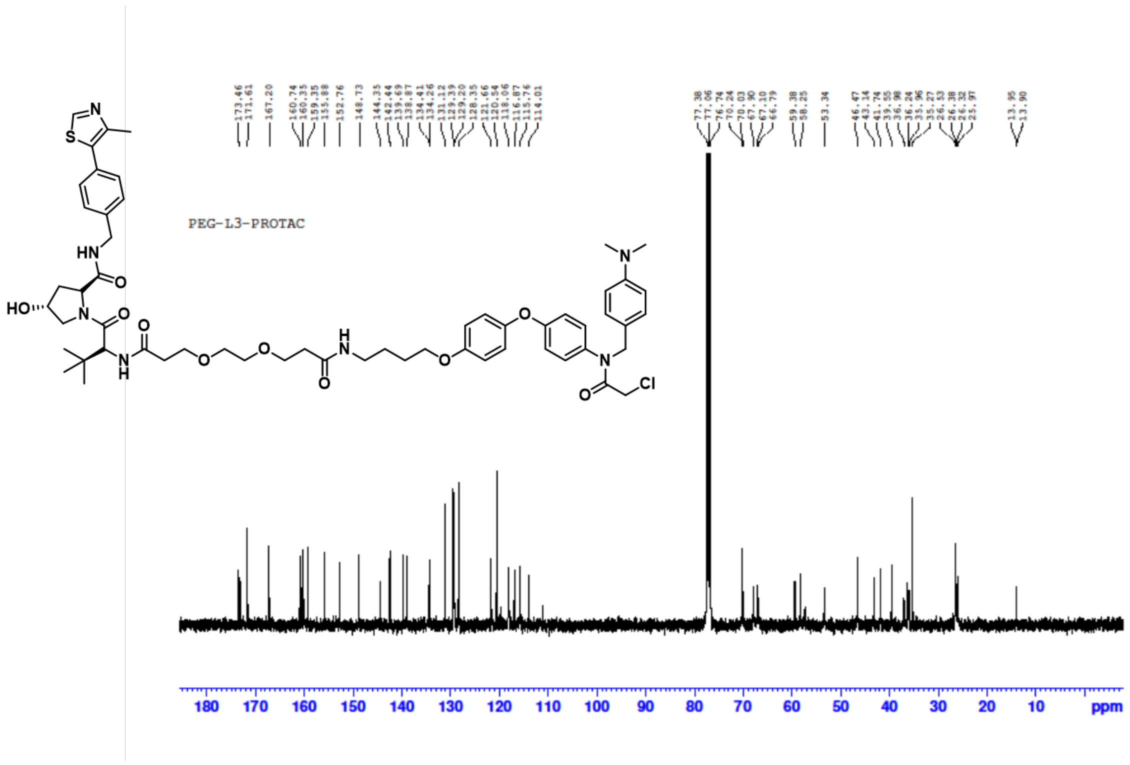

**HRMS of R4VP L3-1:**

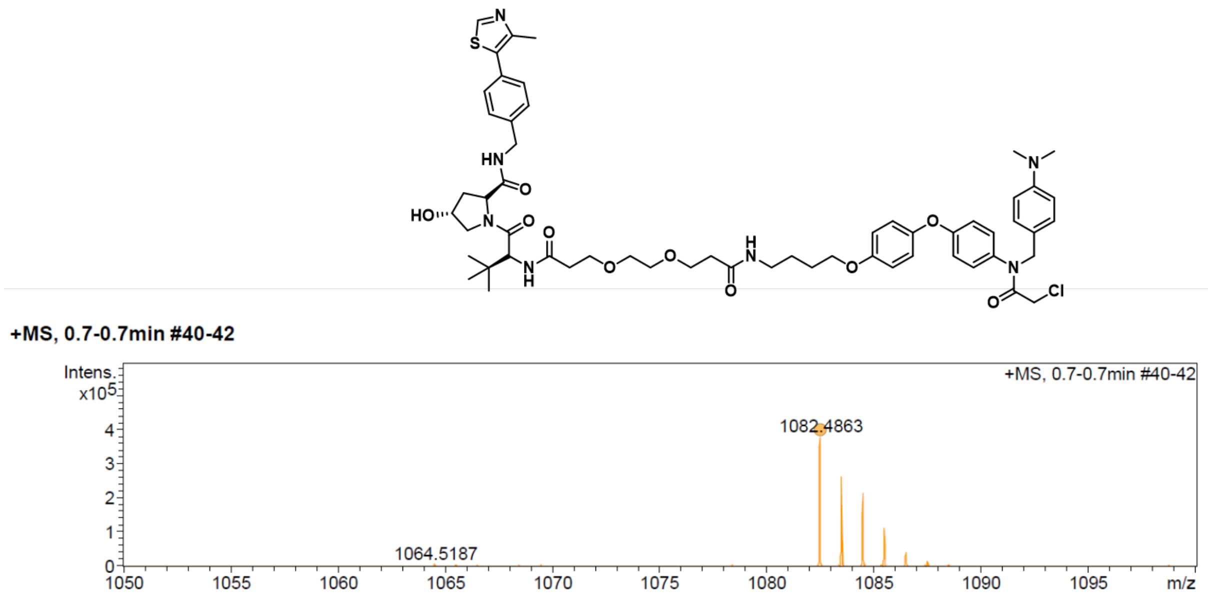


**[1]H NMR of compound 7:**

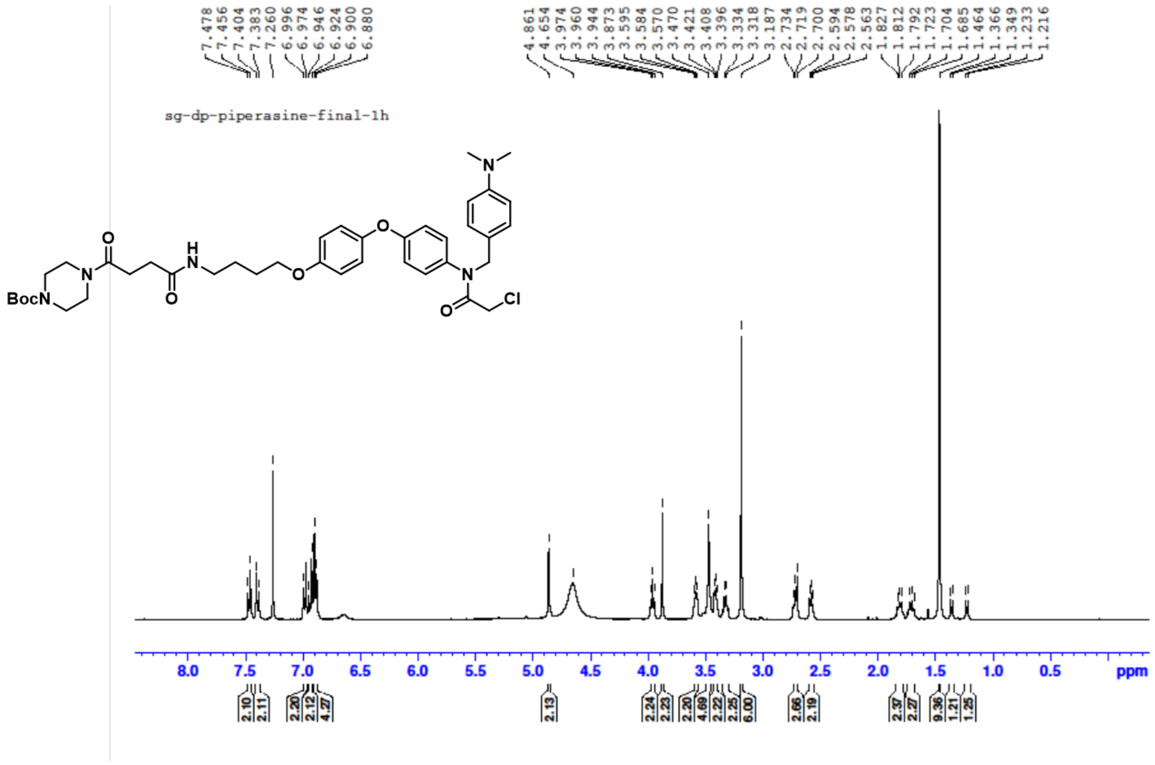

**$^{13}$C NMR of compound 7:**

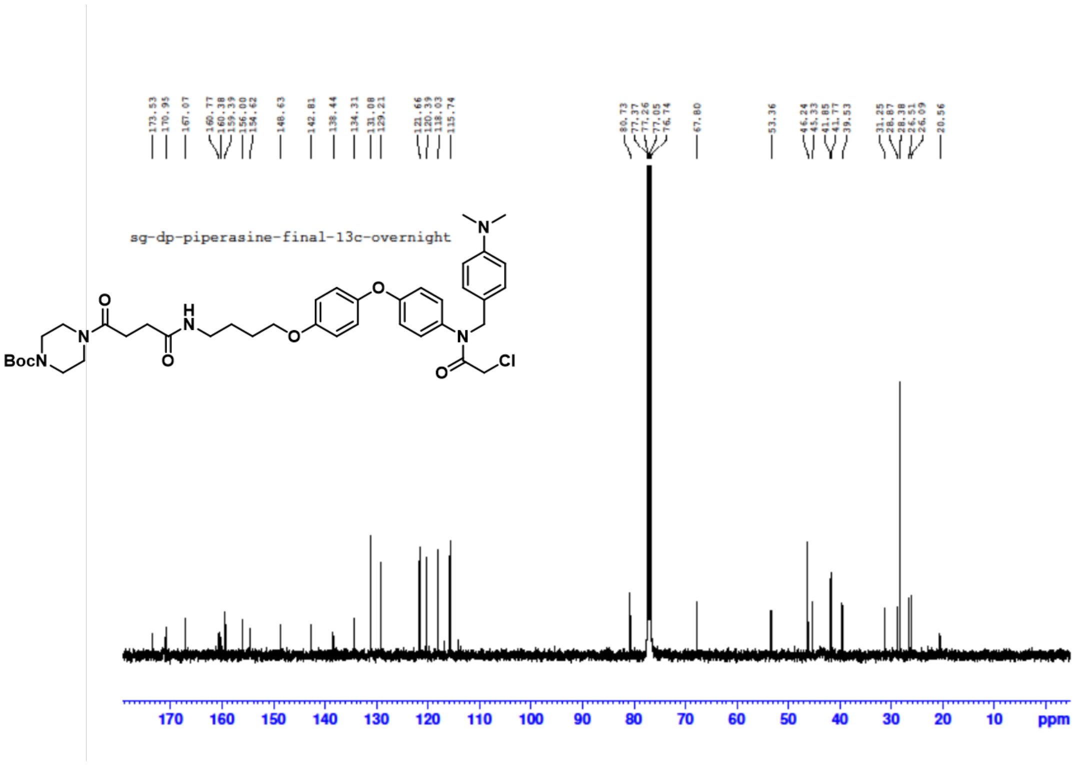


**HRMS of compound 7:**

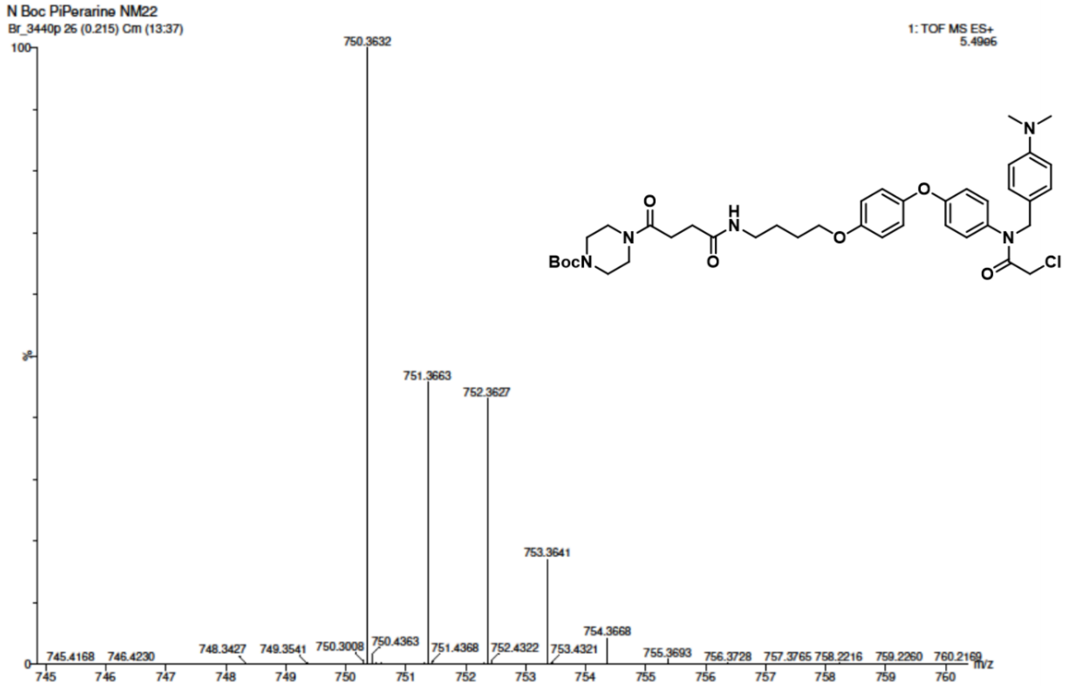

[1]H NMR of R4VPL3-2:

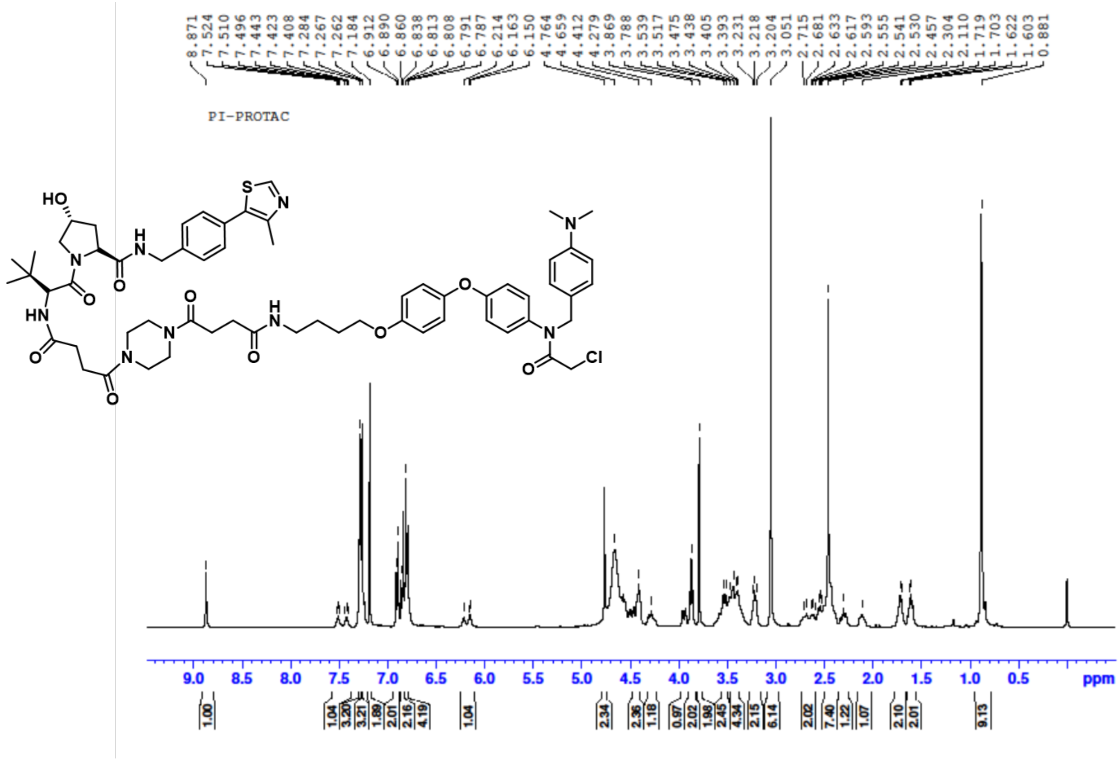

[13]C NMR of R4VPL3-2:

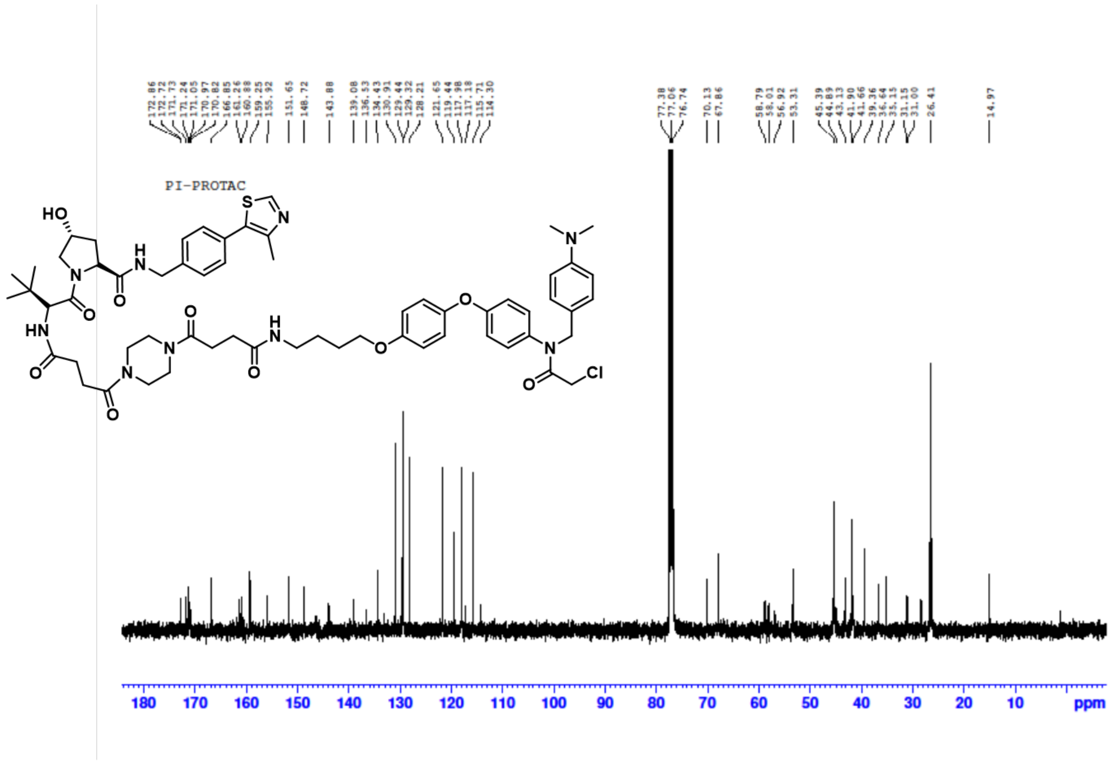

**HRMS of R4VPL3-3:**

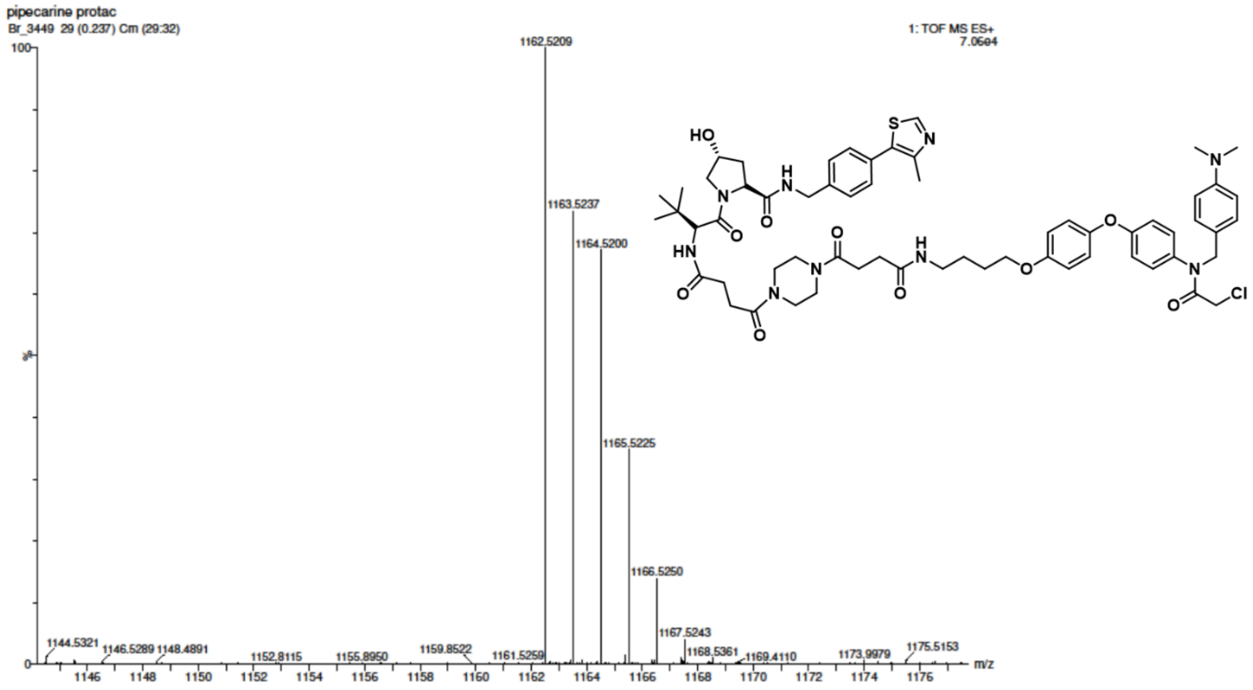


**[1]H NMR of R4VPL3-3:**

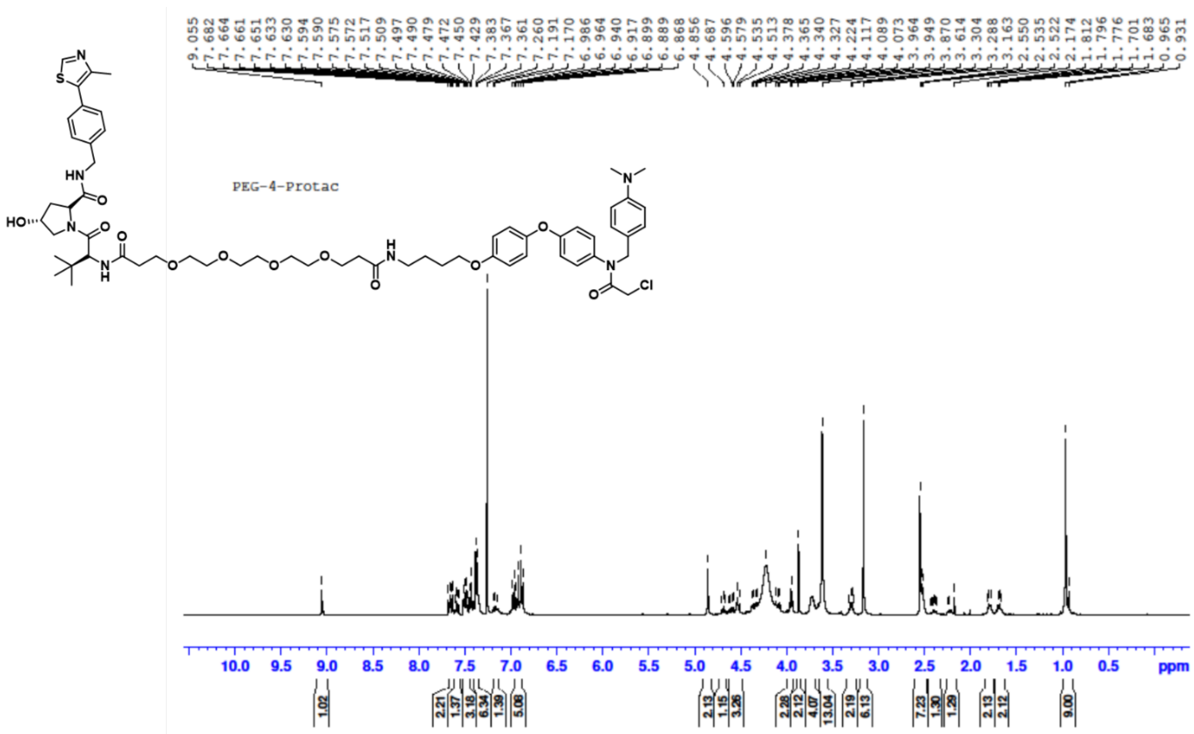

#### 13C NMR of R4VPL3-3:

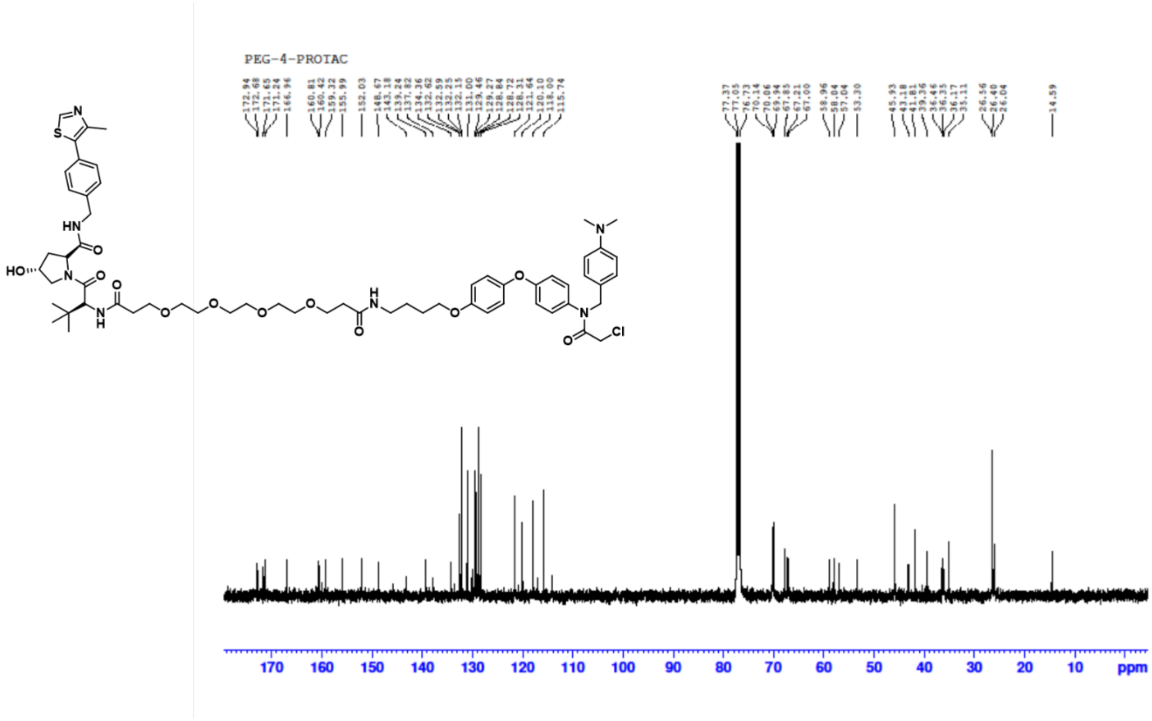


#### HRMS of R4VPL3-3:

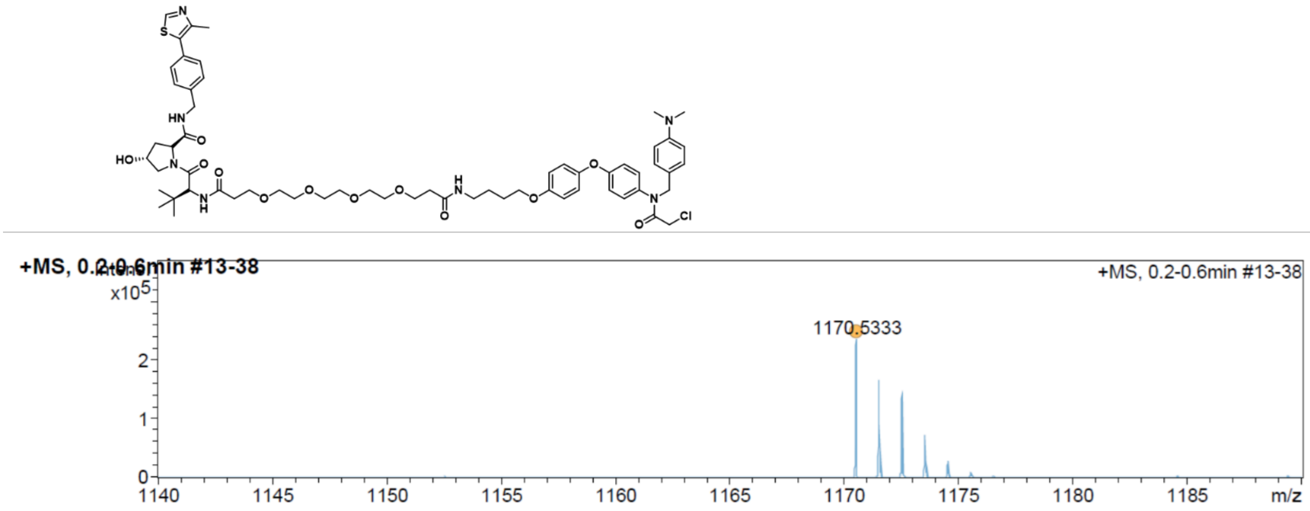

[1]H NMR of Peg-2-Control PROTAC-like:

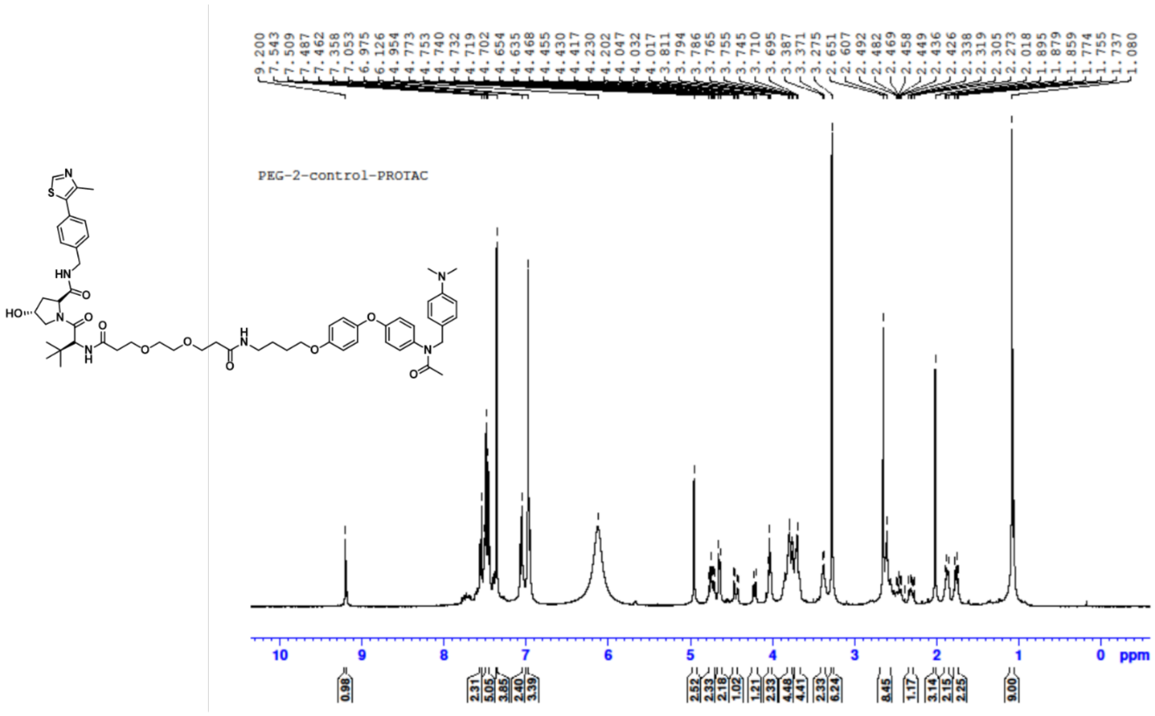

[13]C NMR of Peg-2-Control PROTAC-Like:

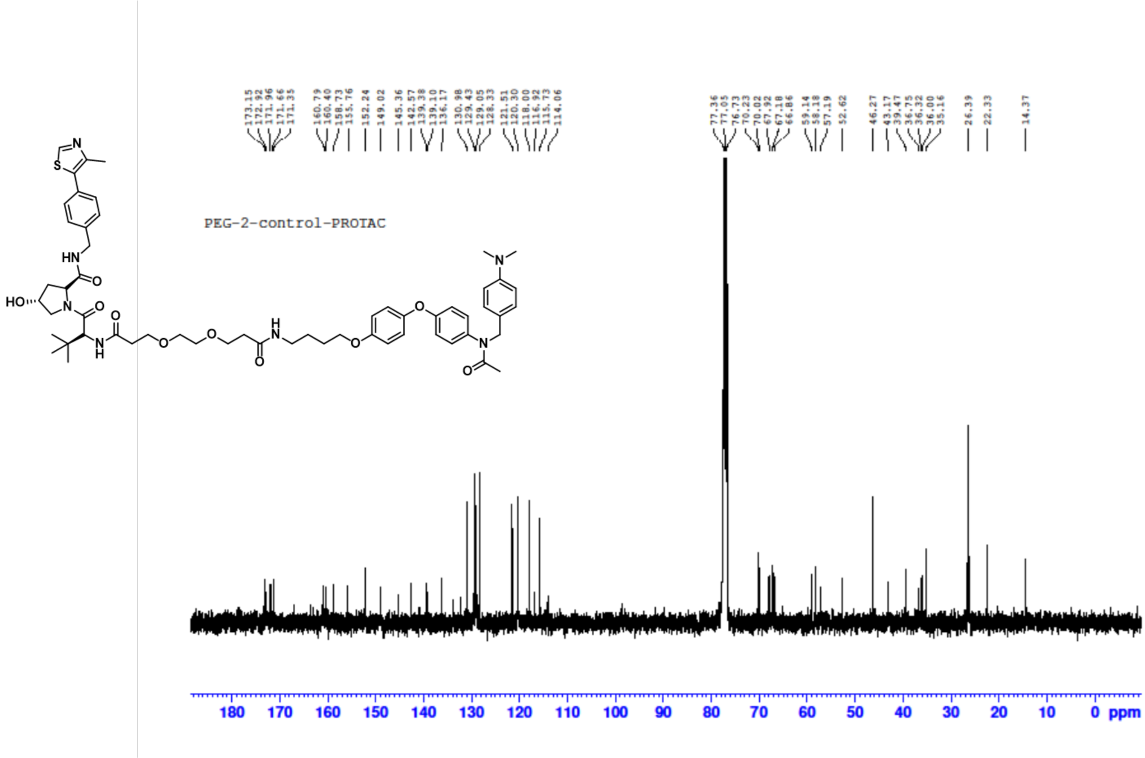

#### [1]H NMR of Biotin-R4VPL3-1:

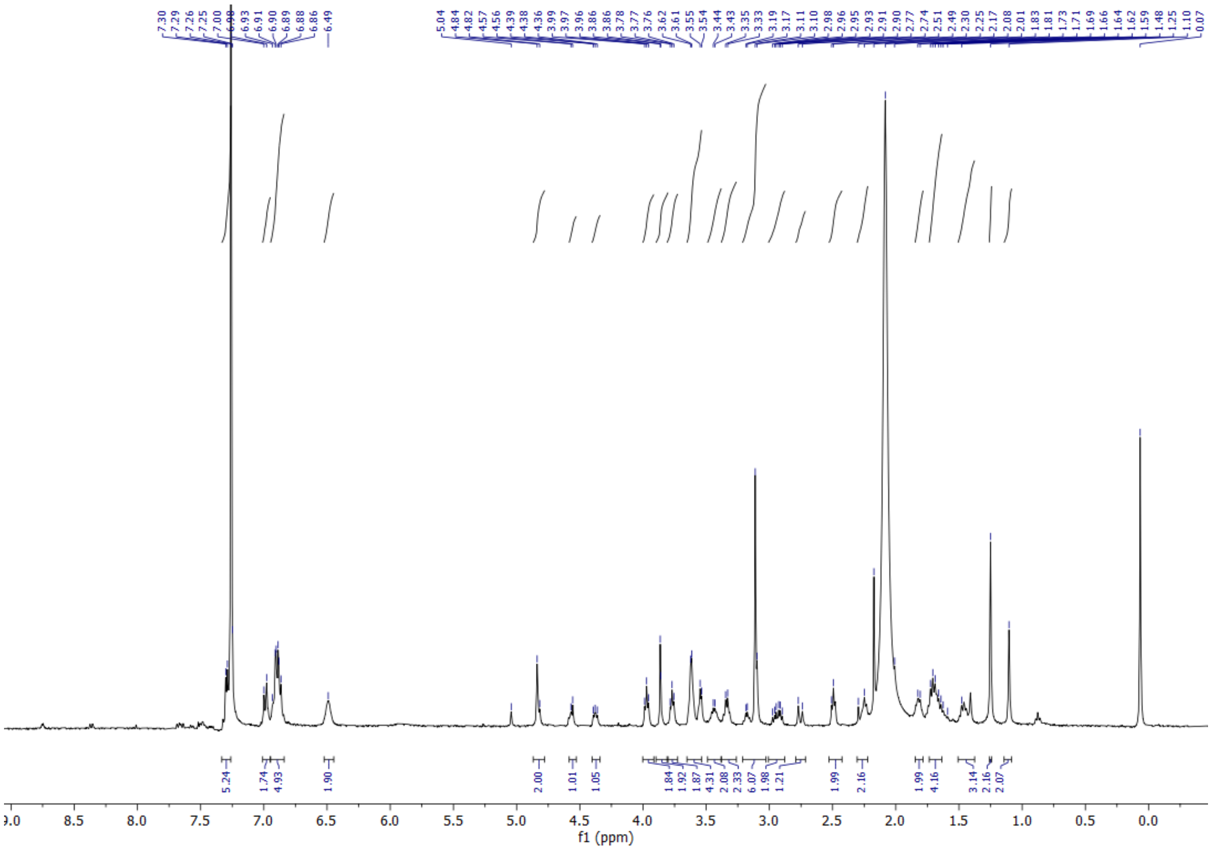


#### HRMS of Biotin-R4VPL3-1:

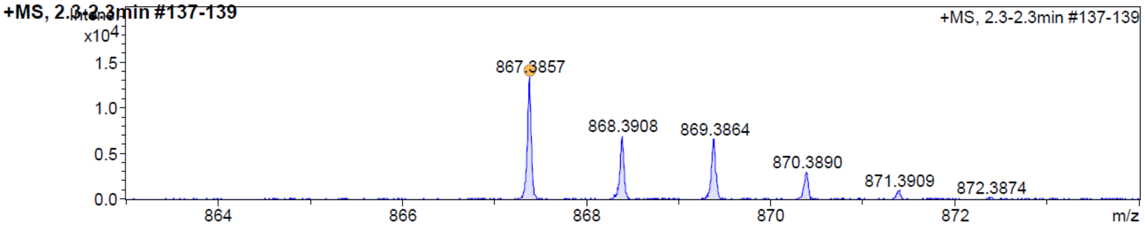


| Meas. m/z | # | Ion Formula | m/z | err [ppm] | mSigma | # mSigma | Score | rdb | e⁻ Conf | N-Rule | \|err\| [mDa] |
|---|---|---|---|---|---|---|---|---|---|---|---|
| 867.3857 | 1 | C44H60ClN6O8S | 867.3876 | 2.2 | 7.7 | 1 | 100.00 | 17.5 | even | ok | 1.9 |
